# Reproducible, portable, and efficient ancient genome reconstruction with nf-core/eager

**DOI:** 10.1101/2020.06.11.145615

**Authors:** James A. Fellows Yates, Thiseas C. Lamnidis, Maxime Borry, Aida Andrades Valtueña, Zandra Fagernäs, Stephen Clayton, Maxime U. Garcia, Judith Neukamm, Alexander Peltzer

## Abstract

The broadening utilisation of ancient DNA to address archaeological, palaeontological, and biological questions is resulting in a rising diversity in the size of laboratories and scale of analyses being performed. In the context of this heterogeneous landscape, we present nf-core/eager, an advanced and entirely redesigned and extended version of the EAGER pipeline for the analysis of ancient genomic data. This Nextflow pipeline aims to address three main themes: accessibility and adaptability to different computing configurations, reproducibility to ensure robust analytical standards, and updating the pipeline to the latest routine ancient genomic practises. This new version of EAGER has been developed within the nf-core initiative to ensure high-quality software development and maintenance support; contributing to a long-term lifecycle for the pipeline. nf-core/eager will assist in ensuring that ancient DNA sequencing data can be used by a diverse range of research groups and fields.

## Introduction

Ancient DNA (aDNA) has become a widely accepted source of biological data, helping to provide new perspectives for a range of fields including archaeology, cultural heritage, evolutionary biology, ecology, and palaeontology. The utilisation of short-read high-throughput sequencing has allowed the recovery of whole genomes and genome-wide data from a wide variety of sources, including (but not limited to), the skeletal remains of animals [1,2,3,4], modern and archaic humans [5,6,7,8], bacteria [9,10,11], viruses [12,13], plants [14,15], palaeofaeces [16,17], dental calculus [18,19], sediments [20,21], medical slides [22], parchment [23], and recently, ancient ‘chewing gum’ [24,25]. Improvement in laboratory protocols to increase yields of otherwise trace amounts of DNA has at the same time led to studies that can total hundreds of ancient individuals [26,27], spanning single [28] to thousands of organisms [18]. These differences of disciplines have led to a heterogeneous landscape in terms of the types of analyses undertaken, and their computational resource requirements [29,30]. Taking into consideration the unequal distribution of resources (and infrastructure such as internet connection), easy-to-deploy, streamlined and efficient pipelines can help increase accessibility to high-quality analyses.

The degraded nature of aDNA poses an extra layer of complexity to standard modern genomic analysis. Through a variety of processes [31] DNA molecules fragment over time, resulting in ultra-short molecules [32]. These sequences have low nucleotide complexity making it difficult to identify with precision which part of the genome a read (a sequenced DNA molecule) is derived from. Fragmentation without a ‘clean break’ leads to uneven ends, consisting of single-stranded ‘overhangs’ at end of molecules, which are susceptible to chemical processes such as deamination of nucleotides. These damaged nucleotides then lead to misincorporation of complementary bases during library construction for high-throughput DNA sequencing [33]. On top of this, taphonomic processes such as heat, moisture, and microbial- and burial-environment processes lead to varying rates of degradation [34,35]. The original DNA content of a sample is therefore increasingly lost over time and supplanted by younger ‘environmental’ DNA. Later handling by archaeologists, museum curators, and other researchers can also contribute ‘modern’ contamination. While these characteristics can help provide evidence towards the ‘authenticity’ of true aDNA sequences (e.g. the aDNA cytosine to thymine or C to T ‘damage’ deamination profiles [36]), they also pose specific challenges for genome reconstruction, such as unspecific DNA alignment and/or low coverage and miscoding lesions that can result in low-confidence genotyping. These factors often lead to prohibitive sequencing costs when retrieving enough data for modern high-throughput short-read sequencing data pipelines (such as more than 1 billion reads for a 1X depth coverage *Yersinia pestis* genome [37]), and thus aDNA-tailored methods and techniques are required to overcome these challenges.

Two previously published and commonly used pipelines in the field are PALEOMIX [38] and EAGER [39]. These two pipelines take a similar approach to link together standard tools used for Illumina high-throughput short-read data processing (sequencing quality control, sequencing adapter removal/and or paired-end read merging, mapping of reads to a reference genome, genotyping, etc.). However, they have a specific focus on tools that are designed for, or well-suited for aDNA (such as the bwa aln algorithm for ultra-short molecules [40] and mapDamage [41] for evaluation of aDNA characteristics). Yet, neither of these genome reconstruction pipelines have had major updates to bring them in-line with current routine aDNA analyses. *Meta*genomic screening of off-target genomic reads for pathogens or microbiomes [18,19] has become particularly common in palaeo- and archaeogenetics, given its role in revealing widespread infectious disease and possible epidemics that have sometimes been previously undetected in the archaeological record [12,13,37,42]. Without easy access to the latest field-established analytical routines, ancient genomic studies risk being published without the necessary quality control checks that ensure aDNA authenticity, as well as limiting the full range of possibilities from their data. Given that material from samples is limited, there are both ethical as well as economical interests to maximise analytical yield [43].

To address these shortcomings, we have completely re-implemented the latest version of the EAGER pipeline in Nextflow [44] (a domain-specific-language or ‘DSL’, specifically designed for the construction of omics analysis pipelines), introduced new features, and more flexible pipeline configuration. In addition, the renamed pipeline - nf-core/eager - has been developed in the context of the nf-core community framework [45], which enforces strict guidelines for best-practices in software development.

## Results and Discussion

### Scalability, Portability, and Eficiency

The re-implementation of EAGER into Nextflow offers a range of benefits over the original custom pipeline framework.

Firstly, the new framework provides immediate integration of nf-core/eager into various job schedulers in POSIX High-Performance-Cluster (HPC) environments, cloud computing resources, as well as local workstations. This portability allows users to set up nf-core/eager regardless of the type of computing infrastructure or cluster size (if applicable), with minimal effort or configuration. This facilitates reproducibility and therefore maintenance of standards within the field. Portability is further assisted by the in-built compatibility with software environments and containers such as Conda [46], Docker [47] and Singularity [48]. These are isolated software ‘sandbox’ environments that include all software (with exact versions) required by the pipeline, in a form that is installable and runnable by users regardless of the set up of their local software environment. Another major change with nf-core/eager is that the primary user interaction mode of a pipeline run set up is now with a command-line interface (CLI), replacing the graphical-user-interface (GUI) of the original EAGER pipeline. This is more portable and compatible with most HPCs (that may not offer display of a window system), and is in line with the vast majority of bioinformatics tools. We therefore believe this will not be a hindrance to new researchers from outside computational biology. However, a GUI-based pipeline set up is still avaliable via the nf-core website’s Launch page [49], which provides a common GUI format across multiple pipelines, as well as additional robustness checks of input parameters for those less familiar with CLIs. Typically the output of the launch functionality is a JSON file that can be used with a nf-core/tools launch command as a single parameter (similar to the original EAGER), however integration with Nextflow’s companion monitoring tool tower.nf [50] also allows direct submission of pipelines without any command line usage.

Secondly, reproducibility is made easier through the use of ‘profiles’ that can define configuration parameters. These profiles can be managed at different hierarchical levels. *HPC-level profiles* can specify parameters for the computing environment (job schedulers, cache locations for containers, maximum memory and CPU resources etc.), which can be centrally managed to ensure all users of a group use the same settings. *Pipeline-level profiles*, specifying parameters for nf-core/eager itself, allow fast access to routinely-run pipeline parameters via a single flag in the nf-core/eager run command, without having to configure each new run from scratch. Compared to the original EAGER, which utilised per-FASTQ XML files with hardcoded filepaths for a specific user’s server, nf-core/eager allows researchers to publish the specific profile used in their runs alongside their publications, that can also be used by other groups to generate the same results. Usage of profiles can also reduce mistakes caused by insufficient ‘prose’ based reporting of program settings that can be regularly found in the literature. The default nf-core/eager profile uses parameters evaluated in different aDNA-specific contexts (e.g. in [51]), and will be updated in each new release as new studies are published.

Finally, nf-core/eager provides improved efficiency over the original EAGER pipeline by replacing sample-by-sample sequential processing with Nextflow’s asynchronous job parallelisation, whereby multiple pipeline steps and samples are run in parallel (in addition to natively parallelised pipeline steps). This is similar to the approach taken by PALEOMIX, however nf-core/eager expands this by utilising Nextflow’s ability to customise the resource parameters for every job in the pipeline; reducing unnecessary resource allocation that can occur with unfamiliar users to each step of a high-throughput short-read data processing pipeline. This is particularly pertinent given the increasing use of centralised HPCs or cloud computing that often use per-hour cost calculations.

### Updated Worklow

nf-core/eager follows a similar structural foundation to the original version of EAGER and partially to PALEOMIX. Given Illumina short-read FASTQ and/or BAM files and a reference FASTA file, the core functionality of nf-core/eager can be split in five main stages:

1. Pre-processing
  - Sequencing quality control: FastQC [52]
  - Sequencing artefact clean-up (merging, adapter clipping): AdapterRemoval2 [53], fastp [54]
  - Pre-processing statistics generation: FastQC
2. Mapping and post-processing
  - Alignment against reference genome: BWA aln and mem [40,55], CircularMapper [39], Bowtie2 [57]
  - Mapping quality filtering: SAMtools [58]
  - PCR duplicate removal: DeDup [39], Picard MarkDuplicates [59]
  - Mapping statistics generation: SAMTools, PreSeq [60], Qualimap2 [61], bedtools [62], Sex.DetERRmine [63]
3. aDNA evaluation and modification
  - Damage profiling: DamageProfiler [64]
  - aDNA reads selection: PMDtools [65]
  - Damage removal/Base trimming: Bamutils [66]
  - Human nuclear contamination estimation: ANGSD [67]
4. Variant calling and consensus sequence generation: GATK UnifiedGenotyper and HaploTypeCaller [59], ANGSD [67], sequenceTools pileupCaller [68], VCF2Genome [39], MultiVCFAnalyzer [9]
5. Report generation: MultiQC [69]

In nf-core/eager, all tools originally used in EAGER have been updated to their latest versions, as available on Bioconda [70] and Conda-forge [71], to ensure widespread accessibility and stability of utilised tools. The mapDamage2 (for damage profile generation) [36] and Schmutzi (for mitochondrial contamination estimation) [72] methods have not been carried over to nf-core/eager, the first because a more performant successor method is now available (DamageProfiler), and the latter because a stable release of the method could not be migrated to Bioconda. We anticipate that there will be an updated version of Schmutzi in the near future that will allow us to integrate the method again into nf-core/eager. As an alternative, estimation of human *nuclear* contamination is now offered through ANGSD [67]. Support for the Bowtie2 aligner [57] has been updated to have default settings optimised for aDNA [73].

New tools to the basic workflow include fastp [54] for the removal of ‘poly-G’ sequencing artefacts that are common in 2-colour Illumina sequencing machines (such as the increasingly popular NextSeq and NovaSeq platforms [74]). For variant calling, we have now included FreeBayes [75] as an alternative to the human-focused GATK tools, and have also added pileupCaller [68] for generation of genotyping formats commonly utilised in ancient human population analysis. We have also maintained the possibility of using the now unsupported GATK UnifiedGenotyper, as the supported replacement, GATK HaplotypeCaller, performs *de novo* assembly around possible variants; something that may not be suitable for low-coverage aDNA data.

Additional functionality tailored for ancient bacterial genomics includes integration of a SNP alignment generation tool, MultiVCFAnalyzer [9], which includes the ability to make an assessment of levels of cross-mapping from different related taxa to a reference genome - a common challenge in ancient bacterial genome reconstruction [35]. The output SNP consensus alignment FASTA file can then be used for downstream analyses such as phylogenetic tree construction. Simple coverage statistics of particular annotations (e.g. genes) of an input reference is offered by bedtools [62], which can be used in cases such as for providing initial indications of functional differences between ancient bacterial strains (as in [42]). When using a human reference genome, nf-core/eager can also give estimates of the relative coverage on the X and Y chromosomes with Sex.DetERRmine that can be used to infer the biological sex of a given human individual [63]. A dedicated ‘endogenous DNA’ calculator (endorS.py) is also included, to provide a percentage estimate of the sequenced reads matching the reference (‘on-target’) from the total number of reads sequenced per library.

Given the large amount of sequencing often required to yield sufficient genome coverage from aDNA data, palaeogeneticists tend to use multiple (differently treated) libraries, and/or merge data from multiple sequencing runs of each library or even samples. The original EAGER pipeline could only run a single library at a time, and in these contexts required significant manual user input in merging different FASTQ or BAM files of related libraries. A major upgrade in nf-core/eager is that the new pipeline supports automated processing of complex sequencing strategies for many samples, similar to PALEOMIX. This is facilitated by the optional use of a simple table (in TSV format, a format more commonly used in wet-lab stages of data generation, compared to PALEOMIX’s YAML format) that includes file paths and additional metadata such as sample name, library name, sequencing lane, colour chemistry, and UDG treatment. This allows automated and simultaneous processing and appropriate merging and treatment of heterogeneous data from multiple sequencing runs and/or library types.

The original EAGER and PALEOMIX pipelines required users to look through many independent output directories and files to make full assessment of their sequencing data. This has now been replaced in nf-core/eager with a much more extensive MultiQC report [69]. This tool aggregates the log files of every supported tool into a single interactive report, and assists users in making a fuller assessment of their sequencing and analysis runs. We have developed a corresponding MultiQC module for every tool used by nf-core/eager, where possible, to enable comprehensive evaluation of all stages of the pipeline.

We have further extended the functionality of the original EAGER pipeline by adding ancient metagenomic analysis; allowing reconstruction of the wider taxonomic content of a sample. We have added the possibility to screen all off-target reads (not mapped to the reference genome) with two metagenomic profilers: MALT [76,77] and Kraken2 [78], in parallel to the mapping to a given reference genome (typically of the host individual, assuming the sample is a host organism). Characterisation of properties of authentic aDNA from metagenomic MALT alignments is carried out with MaltExtract of the HOPS pipeline [79]. This functionality can be used either for microbiome screening or putative pathogen detection. Ancient metagenomic studies sometimes include comparative samples from living individuals [80]. To support open data, whilst respecting personal data privacy, nf-core/eager includes a ‘FASTQ host removal’ script that creates raw FASTQ files, but with all reads successfully mapped to the reference genome removed. This allows for safe upload of metagenomic non-host sequencing data to public repositories after removal of identifiable (human) data, for example for microbiome studies.

An overview of the entire pipeline is shown in Figure 1, and a tabular comparison of functionality between EAGER, PALEOMIX and nf-core/eager is in Table 1.

**Table 1:** Comparison of pipeline functionality of common ancient DNA processing pipelines. Tick represents full functionality, tilde represents partial functionality, and cross represents not implemented.

| Category | Functionality | EAGER | PALEOMIX | nf-core/eager |
| --- | --- | --- | --- | --- |
| Infrastructure | Reproducible software environments offered | ✓ | ✗ | ✓ |
|  | HPC scheduler integration | ✗ | ✗ | ✓ |
|  | Cloud computing integration | ✗ | ✗ | ✓ |
|  | Per-process resource optimisation | ✗ | ~ | ✓ |
|  | Pipeline-step parallelisation | ✗ | ✓ | ✓ |
|  | Command line set up | ✗ | ✓ | ✓ |
|  | GUI set up | ✓ | ✗ | ✓ |
| Preprocessing | Sequencing lane merging | ✓ | ✓ | ✓ |
|  | Sequencing quality control | ✓ | ✗ | ✓ |
|  | Sequencing artefact removal | ✗ | ✗ | ✓ |
|  | Adapter clipping/read merging | ✓ | ✓ | ✓ |
|  | Post-processing sequencing QC | ✗ | ✗ | ✓ |
| Alignment | Reference mapping | ✓ | ✓ | ✓ |
|  | Reference mapping statistics | ✓ | ✓ | ✓ |
|  | Multi-reference mapping | ✗ | ✓ | ✗ |
| Postprocessing | Mapped reads filtering | ✓ | ✓ | ✓ |
|  | Off-target metagenomic profiling | ✗ | ✗ | ✓ |
|  | Off-target metagenomic authentication | ✗ | ✗ | ✓ |
|  | Library complexity estimation | ✓ | ✗ | ✓ |
|  | Duplicate removal | ✓ | ✗ | ✓ |
|  | BAM merging | ✗ | ✓ | ✓ |
| Authentication | Damage read filtering | ✓ | ✗ | ✓ |
|  | Contamination estimation (Human) | ✓ | ✗ | ✓ |
|  | Biological sex determination (Human) | ✗ | ✗ | ✓ |
|  | Genome coverage estimation | ✓ | ✓ | ✓ |
|  | Damage calculation | ✓ | ✓ | ✓ |
|  | Damage rescaling | ✗ | ✓ | ~ |
| Downstream | SNP Calling/Genotyping | ✓ | ~ | ✓ |
|  | Consensus sequence generation | ✓ | ~ | ✓ |
|  | Regions of interest statistics | ~ | ✓ | ✓ |

**Figure 1:**
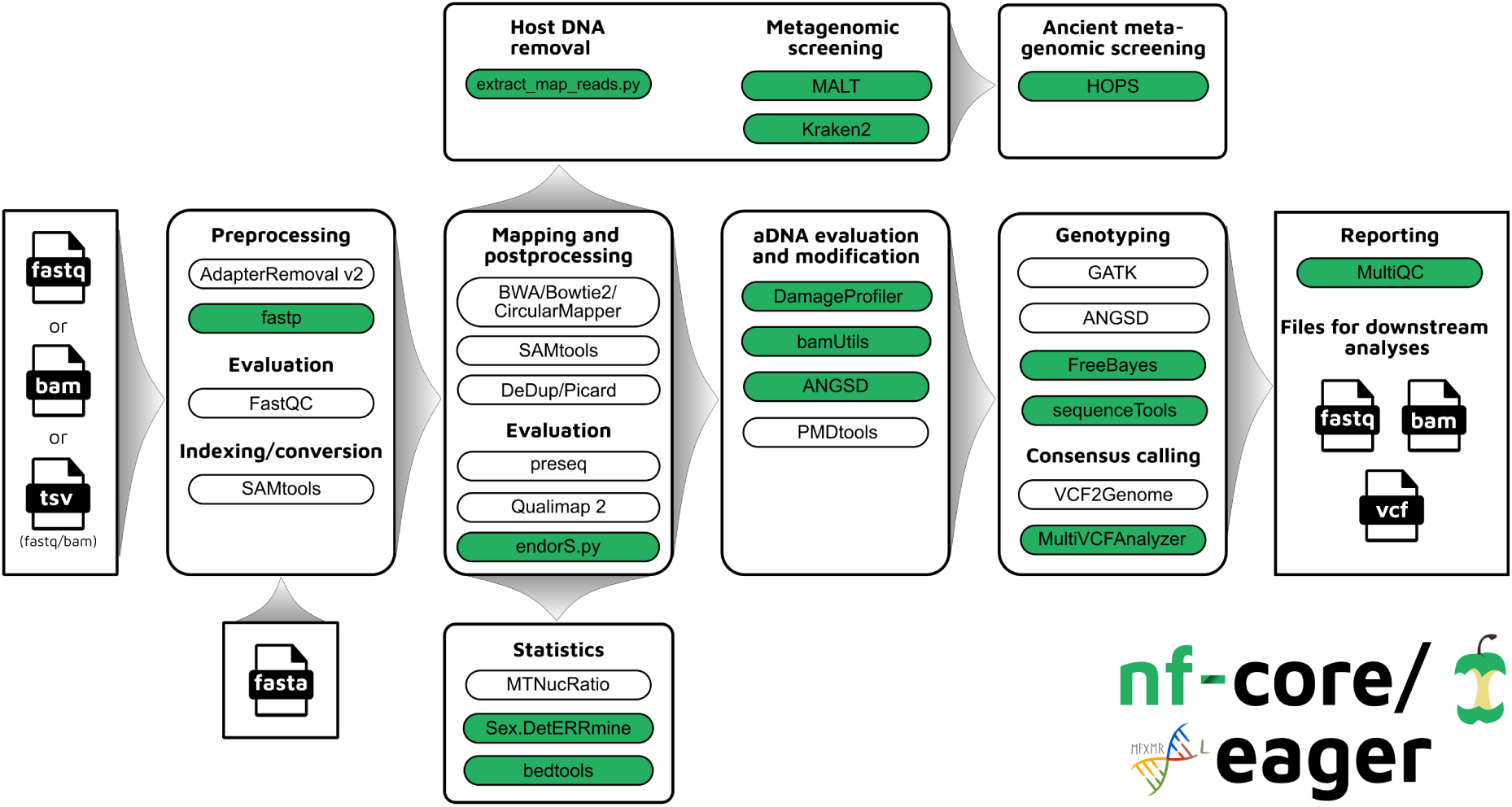
Simplified schematic of the nf-core/eager workflow pipeline. Green filled bubbles indicate new functionality added over the original EAGER pipeline.

To demonstrate the simultaneous genomic analysis of human DNA and metagenomic screening for putative pathogens, as well as improved results reporting, we re-analysed data from Barquera et al. 2020 [81], who performed a multi-discipline study of three 16th century individuals excavated from a mass burial site in Mexico City. The authors reported genetic results showing sufficient on-target human DNA (>1%) with typical aDNA damage (>20% C to T reference mismatches in the first base of the 5’ ends of reads) for downstream population-genetic analysis and Y-chromosome coverage indicative that the three individuals were genetically male. In addition, one individual (Lab ID: SJN003) contained DNA suggesting a possible infection by *Treponema pallidum*, a species with a variety of strains that can cause diseases such as syphilis, bejel and yaws, and a second individual (Lab ID: SJN001) displayed reads similar to the Hepatitis B virus. Both results were confirmed by the authors via in-solution enrichment approaches.

We were able to successfully replicate the human and pathogen screening results in a single run of nf-core/eager. Mapping to the human reference genome (hs37d5) with BWA aln and binning of off-target reads with MALT to the NCBI Nucleotide database (2017-10-26), yielded the same results of all individuals having a biological sex of male, as well as the same frequency of C to T miscoding lesions and short fragment lengths (both characteristic of true aDNA). Metagenomic hits to both pathogens from the corresponding individuals that also yielded complete genomes in the original publication were also detected. Both results and other processing statistics were identified via a single interactive MultiQC report, excerpts of which can be seen in Figure 2. The full interactive report can be seen in the supplementary information.

**Figure 2:**
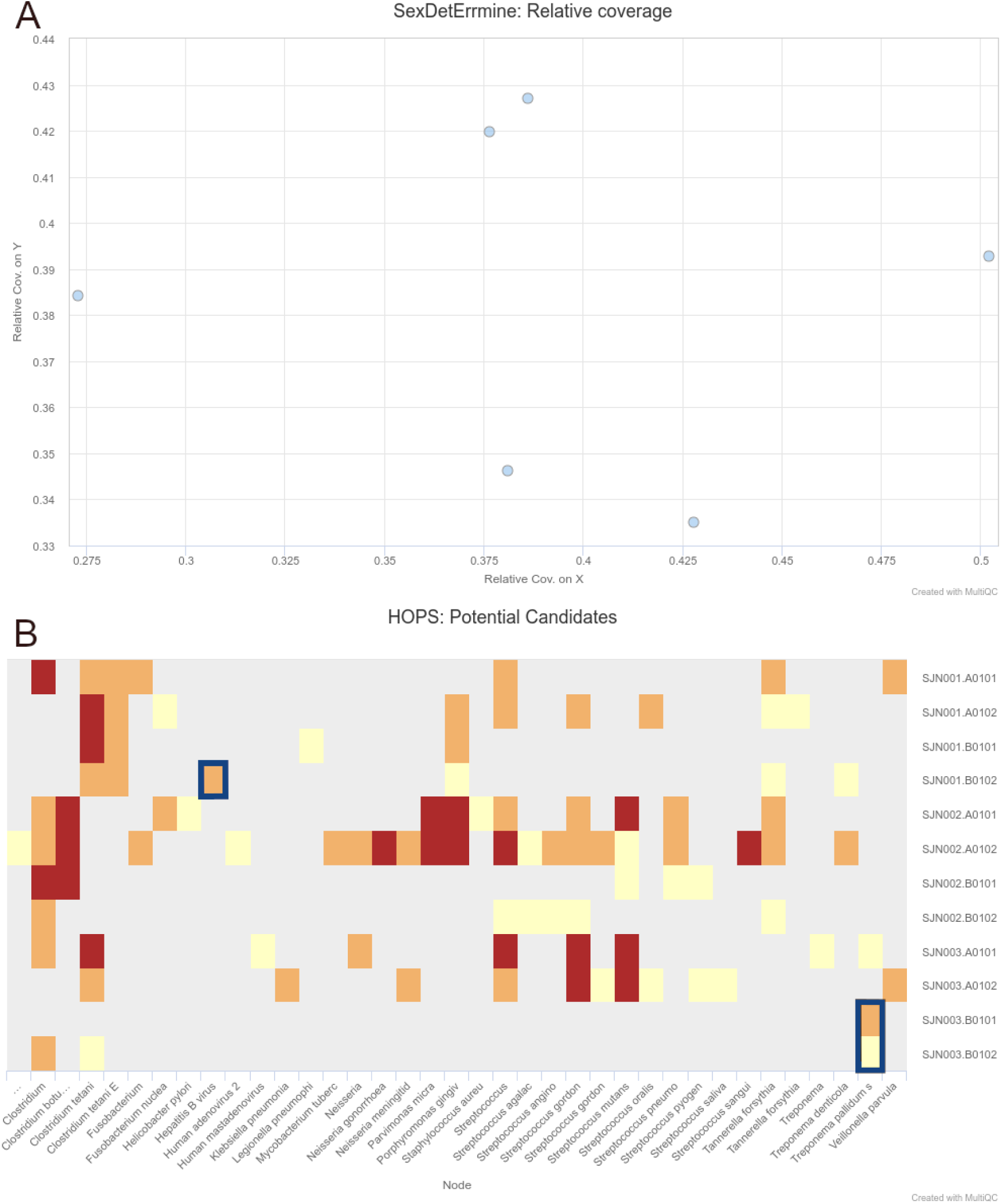
Sections of a MultiQC report (v1.10dev) with the outcome of simultaneous human DNA and microbial pathogen screening with nf-core/eager, including **A** Sex.DetERRmine output of biological sex assignment with coverages on X and Y being half of that of autosomes, indicative of male individuals, and **B** HOPS output with positive detection of both *Treponema pallidum* and Hepatitis B virus reads - indicated with blue boxes. Other taxa in HOPS output represent typical environmental contamination and oral commensal microbiota found in archaeological teeth. Data was Illumina shotgun sequencing data from Barquera et al. 2020 [81], and replicated results here were originally verified in the publication via enrichment methods. The full interactive reports for both MultiQC v1.9 and v1.10 (see methods) can be seen in the supplementary information.

### Accessibility

Alongside the interactive MultiQC report, we have written extensive documentation on all parts of running and interpreting the output of the pipeline. Given that a large fraction of aDNA researchers come from fields outside computational biology, and thus may have limited computational training, we have written documentation and tutorials [82] that also gives guidance on how to run the pipeline and interpret each section of the report in the context of high-throughput sequencing data, but with with a special focus on aDNA. This includes best practice or expected output schematic images that are published under CC-BY licenses to allow for use in other training material (an example can be seen in Figure 3). We hope this open-access resource will make the study of aDNA more accessible to researchers new to the field, by providing practical guidelines on how to evaluate characteristics and effects of aDNA on downstream analyses.

**Figure 3:**
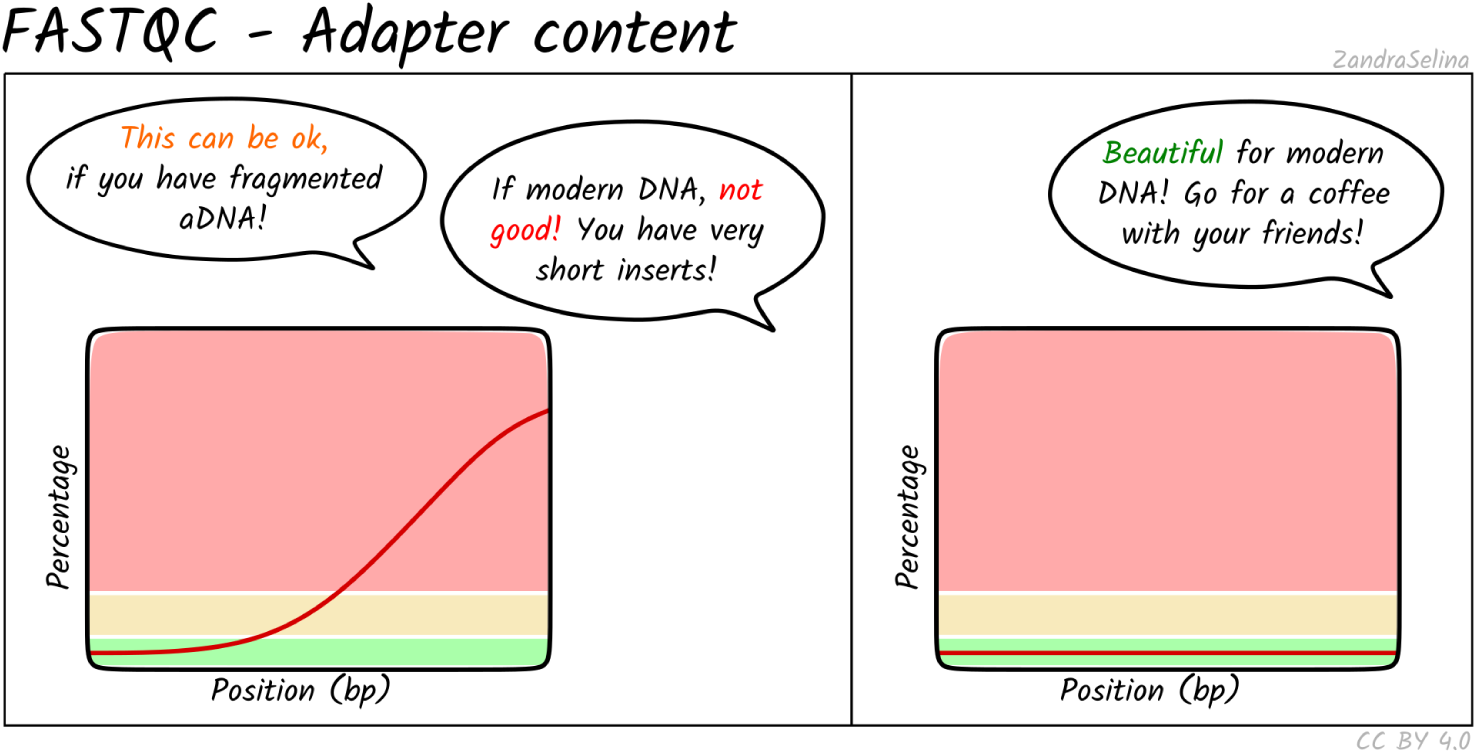
Example schematic images of pipeline output documentation that can assist new users in the interpretation of high-throughput sequencing aDNA processing.

The development of nf-core/eager in Nextflow and the nf-core initiative will also improve open-source development, while ensuring the high quality of community contributions to the pipeline. While Nextflow is written primarily in Groovy, the Nextflow DSL simplifies a number of concepts to an intermediate level that bioinformaticians without Java/Groovy experience can easily access (regardless of own programming language experience). Furthermore, Nextflow places ubiquitous and more widely known command-line interfaces, such as bash, in a prominent position within the code, rather than custom Java code and classes (as in EAGER). We hope this will motivate further bug fixes and feature contributions from the community, to keep the pipeline state-of-the-art and ensure a longer life-cycle. This will also be supported by the open and active nf-core community who provide general guidance and advice on developing Nextflow and nf-core pipelines.

### Comparisons with other pipelines

The scope of nf-core/eager is as a generic, initial data processing and screening tool, and not to act as a tool for performing more experimental analyses that requires extensive parameter testing such as modelling. As such, while similar pipelines designed for aDNA have also been released, for example ATLAS [83], these generally have been designed with specific contexts in mind (e.g. human population genetics). We therefore have opted to not include common downstream analysis such as Principal Component Analysis for population genetics, or phylogenetic analysis for microbial genomics, but rather focus on ensuring nf-core/eager produces useful files that can be easily used as input for common but more experimental and specialised downstream analysis.

Therefore, we compared pipeline run-times of two functionally equivalent and previously published pipelines to show that the new implementation of nf-core/eager is equivalent or more efficient than EAGER or PALEOMIX.

We ran each pipeline on a subset of Viking-age genomic data of cod (*Gadus morhua*) from Star et al. 2017 [4]. This data was originally run using PALEOMIX, and was re-run here as described, but with the latest version of PALEOMIX (v1.2.14), and with equivalent settings for the other two pipelines as close as possible to the original paper (EAGER with v1.92.33, and nf-core/EAGER with v2.2.0dev, commit 830c22d). The respective benchmarking environment and exact pipeline run settings can be seen in the Methods and Supplementary Information. Two samples each with three Illumina paired-end sequencing runs were analysed, with adapter clipping and merging (AdapterRemoval), mapping (BWA aln), duplicate removal (Picard’s MarkDuplicates) and damage profiling (PALEOMIX: mapDamage2, EAGER and nf-core/EAGER: DamageProfiler) steps being performed. We ran the commands for each tool sequentially, but repeated these batches of commands 10 times - to account for variability in the cloud service’s IO connection. Run times were measured using the GNU time tool (v1.7).

A summary of runtimes of the benchmarking tests can be seen in Table 2. nf-core/eager showed fastest runtimes across all three time metrics when running on default parameters. This highlights the improved efficiency of nf-core/eager’s asynchronous processing system and per-process resource customisation (here represented by nf-core/eager defaults designed for typical HPC set ups).

**Table 2:**
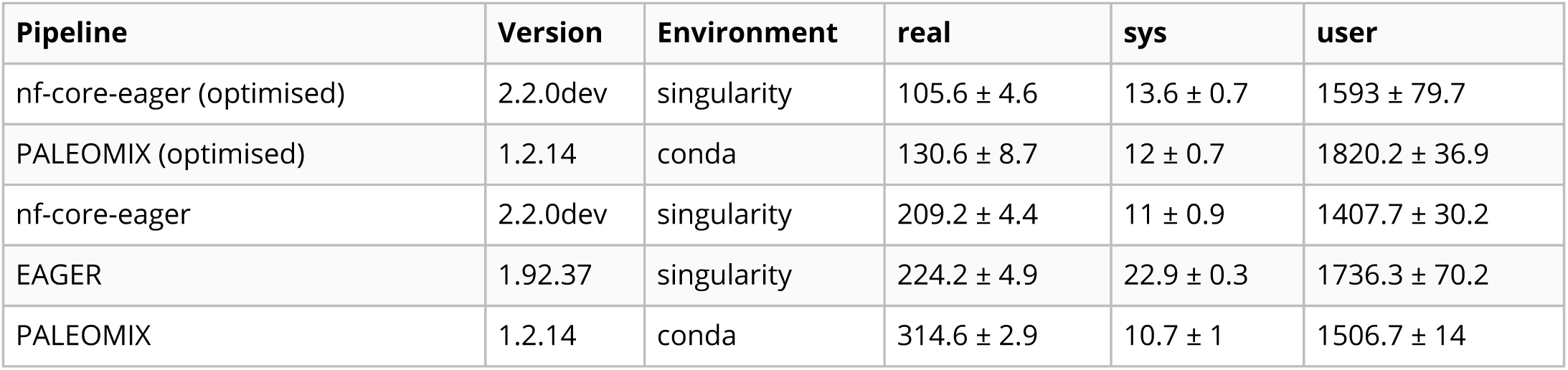
Comparison of run times in minutes between three ancient DNA pipelines. PALEOMIX and nf-core/eager have additional runs with ‘optimised’ parameters with fairer computational resources matching modern multi-threading strategies. Values represent mean and standard deviation of run times in minutes, calculated from the output of the GNU time tool. Real: real time, System: cumulative CPU system-task times, User: cumulative CPU time of all tasks.

| Pipeline | Version | Environment | real | sys | user |
| --- | --- | --- | --- | --- | --- |
| nf-core-eager (optimised) | 2.2.0dev | singularity | 105.6 ± 4.6 | 13.6 ± 0.7 | 1593 ± 79.7 |
| PALEOMIX (optimised) | 1.2.14 | conda | 130.6 ± 8.7 | 12 ± 0.7 | 1820.2 ± 36.9 |
| nf-core-eager | 2.2.0dev | singularity | 209.2 ± 4.4 | 11 ± 0.9 | 1407.7 ± 30.2 |
| EAGER | 1.92.37 | singularity | 224.2 ± 4.9 | 22.9 ± 0.3 | 1736.3 ± 70.2 |
| PALEOMIX | 1.2.14 | conda | 314.6 ± 2.9 | 10.7 ± 1 | 1506.7 ± 14 |

As a more realistic demonstration of modern computing multi-threading set ups, we also re-ran PALEOMIX with the flag –max-bwa-threads set to 4 (listed in Table 2 as ‘optimised’), which is equivalent to a single BWA aln process of nf-core/eager. This resulted in a much faster run-time than that of default nf-core/eager, due to the approach of PALEOMIX of mapping each lane of a library separately, whereas nf-core/eager will map all lanes of a single library merged together. Therefore, given that each library was split across three lanes, increasing the threads of BWA aln to 4 resulted in 12 per library, whereas nf-core/eager only gave 4 (by default) for a single BWA aln process of one library. While the PALEOMIX approach is valid, we opted to retain the per-library mapping as it is often the longest running step of high-throughput sequencing genome-mapping pipelines, and it prevents flooding of HPC scheduling systems with many long-running jobs. Secondly, if users regularly use multi-lane data, due to nf-core/eager’s fine-granularity control, they can simply modify nf-core/eager’s BWA aln process resources via config files to account for this. When we optimised parameters that were used for BWA aln’s multi-threading, and the number of multiple lanes to the same number of BWA aln threads as the optimised PALEOMIX run, nf-core/eager again displayed faster runtimes. All metrics including mapped reads, percentage on-target, mean depth coverage and mean read lengths across all pipelines were extremely similar across all pipelines and replicates (see methods and Table 3).

**Table 3:**
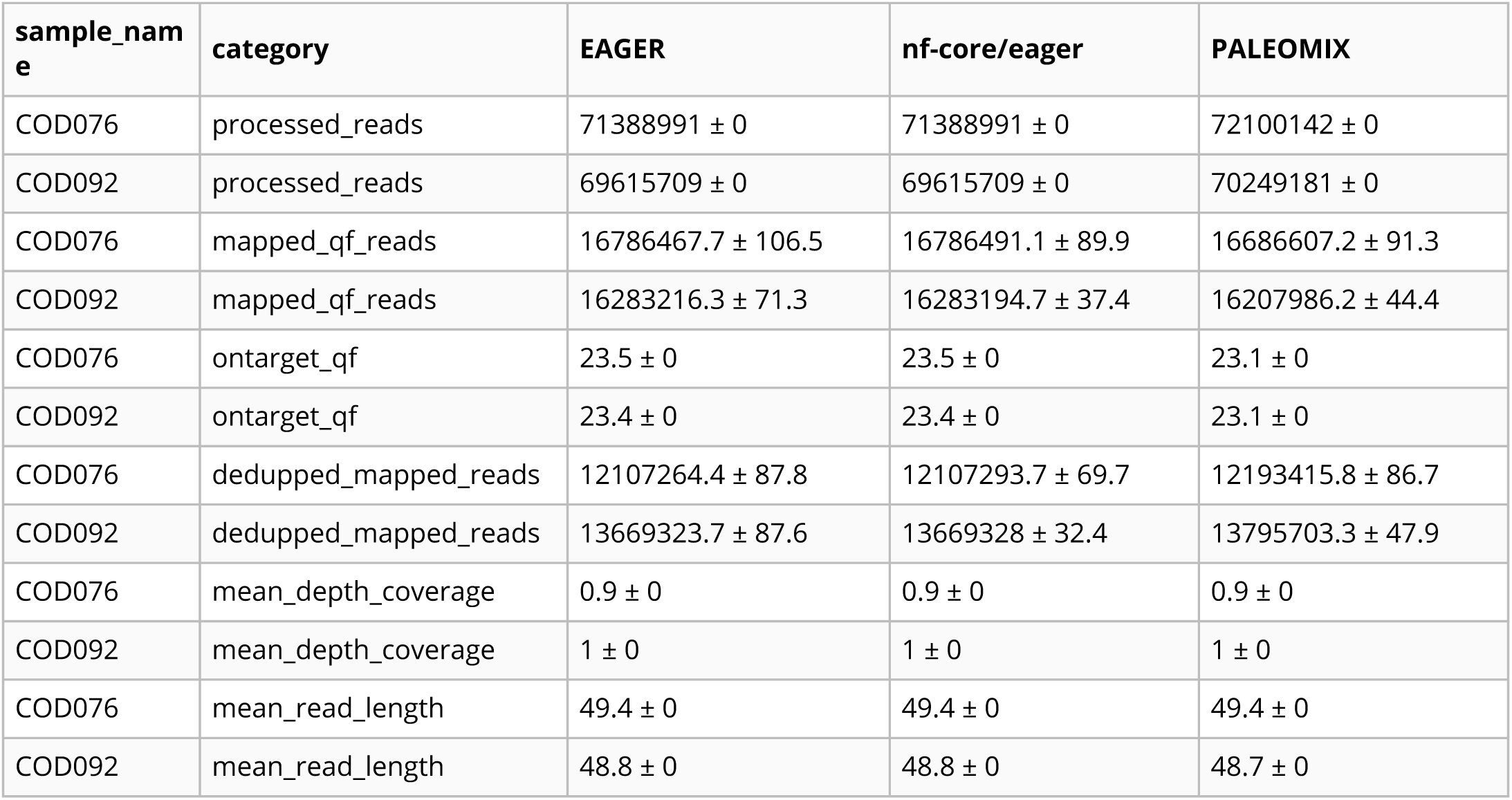
Comparison of common results values of key high-throughput short-read data processing and mapping steps across the three pipelines. ‘qf’ stands for mapping-quality filtered reads. All values represent mean and standard deviation across 10 replicates of each pipeline, calculated from the output of the GNU time tool.

## Conclusion

nf-core/eager is an efficient, portable, and accessible pipeline for processing and screening ancient (meta)genomic data. This re-implementation of EAGER into Nextflow and nf-core will improve reproducibility and scalability of rapidly increasing aDNA datasets, for both large and small laboratories. Extensive documentation also enables newcomers to the field to get a practical understanding on how to interpret aDNA in the context of NGS data processing. Ultimately, nf-core/eager provides easier access to the latest tools and routine screening analyses commonly used in the field, and sets up the pipeline for remaining at the forefront of palaeogenetic analysis.

## Methods

### Installation

nf-core/eager requires only three dependencies: Java (version >= 8), Nextflow, and either a functional Conda installation *or* Docker/Singularity engine installation. A quick installation guide to follow to get started can be found in the *Quick start* section of the nf-core/eager repository [84].

### Running

After installation, users can run the pipeline using standard test data by utilising some of the test profiles we provide (e.g. using Docker):

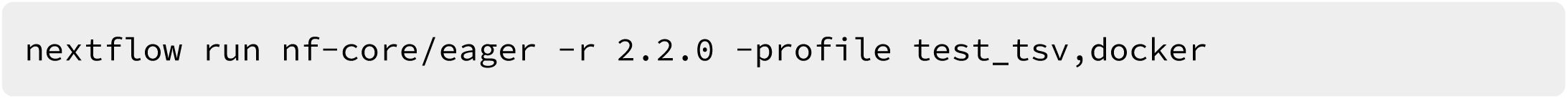

This will download test data automatically (as recorded in the test_tsv profile), run the pipeline locally with all software tools containerised in a Docker image. The pipeline will store the output of that run in the default ‘./results’ folder of the current directory.

The default pipeline settings assumes paired-end FASTQ data, and will run:

- FastQC
- AdapterRemoval2 (merging and adapter clipping)
- post-clipping FastQC (for AdapterRemoval2 performance evaluation) BWA mapping (with the ‘aln’ algorithm)
- samtools flagstat (for mapping statistics) endorS.py (for endogenous DNA calculation)
- Picard MarkDuplicates (for PCR amplicon deduplication) PreSeq (for library complexity evaluation)
- DamageProfiler and Qualimap2 (for genome coverage statistics)
- MultiQC pipeline run report

If no additional FASTA indices are given, these will also be generated.

The pipeline is highly configurable and most modules can be turned on-and-off using different flags at the request of the user, to allow a high level of customisation to each user’s needs. For example, to include metagenomic screening of off-target reads, and sex determination based on on-target mappings of pre-clipped single-end data:

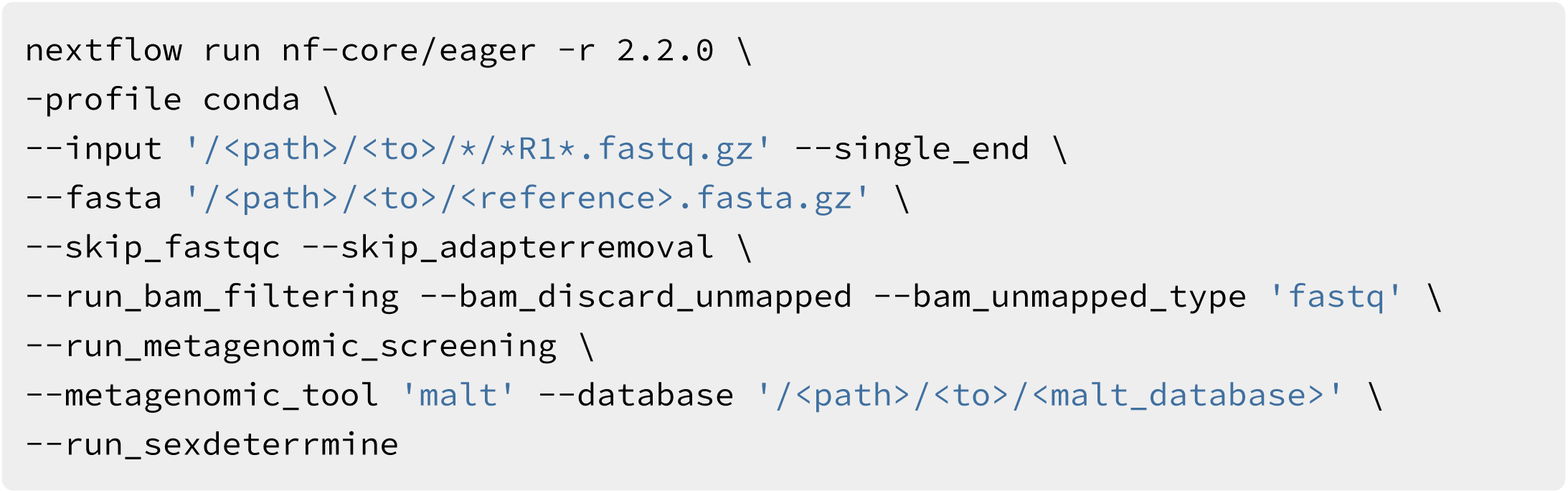

### Profiles

In addition to private locally defined profiles, we utilise a central configuration repository to enable users from various institutions to use pipelines on their particular infrastructure more easily [85]. There are multiple resources listed in this repository with information on how to add a user’s own institutional configuration profile with help from the nf-core community. These profiles can be both generic for all nf-core pipelines, but also customised for specific pipelines.

Users can customise this infrastructure profile by themselves, with the nf-core community, or with their local system administrator to make sure that the pipeline runs successfully, and can then rely on the Nextflow and nf-core framework to ensure compatibility upon further infrastructure changes. For example, in order to run the nf-core/eager pipeline at the Max Planck Institute for the Science of Human History (MPI-SHH), users only have to run:

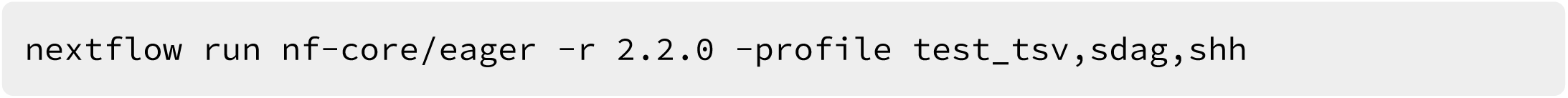

This runs the testing profile of the nf-core/eager pipeline with parameters specifically adapted to a specific HPC system at the MPI-SHH. In some cases, similar institutional configs for other institutions may already exist (originally utilised for different nf-core pipelines), so users need not necessarily write their own.

### Inputs

The pipeline can be started using (raw) FASTQ files from sequencing or pre-mapped BAM files. Additionally, the pipeline requires a FASTA reference genome. If BAM input is provided, an optional conversion to FASTQ is offered, otherwise BAM files processing will start from the post-mapping stage.

If users have complex set-ups, e.g. multiple sequencing lanes that require merging of files, the pipeline can be supplied with a tab separated value (TSV) file to enable such complex data handling. Both FASTQs and BAMs can be provided in this set up. FASTQs with the same library name and sequencing chemistry but sequenced across multiple lanes will be concatenated after adapter removal and prior mapping. Libraries with different sequencing chemistry kits (paired- vs. single-end) will be merged after mapping. Libraries with the same sample name and with the same UDG treatment, will be merged after deduplication. If libraries with the sample name have different UDG treatment, these will be merged after the aDNA modification stage (i.e. BAM trimming or PMDtools, if turned on), prior to genotyping, as shown in Figure 4.

**Figure 4:**
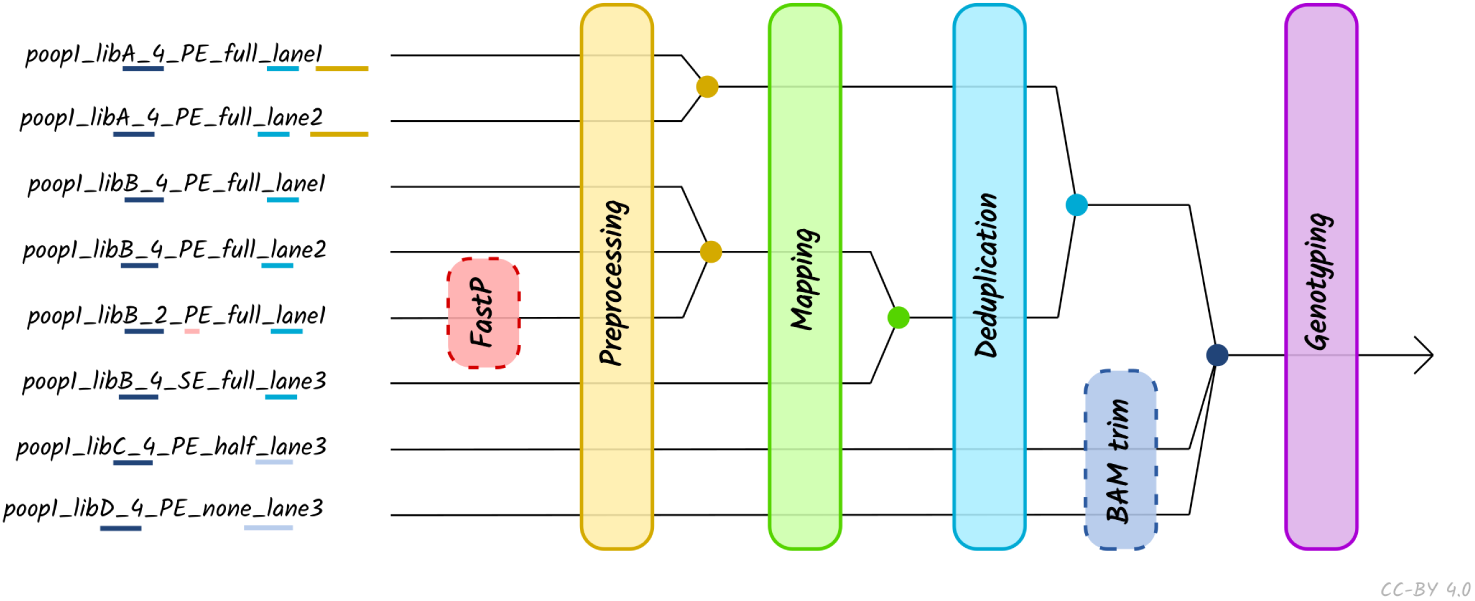
Schematic of different processing and merging points based on the nature of different libraries, as specified by the metadata of a TSV file. Dashed boxes represent optional library-specific processes. Colours refer to each merge points, which occur at certain points along the pipeline depending on the metadata columns defined in the TSV file.

As Nextflow will automatically download files from URLs, profiles and/or TSV files, users can include links to publicly available data (e.g. the European Bioinformatics Institutes’s ENA FTP server). This assists in reproducibility, because if profiles or TSV files are uploaded with a publication, a researcher wishing to re-analyse the data in the same way can use the exact settings and file merging procedures in the original publication, without having to reconstruct this from prose.

### Monitoring

Users can either monitor their pipeline execution with the messages Nextflow prints to the console while running, or utilise companion tools such as Nextflow’s Tower [50] to monitor their analysis pipeline during runtime.

### Output

The pipeline produces a multitude of output files in various file formats, with a more detailed listing available in the user documentation. These include metrics, statistical analysis data, and standardised output files (BAM, VCF) for close inspection and further downstream analysis, as well as a MultiQC report. If an emailing daemon is set up on the server, the latter can be emailed to users automatically, when starting the pipeline with a dedicated option (--).

### Benchmarking

#### Dual Screening of Human and Microbial Pathogen DNA

Full step-by-step instructions on the set up of the human and pathogen screening demonstration (including input TSV file) can be seen in the supplementary information. To demonstrate the efficiency and conciseness of nf-core/eager pipeline in it’s dual role for both human and microbial screening of ancient material, we replicated the results of Barquera et al. 2020 [81] using v2.2.0 (commit: e7471a7 and Nextflow version: 20.04.1).

The following command was used to run the pipeline on the in-house servers at the MPI-SHH, including a 2 TB memory node for running MALT against the NCBI Nt (Nucleotide) database, and therefore the centralised custom profile for this cluster was used.

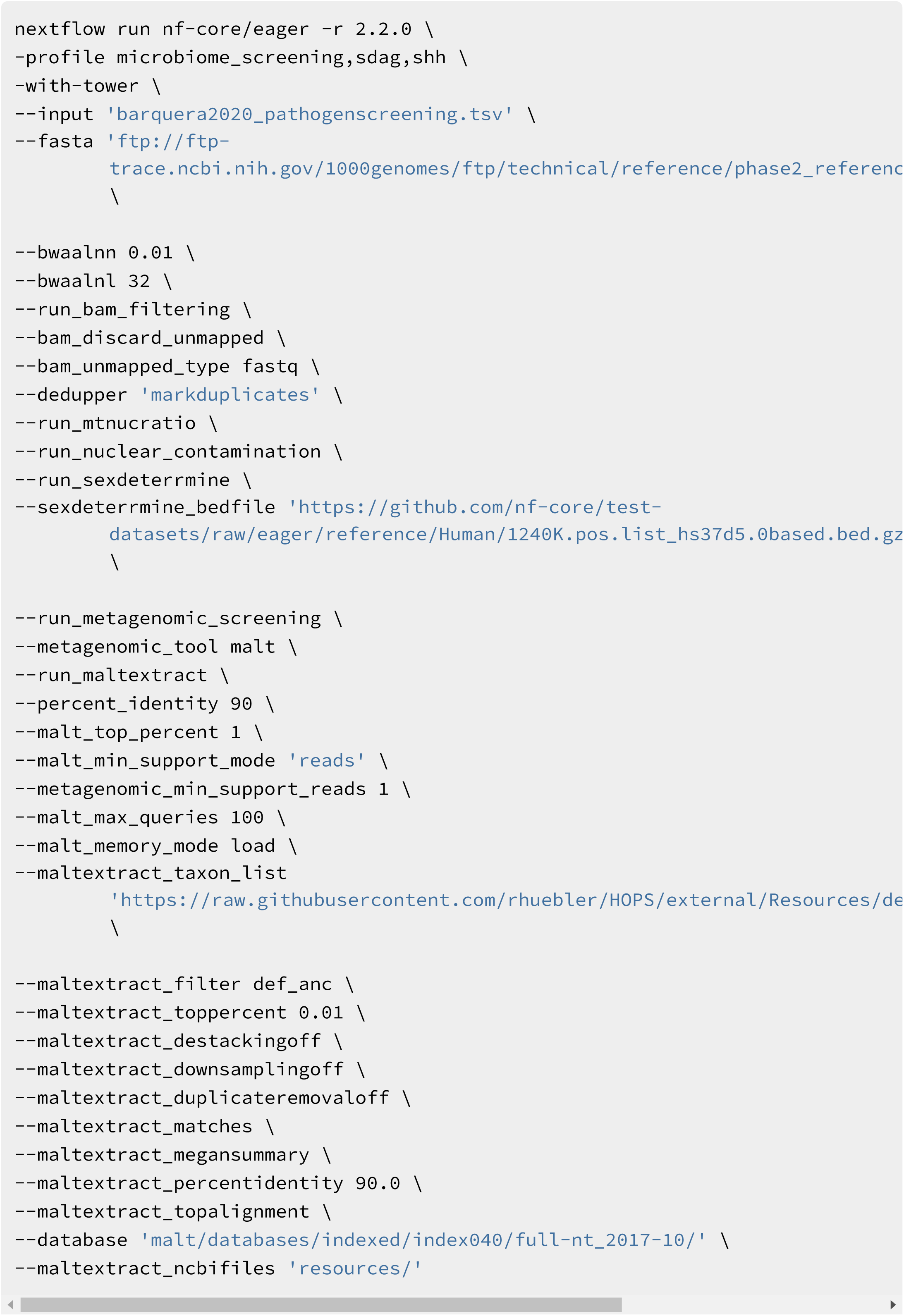

To include the HOPS results from metagenomic screening in the report, we also re-ran MultiQC with the upcoming version v1.10 (to be integrated into nf-core/eager on release). After then installing the development version of MultiQC (commit: 7584e64), as described in the MultiQC documentation [86], we ran the following command in the results directory of the nf-core/eager run, using the same configuration file.

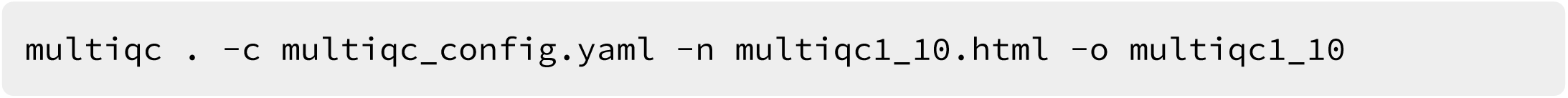

Until MultiQC v1.10 is released, the HOPS heatmap is exported by nf-core/eager in the corresponding MaltExtract results directory. Reports from both versions (and the standalone HOPS PDF) can be seen in the supplementary information.

### Pipeline Comparison

Full step-by-step instructions on the set up of the pipeline run-time benchmarking, including environment and tool versions, can be seen in the supplementary information. EAGER (v1.92.37) and nf-core/eager (v2.2.0, commit: 830c22d; Nextflow v20.04.1) used the provided pre-built singularity containers for software environments, whereas for PALEOMIX (v1.2.14) we generated a custom conda environment (see supplementary information for the `environmental.yaml` file). Run time comparisons were performed on a 32 CPU (AMD Opteron 23xx) and 256 GB memory Red Hat QEMU Virtual Machine running the Ubuntu 18.04 operating system (Linux Kernel 4.15.0-112). Resource parameters of each tool were only modified to specify the maximum available on the server and otherwise left as default.

The following commands were used for each pipeline, with the commands run 10 times, each after cleaning up reference and results directories using a for loop. Run times of the run commands themselves were measured using GNU Time.

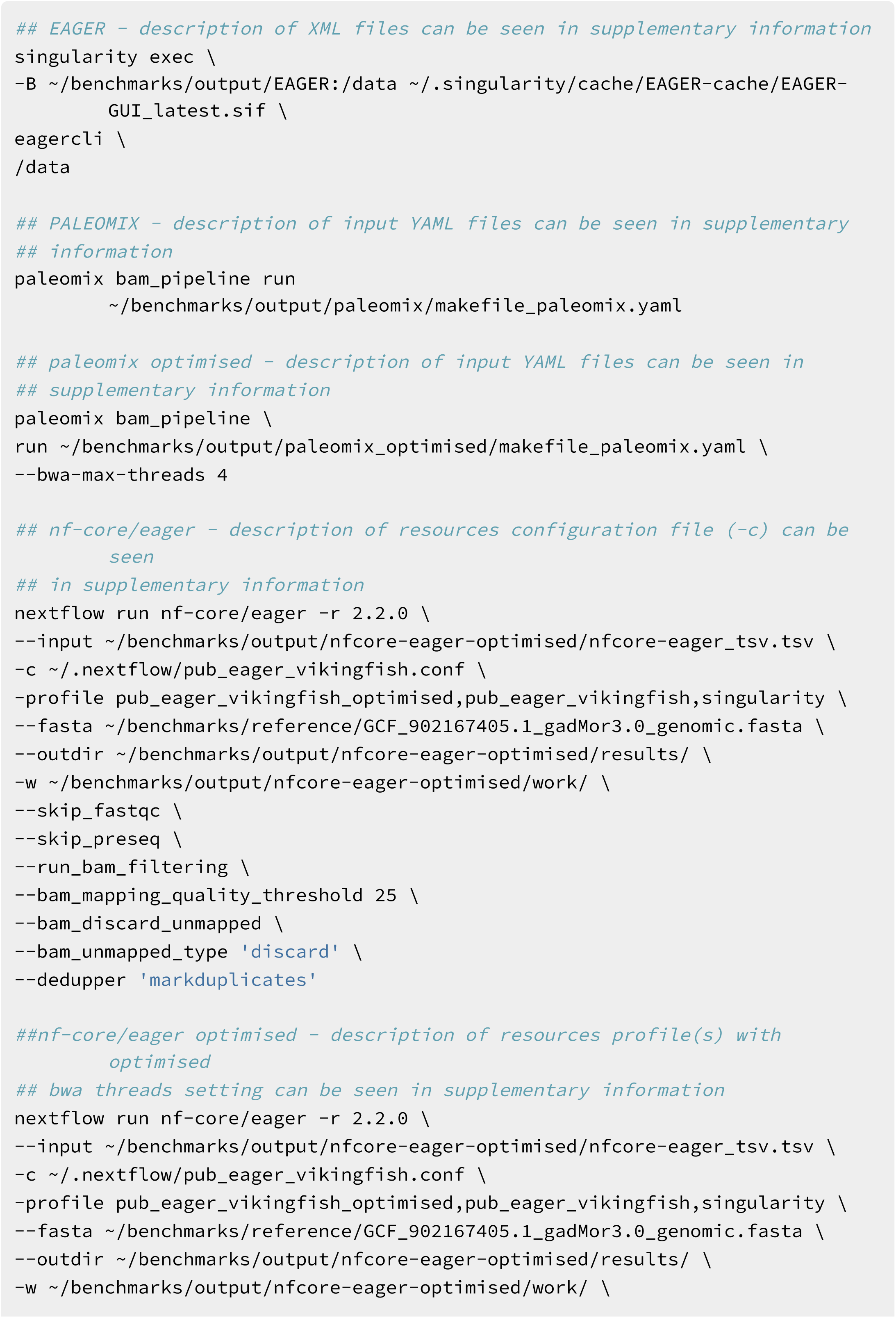
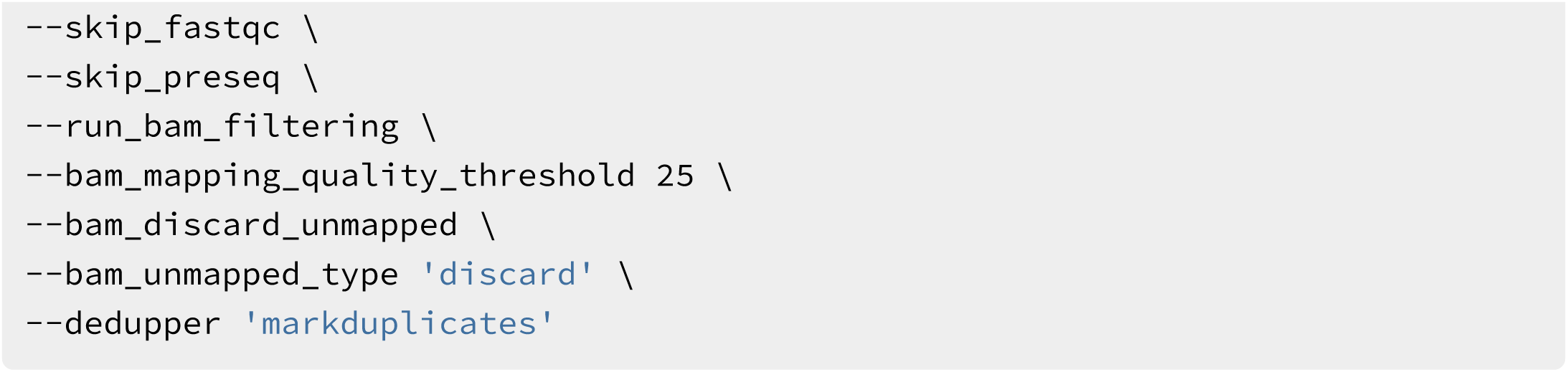

Mapping results across all pipelines showed very similar values, with low variation across replicates as can be seen in Table 3.

## Supporting information

Supplemental file archive of walkthrough and results files

## Data and software availability

All pipeline code is available on GitHub at https://github.com/nf-core/eager and archived with Zenodo under the DOI 10.5281/zenodo.1465061. The version of nf-core/eager that this manuscript is based on was the ‘dev’ branch of the GitHub repository (2.2.0dev), and was released as v2.2.0.

Demonstration data for dual ancient human and pathogen screening from Barquera et al. [81] is publicly available on the European Nucleotide Archive (ENA) under project accession PRJEB37490. The human reference genome (hs37d5) and screening database (Nucleotide or ‘nt’, October 2017) was downloaded from National Center for Biotechnology Information FTP server. Ancient Cod genomic data from Star et al. [4] used for benchmarking is publicly available on the ENA under project accession PRJEB20524. The *Gadus morhua* reference genome NCBI accession ID is: GCF_902167405.1.

This paper was collaboratively written with Manubot [87], and supplementary information including demonstration and benchmarking environments descriptions and walk-through can be seen on GitHub at https://github.com/apeltzer/eager2-paper/ and the `supplement/` directory.

## Acknowledgements

We thank the nf-core community for general support and suggestions during the writing of the pipeline. We also thank Arielle Munters, Hester van Schalkwyk, Irina Velsko, Katherine Eaton, Luc Venturini, Marcel Keller, Pierre Lindenbaum, Pontus Skoglund, Raphael Eisenhofer, Torsten Günter, Kevin Lord, and Åshild Vågene for bug reports and feature suggestions. We are grateful to the members of the Department of Archaeogenetics at the Max Planck Institute for the Science of Human History who performed beta testing of the pipeline. We thank the aDNA twitter community for responding to polls regarding design decisions during development.

The Gesellschaft für wissenschaftliche Datenverarbeitung (GWDG, Göttingen) kindly provided computational infrastructure for benchmarking. We also want to thank Selina Carlhoff, Maria Spyrou, Elisabeth Nelson, Alexander Herbig and Wolfgang Haak for providing comments and suggestions on this manuscript, and acknowledge Christina Warinner, Stephan Schiffels and the Max Planck Society who provided funds for travel to nf-core events. This project was also supported by the ERC Starting Grant project (FoodTransforms) ERC-2015-StG 678901 funded by the European Research Council awarded to Philipp W. Stockhammer (Ludwig Maximilian University, Munich).

