## Supplemental file archive of walkthrough and results files for "Reproducible, portable, and efficient ancient genome reconstruction with nf-core/eager": benchmarking_results_parsing.html

nf-core/eager Benchmarking Results Comparisons


### nf-core/eager Benchmarking Results Comparisons

###### James A. Fellows Yates

#### Notebook Setup

This notebook will use the `tidyverse` set of packages for data loading, manipulation and plotting, and `knitr` for easy printing of markdown tables for the main paper.

```
library(tidyverse)
```

```
## ── Attaching packages ──────────────────────────────────────────────────────────────────────────────────────────────────────────────────────────────── tidyverse 1.3.0 ──
```

```
## ✓ ggplot2 3.3.2     ✓ purrr   0.3.4
## ✓ tibble  3.0.3     ✓ dplyr   1.0.1
## ✓ tidyr   1.1.1     ✓ stringr 1.4.0
## ✓ readr   1.3.1     ✓ forcats 0.5.0
```

```
## ── Conflicts ─────────────────────────────────────────────────────────────────────────────────────────────────────────────────────────────────── tidyverse_conflicts() ──
## x dplyr::filter() masks stats::filter()
## x dplyr::lag()    masks stats::lag()
```

```
library(knitr)

sessionInfo()
```

```
## R version 3.6.3 (2020-02-29)
## Platform: x86_64-pc-linux-gnu (64-bit)
## Running under: Ubuntu 20.04.1 LTS
## 
## Matrix products: default
## BLAS:   /usr/lib/x86_64-linux-gnu/blas/libblas.so.3.9.0
## LAPACK: /usr/lib/x86_64-linux-gnu/lapack/liblapack.so.3.9.0
## 
## locale:
##  [1] LC_CTYPE=en_GB.UTF-8       LC_NUMERIC=C              
##  [3] LC_TIME=en_GB.UTF-8        LC_COLLATE=en_GB.UTF-8    
##  [5] LC_MONETARY=en_GB.UTF-8    LC_MESSAGES=en_GB.UTF-8   
##  [7] LC_PAPER=en_GB.UTF-8       LC_NAME=C                 
##  [9] LC_ADDRESS=C               LC_TELEPHONE=C            
## [11] LC_MEASUREMENT=en_GB.UTF-8 LC_IDENTIFICATION=C       
## 
## attached base packages:
## [1] stats     graphics  grDevices utils     datasets  methods   base     
## 
## other attached packages:
##  [1] knitr_1.29      forcats_0.5.0   stringr_1.4.0   dplyr_1.0.1    
##  [5] purrr_0.3.4     readr_1.3.1     tidyr_1.1.1     tibble_3.0.3   
##  [9] ggplot2_3.3.2   tidyverse_1.3.0
## 
## loaded via a namespace (and not attached):
##  [1] Rcpp_1.0.5       cellranger_1.1.0 pillar_1.4.6     compiler_3.6.3  
##  [5] dbplyr_1.4.4     tools_3.6.3      digest_0.6.25    lubridate_1.7.9 
##  [9] jsonlite_1.7.0   evaluate_0.14    lifecycle_0.2.0  gtable_0.3.0    
## [13] pkgconfig_2.0.3  rlang_0.4.7      reprex_0.3.0     cli_2.0.2       
## [17] rstudioapi_0.11  DBI_1.1.0        yaml_2.2.1       haven_2.3.1     
## [21] xfun_0.16        withr_2.2.0      xml2_1.3.2       httr_1.4.2      
## [25] fs_1.5.0         hms_0.5.3        generics_0.0.2   vctrs_0.3.2     
## [29] grid_3.6.3       tidyselect_1.1.0 glue_1.4.1       R6_2.4.1        
## [33] fansi_0.4.1      readxl_1.3.1     rmarkdown_2.3    modelr_0.1.8    
## [37] blob_1.2.1       magrittr_1.5     backports_1.1.8  scales_1.1.1    
## [41] ellipsis_0.3.1   htmltools_0.5.0  rvest_0.3.6      assertthat_0.2.1
## [45] colorspace_1.4-1 stringi_1.4.6    munsell_0.5.0    broom_0.7.0     
## [49] crayon_1.3.4
```

#### Notebook Infrastructure

Due to heterogeneous reporting across each pipeline, we need to specify which are comparable and standardise the column names.

```
## paleomix only reports QF reads so will only use those
eager_cols <- c(
  replicate = "Replicate",
  sample_name = "Sample Name",
  processed_reads = "# reads after C&M prior mapping",
  mapped_qf_reads = "# mapped reads prior RMDup QF",
  ontarget_qf = "Endogenous DNA QF (%)",
  dedupped_mapped_reads = "Mapped Reads after RMDup",
  mean_depth_coverage = "Mean Coverage",
  mean_read_length = "average fragment length"
)

nfcoreeager_cols <- c(
  replicate = "Replicate",
  sample_name = "Sample",
  processed_reads = "Samtools Flagstat (pre-samtools filter)_mqc-generalstats-samtools_flagstat_pre_samtools_filter-flagstat_total",
  mapped_qf_reads = "Samtools Flagstat (post-samtools filter)_mqc-generalstats-samtools_flagstat_post_samtools_filter-mapped_passed",
  ontarget_qf = "endorSpy_mqc-generalstats-endorspy-endogenous_dna_post",
  dedupped_mapped_reads = "QualiMap_mqc-generalstats-qualimap-mapped_reads",
  mean_depth_coverage = "QualiMap_mqc-generalstats-qualimap-mean_coverage",
  mean_read_length = "DamageProfiler_mqc-generalstats-damageprofiler-mean_readlength"
)

paleomix_cols <- c(
  replicate = "Replicate",
  sample_name = "Target",
  processed_reads = "seq_retained_reads",
  mapped_qf_reads = "hits_raw(GCF_902167405.1_gadMor3.0_genomic)",
  ontarget_qf = "hits_raw_frac(GCF_902167405.1_gadMor3.0_genomic)",
  dedupped_mapped_reads = "hits_unique(GCF_902167405.1_gadMor3.0_genomic)",
  mean_depth_coverage = "hits_coverage(GCF_902167405.1_gadMor3.0_genomic)",
  mean_read_length = "hits_length(GCF_902167405.1_gadMor3.0_genomic)"
)

## Function to standardise colum names using above lists to allow easy joining
rename_col <- function(x, list, inverted = T) {
  if (inverted) {
    replacement_list <- names(list)
    names(replacement_list) <- list
    replacement_list[x]
  } else {
    names(list[x])
  }
}
```

#### Data Loading

Next we can actually load the files

For EAGER and nf-core/eager

```
raw_eager <- Sys.glob("results/Report_EAGER*") %>% 
  enframe(name = NULL, value = "File") %>% 
  mutate(Replicate = map(File, ~tools::file_path_sans_ext(.x) %>% str_split("_") %>% unlist %>% tail(n = 1)) %>% unlist,
         Contents = map(File, ~read_csv(.x))) %>%
  unnest() %>%
  filter(Replicate != 1)
```

```
## Warning: `cols` is now required when using unnest().
## Please use `cols = c(Contents)`
```

```
raw_nfcoreeager <- Sys.glob("results/nf*") %>%
  enframe(name = NULL, value = "File") %>% 
  mutate(Replicate = map(File, ~tools::file_path_sans_ext(.x) %>% str_split("_") %>% unlist %>% tail(n = 1)) %>% unlist,
         Contents = map(File, ~read_tsv(.x))) %>%
  unnest() %>%
  filter(Replicate != 1,
         !grepl("_", Sample)) %>%
  mutate(Sample = str_sub(Sample, 1, 6))
```

```
## Warning: `cols` is now required when using unnest().
## Please use `cols = c(Contents)`
```

And for paleomix

```
raw_paleomix <- Sys.glob("results/paleomix_C*") %>%
  enframe(name = NULL, value = "File") %>%
  mutate(Replicate = map(File, ~tools::file_path_sans_ext(.x) %>% str_split("_") %>% unlist %>% tail(n = 1)) %>% unlist,
         Contents = map(File, ~read_tsv(.x, comment = "#"))) %>%
  unnest() %>%
  filter(Measure != "lib_type") %>%
  mutate(Value = as.numeric(Value)) %>%
  pivot_wider(names_from = Measure, values_from = Value) %>%
  filter(Replicate != 1) %>%
  mutate(`hits_raw_frac(GCF_902167405.1_gadMor3.0_genomic)` = `hits_raw_frac(GCF_902167405.1_gadMor3.0_genomic)` * 100)
```

```
## Warning: `cols` is now required when using unnest().
## Please use `cols = c(Contents)`
```

#### Standardisation and Joining

The following function will allow us to make the input data of all pipelines into a standard (long) format. It will then summarise across the replicates.

```
standardise_summarise <- function(x, cols) {
  x %>%
    select(cols) %>%
    group_by(sample_name) %>%
    select(-replicate) %>%
    pivot_longer(cols = !contains("sample_name"), names_to = "category", values_to = "value") %>%
    group_by(sample_name, category) %>%
    summarise(
      Mean = round(mean(value), digits = 1),
      SD = round(sd(value), digits = 1)
    ) %>%
    unite(col = "mean_sd_values", Mean, SD, sep = " ± ")
}
```

And now we can combine

```
combined <- bind_rows(
  standardise_summarise(raw_eager, eager_cols) %>% mutate(pipeline = "eager"),
  standardise_summarise(raw_nfcoreeager, nfcoreeager_cols) %>% mutate(pipeline = "nf-core-eager"),
  standardise_summarise(raw_paleomix, paleomix_cols) %>% mutate(pipeline = "paleomix")) %>%
  select(pipeline, everything()) %>%
  pivot_wider(names_from = pipeline, values_from = mean_sd_values) %>%
  mutate(category = factor(category, levels = c("processed_reads", "mapped_qf_reads", "ontarget_qf", "dedupped_mapped_reads", "mean_depth_coverage", "mean_read_length"))) %>%
  arrange(category)
```

```
## Note: Using an external vector in selections is ambiguous.
## ℹ Use `all_of(cols)` instead of `cols` to silence this message.
## ℹ See <https://tidyselect.r-lib.org/reference/faq-external-vector.html>.
## This message is displayed once per session.
```

```
## `summarise()` regrouping output by 'sample_name' (override with `.groups` argument)
## `summarise()` regrouping output by 'sample_name' (override with `.groups` argument)
## `summarise()` regrouping output by 'sample_name' (override with `.groups` argument)
```

#### Reporting

Finally we can plot as markdown for transfer into the main paper.

```
combined %>%
  kable()
```

| sample\_name | category | eager | nf-core-eager | paleomix |
| --- | --- | --- | --- | --- |
| COD076 | processed\_reads | 71388991 ± 0 | 71388991 ± 0 | 72100142 ± 0 |
| COD092 | processed\_reads | 69615709 ± 0 | 69615709 ± 0 | 70249181 ± 0 |
| COD076 | mapped\_qf\_reads | 16786467.7 ± 106.5 | 16786491.1 ± 89.9 | 16686607.2 ± 91.3 |
| COD092 | mapped\_qf\_reads | 16283216.3 ± 71.3 | 16283194.7 ± 37.4 | 16207986.2 ± 44.4 |
| COD076 | ontarget\_qf | 23.5 ± 0 | 23.5 ± 0 | 23.1 ± 0 |
| COD092 | ontarget\_qf | 23.4 ± 0 | 23.4 ± 0 | 23.1 ± 0 |
| COD076 | dedupped\_mapped\_reads | 12107264.4 ± 87.8 | 12107293.7 ± 69.7 | 12193415.8 ± 86.7 |
| COD092 | dedupped\_mapped\_reads | 13669323.7 ± 87.6 | 13669328 ± 32.4 | 13795703.3 ± 47.9 |
| COD076 | mean\_depth\_coverage | 0.9 ± 0 | 0.9 ± 0 | 0.9 ± 0 |
| COD092 | mean\_depth\_coverage | 1 ± 0 | 1 ± 0 | 1 ± 0 |
| COD076 | mean\_read\_length | 49.4 ± 0 | 49.4 ± 0 | 49.4 ± 0 |
| COD092 | mean\_read\_length | 48.8 ± 0 | 48.8 ± 0 | 48.7 ± 0 |
