## Supplemental file archive of walkthrough and results files for "Reproducible, portable, and efficient ancient genome reconstruction with nf-core/eager": benchmarking_runtime_parsing.html

nf-core/eager Runtime Benchmarking Comparisons


### nf-core/eager Runtime Benchmarking Comparisons

###### James A. Fellows Yates

#### Notebook Setup

This notebook will use the `tidyverse` set of packages for data loading, manipulation and plotting, and `knitr` for easy printing of markdown tables for the main paper.

#### Data Loading

We will load the pre-aggregated and downloaded runtimes as recorded by the GNU `time` unix utility

```
results <- read_tsv("benchmarking_aggregated_runtimes.txt", 
                    col_names = c("Run", "Runtime"))
```

```
## Parsed with column specification:
## cols(
##   Run = col_character(),
##   Runtime = col_character()
## )
```

#### Data Cleaning

Next we can clean up the file names to find the corresponding pipeline name.

```
results_clean <- results %>%
  separate(col = Run, sep = ":", c("File", "Line", "Category")) %>%
  select(-Line) %>%
  mutate(
    File = str_remove(File, "time_") %>%
      str_remove(".log") %>%
      str_remove("runtimes/") %>%
      str_replace("nf-core_eager", "nf-core/eager") %>%
      str_replace("paleomix_optimised", "paleomix-optimised"),
    Runtime_Minutes = map(Runtime, ~ str_split(.x, "m") %>%
      unlist() %>%
      unlist() %>%
      pluck(1)) %>% unlist() %>% as.numeric()
  ) %>%
  separate(File, sep = "_", into = c("Pipeline", "Replicate")) %>%
  select(-Runtime) %>%
  filter(Replicate != 1)
```

#### Data Summaries

To get the final results we will summarise the mean and standard deviation of the three runtime metrics and print the table as markdown.

```
results_final_tidy <- results_clean %>%
  group_by(Pipeline, Category) %>%
  summarise(
    Mean = round(mean(Runtime_Minutes), digits = 1),
    SD = round(sd(Runtime_Minutes), digits = 1)
  ) %>%
  arrange(Category, Mean)
```

```
## `summarise()` regrouping output by 'Pipeline' (override with `.groups` argument)
```

```
results_final_print <- results_final_tidy %>%
  unite(col = "Mean Runtime", Mean, SD, sep = " ± ") %>%
  pivot_wider(names_from = Category, values_from = `Mean Runtime`) %>%
  kable()

results_final_print
```

| Pipeline | real | sys | user |
| --- | --- | --- | --- |
| nf-core-eager-optimised | 105.6 ± 4.6 | 13.6 ± 0.7 | 1593 ± 79.7 |
| paleomix-optimised | 130.6 ± 8.7 | 12 ± 0.7 | 1820.2 ± 36.9 |
| nf-core-eager | 209.2 ± 4.4 | 11 ± 0.9 | 1407.7 ± 30.2 |
| EAGER | 224.2 ± 4.9 | 22.9 ± 0.3 | 1736.3 ± 70.2 |
| paleomix | 314.6 ± 2.9 | 10.7 ± 1 | 1506.7 ± 14 |

#### Data Plotting

We can also plot the results.

```
## Get get order of fastest to slowest based on real time
results_to_plot <- results_clean %>% mutate(Pipeline = factor(Pipeline, levels = rev(results_final_tidy$Pipeline %>% unique)))

ggplot(results_to_plot, aes(Runtime_Minutes, Pipeline)) +
  geom_violin(aes(colour = Pipeline)) +
  geom_point(pch = 20, alpha = 0.7) +
  xlab("Runtime (minutes)") +
  facet_wrap(~Category, scales = "free_x") +
  scale_colour_brewer(palette = "Set1", guide = guide_legend(nrow = 2)) +
  theme_minimal() +
  theme(legend.position = "bottom")
```
