## Supplemental file archive of walkthrough and results files for "Reproducible, portable, and efficient ancient genome reconstruction with nf-core/eager": benchmarking_runtime_parsing.Rmd

````
---
title: "nf-core/eager Runtime Benchmarking Comparisons"
author: "James A. Fellows Yates"
output:
  html_document:
    df_print: paged
---

## Notebook Setup

This notebook will use the `tidyverse` set of packages for data loading, 
manipulation and plotting, and `knitr` for easy printing of markdown tables
for the main paper.

```{r}
library(tidyverse)
library(knitr)

sessionInfo()
```

## Data Loading

We will load the pre-aggregated and downloaded runtimes as recorded by the 
GNU `time` unix utility

```{r}
results <- read_tsv("benchmarking_aggregated_runtimes.txt", 
                    col_names = c("Run", "Runtime"))
```

## Data Cleaning

Next we can clean up the file names to find the corresponding pipeline name.

```{r}
results_clean <- results %>%
  separate(col = Run, sep = ":", c("File", "Line", "Category")) %>%
  select(-Line) %>%
  mutate(
    File = str_remove(File, "time_") %>%
      str_remove(".log") %>%
      str_remove("runtimes/") %>%
      str_replace("nf-core_eager", "nf-core/eager") %>%
      str_replace("paleomix_optimised", "paleomix-optimised"),
    Runtime_Minutes = map(Runtime, ~ str_split(.x, "m") %>%
      unlist() %>%
      unlist() %>%
      pluck(1)) %>% unlist() %>% as.numeric()
  ) %>%
  separate(File, sep = "_", into = c("Pipeline", "Replicate")) %>%
  select(-Runtime) %>%
  filter(Replicate != 1)

```

## Data Summaries

To get the final results we will summarise the mean and standard deviation
of the three runtime metrics across all replicats per-pipeline and print the 
table as markdown.

```{r}
results_final_tidy <- results_clean %>%
  group_by(Pipeline, Category) %>%
  summarise(
    Mean = round(mean(Runtime_Minutes), digits = 1),
    SD = round(sd(Runtime_Minutes), digits = 1)
  ) %>%
  arrange(Category, Mean)

results_final_print <- results_final_tidy %>%
  unite(col = "Mean Runtime", Mean, SD, sep = " ± ") %>%
  pivot_wider(names_from = Category, values_from = `Mean Runtime`) %>%
  kable()

results_final_print

```

## Data Plotting

We can also plot the results.

```{r, fig.height=3, fig.width=7}
## Get get order of fastest to slowest based on real time
results_to_plot <- results_clean %>% mutate(Pipeline = factor(Pipeline, levels = rev(results_final_tidy$Pipeline %>% unique)))

ggplot(results_to_plot, aes(Runtime_Minutes, Pipeline)) +
  geom_violin(aes(colour = Pipeline)) +
  geom_point(pch = 20, alpha = 0.7) +
  xlab("Runtime (minutes)") +
  facet_wrap(~Category, scales = "free_x") +
  scale_colour_brewer(palette = "Set1", guide = guide_legend(nrow = 2)) +
  theme_minimal() +
  theme(legend.position = "bottom")
  
  

```
````
