## Supplemental file archive of walkthrough and results files for "Reproducible, portable, and efficient ancient genome reconstruction with nf-core/eager": benchmarking_server_environment_setup.md

# nf-core/eager paper - benchmarking

This document describes the set up of server environment used to benchmark
nf-core/eager against two other popular aDNA NGS pipelines: EAGER vs. Paleomix.

The server environment was constructed on a node instance of the GWDG cloud
server. The server OS was set to Ubuntu 18.04.

## Server setup

When logging in for server preparation and EAGER set up, ensure to log in with
`shh -X` so you can open X11 windows. All other sessions do not require this.

### Preparation

Ensure everything is up-to-date

```bash
sudo apt update
sudo apt upgrade
sudo apt autoremove
```

Check X11 forwarding is available

```bash
sudo apt install x11-apps
xeyes
```

We will also use aspera for faster file downloading of test data

```bash
cd ~/bin/
wget https://download.asperasoft.com/download/sw/cli/3.9.6/ibm-aspera-cli-3.9.6.1467.159c5b1-linux-64-release.sh
sh ibm-aspera-cli-3.9.6.1467.159c5b1-linux-64-release.sh
echo 'export PATH=/home/cloud/.aspera/cli/bin:$PATH' >> ~/.bashrc
source ~/.bashrc
```

Install rename for speedy regex file rename and tree for faster folder structure
look up

```bash
sudo apt install rename
sudo apt tree
```

Install XMLlint for easier reading of EAGER XML files

```bash
sudo apt-get install libxml2-utils
```

For easy hardware summaries

```bash
sudo apt install hwinfo
```

Some personal alias

```bash
echo "alias ltree='tree -C | less -R'" >> ~/.bashrc
echo "alias ll='ls -alFh'" >> ~/.bashrc
echo "alias l='ls -CF'" >> ~/.bashrc

```

### nf-core/eager

Install Java

```bash
## Java
sudo apt install openjdk-11-jdk
```

Now install Nextflow and do a little bit of housekeeping

```bash
## Nextflow
mkdir bin && cd !$
curl -s https://get.nextflow.io | bash

# Put nextflow in PATH, not necessary but :shrug:
echo 'PATH=$PATH:/home/cloud/bin/' > ~/.bashrc
source ~/.bashrc

## Run test
nextflow run hello
```

Should see the following:

```txt
N E X T F L O W  ~  version 20.04.1
Pulling nextflow-io/hello ...
downloaded from https://github.com/nextflow-io/hello.git
Launching `nextflow-io/hello` [distracted_torvalds] - revision: ef60d89ab9 [master]
WARN: The use of `echo` method is deprecated
executor >  local (4)
[d6/104882] process > sayHello (2) [100%] 4 of 4 ✔
Ciao world!

Hola world!

Bonjour world!

Hello world!
```

We will use Singularity as our container system, as this is supported by both
nf-core/eager and EAGER v1. This requires a little more set up over conda,
however it is more robust.

```bash
cd ~
## Singularity Dependencies
sudo apt-get update && sudo apt-get install -y \
    build-essential \
    libssl-dev \
    uuid-dev \
    libgpgme11-dev \
    squashfs-tools \
    libseccomp-dev \
    wget \
    pkg-config \
    git \
    cryptsetup

## Install Go
curl -O https://dl.google.com/go/go1.14.4.linux-amd64.tar.gz
tar xvf go1.14.4.linux-amd64.tar.gz
sudo chown -R root:root ./go
sudo mv go /usr/local
echo 'export GOPATH=$HOME/work
export PATH=$PATH:/usr/local/go/bin:$GOPATH/bin' >> ~/.bashrc
 source ~/.bashrc
rm go1.14.4.linux-amd64.tar.gz

# Singularity itself
export VERSION=3.5.2 && # adjust this as necessary \
    wget https://github.com/sylabs/singularity/releases/download/v${VERSION}/singularity-${VERSION}.tar.gz && \
    tar -xzf singularity-${VERSION}.tar.gz && \
    cd singularity

    ./mconfig && \
    make -C builddir && \
    sudo make -C builddir install

cd ~
rm singularity-3.5.2.tar.gz
mv singularity/ bin/

## Now we will set up a singularity cache so all containers are in the same
## place
mkdir -p .singularity/cache

echo 'export NXF_SINGULARITY_CACHEDIR=/home/cloud/.singularity/cache/nxf-cache' >> ~/.bashrc
source ~/.bashrc
```

Now we can test that nf-core/eager will run. Note the testing version is
2.2.0dev, in the `dev` revision/version.

```bash
## get nf-core/eager
mkdir nfcore-eager_test
cd !$
nextflow pull nf-core/eager -r dev
nextflow run nf-core/eager -r dev -profile test_tsv,singularity
cd ~
```

This pulled commit hash: 830c22d448441e5e19508c198f530a7656c9f25d

Finally, we will generate a 'config' file that sets the maximum resources of the
particular server we are using.

```bash
nano ~/.nextflow/pub_eager_vikingfish.conf
```

Add the following and save. This modifies the CPUs to 32, mem to 250 (leaving 6
for nextflow itself and other background processes), and boost the wall-times
for each process to a large number. These would normally be customised for each
machine, and typical dataset.

```text
profiles {
  pub_eager_vikingfish {
    process {
      cpus = { check_max( 1 * task.attempt, 'cpus' ) }
      memory = { check_max( 7.GB * task.attempt, 'memory' ) }
      time = { check_max( 4.h * task.attempt, 'time' ) }

      errorStrategy = { task.exitStatus in [143,137,104,134,139] ? 'retry' : 'finish' }
      maxRetries = 3
      maxErrors = '-1'

      // Generic resource requirements - s(ingle)c(ore)/m(ulti)c(ore)

      withLabel:'sc_tiny'{
          cpus = { check_max( 1, 'cpus' ) }
          memory = { check_max( 1.GB * task.attempt, 'memory' ) }
          time = { check_max( 24.h * task.attempt, 'time' ) }
      }

      withLabel:'sc_small'{
          cpus = { check_max( 1, 'cpus' ) }
          memory = { check_max( 4.GB * task.attempt, 'memory' ) }
          time = { check_max( 24.h * task.attempt, 'time' ) }
      }

      withLabel:'sc_medium'{
          cpus = { check_max( 1, 'cpus' ) }
          memory = { check_max( 8.GB * task.attempt, 'memory' ) }
          time = { check_max( 24.h * task.attempt, 'time' ) }
      }

      withLabel:'mc_small'{
          cpus = { check_max( 2, 'cpus' ) }
          memory = { check_max( 4.GB * task.attempt, 'memory' ) }
          time = { check_max( 24.h * task.attempt, 'time' ) }
      }

      withLabel:'mc_medium' {
          cpus = { check_max( 4, 'cpus' ) }
          memory = { check_max( 8.GB * task.attempt, 'memory' ) }
          time = { check_max( 24.h * task.attempt, 'time' ) }
      }

      withLabel:'mc_large'{
          cpus = { check_max( 8, 'cpus' ) }
          memory = { check_max( 16.GB * task.attempt, 'memory' ) }
          time = { check_max( 24.h * task.attempt, 'time' ) }
      }

      withLabel:'mc_huge'{
          cpus = { check_max( 32, 'cpus' ) }
          memory = { check_max( 256.GB * task.attempt, 'memory' ) }
          time = { check_max( 24.h * task.attempt, 'time' ) }
      }

      // Process-specific resource requirements (others leave at default, e.g. Fastqc)
      withName:get_software_versions {
        memory = { check_max( 2.GB, 'memory' ) }
        cache = false
      }

      withName:qualimap{
        errorStrategy = 'ignore'
      }

      withName:preseq {
        errorStrategy = 'ignore'
      }

      // Add 141 ignore due to unclean pipe closing by pmdtools https://github.com/pontussk/PMDtools/issues/7
      withName: pmdtools {
        errorStrategy = { task.exitStatus in [141] ? 'ignore' : 'retry' }
      }

      // Add 1 retry as not enough heapspace java error gives exit code 1
      withName: malt {
        errorStrategy = { task.exitStatus in [1] ? 'retry' : 'finish' }
      }

      withName: multiqc {
        errorStrategy = { task.exitStatus in [143,137] ? 'retry' : 'ignore' }
      }
    }

    params {
      // Defaults only, expecting to be overwritten
      max_memory = 250.GB
      max_cpus = 32
      max_time = 240.h
      igenomes_base = 's3://ngi-igenomes/igenomes/'
      config_profile_description = 'Profile for nf-core/eager publication test environment'
    }
  }
  pub_eager_vikingfish_optimised {
    process {
      withName: bwa {
        cpus = { check_max( 12, 'cpus' ) }
      }
    }
  }
}

def check_max(obj, type) {
  if (type == 'memory') {
    try {
      if (obj.compareTo(params.max_memory as nextflow.util.MemoryUnit) == 1)
        return params.max_memory as nextflow.util.MemoryUnit
      else
        return obj
    } catch (all) {
      println "   ### ERROR ###   Max memory '${params.max_memory}' is not valid! Using default value: $obj"
      return obj
    }
  } else if (type == 'time') {
    try {
      if (obj.compareTo(params.max_time as nextflow.util.Duration) == 1)
        return params.max_time as nextflow.util.Duration
      else
        return obj
    } catch (all) {
      println "   ### ERROR ###   Max time '${params.max_time}' is not valid! Using default value: $obj"
      return obj
    }
  } else if (type == 'cpus') {
    try {
      return Math.min( obj, params.max_cpus as int )
    } catch (all) {
      println "   ### ERROR ###   Max cpus '${params.max_cpus}' is not valid! Using default value: $obj"
      return obj
    }
  }
}

```

### PALEOMIX

The [documentation](https://paleomix.readthedocs.io/en/latest/) for PALEOMIX was
a little confusing, so this required a bit of back and forth. For safety I
resorted to using a conda environment rather than the `virtualenv` thing, as it
does a similar thing but I can also contain all the additional software
dependencies.

The virtual env thing doesn't actually include all the software, as I
erroneously thought first time around. Lets try conda instead. Will use 2.7 as
all of PALAEOMIX is in Python2.

Firstly install conda and set up so it can use bioconda, where most of the
bioinformatics tools are derived from.

```bash
cd ~/bin
wget https://repo.anaconda.com/miniconda/Miniconda2-latest-Linux-x86_64.sh
chmod +x Miniconda2-latest-Linux-x86_64.sh
./Miniconda2-latest-Linux-x86_64.sh

## Follow instructions, install under /home/cloud/bin/miniconda2
## Do run init
~/bin/miniconda2/bin/conda config --set auto_activate_base false

## Log out and back in again

conda config --add channels defaults
conda config --add channels bioconda
conda config --add channels conda-forge
```

Now we can create an environment specifically for PALAEOMIX (postscript: to
speed this up I've added an `paleomix_environment.yaml` file than you can then
use instead with `conda env create -f paleomix_environment.yaml`)

```bash
# Make conda environment; note adding missing GATK and R requirement(s) not in docs: https://github.com/MikkelSchubert/paleomix/issues/28
conda create -n paleomix python=2.7 pip=20.1.1 adapterremoval=2.3.1 samtools=1.9 picard=2.22.9 bowtie2=2.3.5.1 bwa=0.7.17 mapdamage2=2.0.9 gatk=3.8 r-base=3.5.1 r-rcpp=1.0.4.6 r-rcppgsl=0.3.7 r-gam=1.16.1 r-inline=0.3.15

conda activate paleomix

## While in the paleomix environment, install paleomix
pip install --user paleomix

## For safety add ~/.local/bin to path
echo 'export PATH=$PATH:/home/cloud/.local/bin' >> ~/.bashrc
source ~/.bashrc

conda activate paleomix
```

Ok now we have some hardcoded dependencies to fix. Firstly we need to get the
GATK JAR, because it isn't actually allowed to be shipped in the bioconda GATK
bioconda recipe because of licensing. We can then symlink this into the ugly
folder PALEOMIX wants for the test (at least). We can also symlink the picard
JAR that came with the bioconda environment.

```bash
wget https://storage.googleapis.com/gatk-software/package-archive/gatk/GenomeAnalysisTK-3.8-1-0-gf15c1c3ef.tar.bz2
gatk3-register GenomeAnalysisTK-3.8-1-0-gf15c1c3ef.tar.bz2
mkdir -p /home/cloud/install/jar_root/
ln -s /home/cloud/bin/miniconda2/envs/paleomix/opt/gatk-3.8/GenomeAnalysisTK.jar /home/cloud/install/jar_root/
ln -s /home/cloud/bin/miniconda2/envs/paleomix/share/picard-2.22.9-0/picard.jar /home/cloud/install/jar_root/
rm ~/Genome*.tar.bz2
```

Now we can test that paleomix is working properly.

```bash
cd ~
paleomix bam_pipeline example .
cd ~/bam_pipeline
paleomix bam_pipeline run 000_makefile.yaml

```

We can generate a global settings file to make sure it runs properly, and bump
resources to match of this server.

Note we will only change the maximum of the server. All other values are kept as
default.

```bash
paleomix bam_pipeline run --write-config
sed -i 's/max_threads = 16/max_threads = 32/g' ~/.paleomix/bam_pipeline.ini
```

### EAGER

Now finally we can set up EAGER v1.

Get the singularity image

```bash
mkdir -p ~/.singularity/cache/EAGER-cache
singularity pull shub://apeltzer/EAGER-GUI

## Test loading
mkdir -p ~/EAGER-test
cd !$

## Download mini test data also used for nf-core/eager
mkdir reference output input/JK2782
wget https://github.com/nf-core/test-datasets/raw/eager/testdata/Mammoth/fastq/JK2782_TGGCCGATCAACGA_L008_{R1,R2}_001.fastq.gz.tengrand.fq.gz
mv *gz input/JK2782
wget https://raw.githubusercontent.com/nf-core/test-datasets/eager/reference/Mammoth/Mammoth_MT_Krause.fna
mv *fna reference


singularity exec -B .:/data ~/.singularity/cache/EAGER-cache/EAGER-GUI_latest.sif eager

## See output/JK2782 for XML file, set mostly on defaults

## To actually run EAGER
singularity exec -B .:/data ~/.singularity/cache/EAGER-cache/EAGER-GUI_latest.sif eagercli /data

```

### Versions

To record all versions of all the install tools, we can see the output of the
following commands

```bash
## Kernel
uname -r
4.15.0-112-generic

## OS
lsb_release -a
No LSB modules are available.
Distributor ID: Ubuntu
Description:    Ubuntu 18.04.4 LTS
Release:    18.04
Codename:   bionic

## Java
java -version
openjdk version "11.0.7" 2020-04-14
OpenJDK Runtime Environment (build 11.0.7+10-post-Ubuntu-2ubuntu218.04)
OpenJDK 64-Bit Server VM (build 11.0.7+10-post-Ubuntu-2ubuntu218.04, mixed mode, sharing)

## Nextflow
nextflow -version

      N E X T F L O W
      version 20.04.1 build 5335
      created 03-05-2020 19:37 UTC (21:37 CEST)
      cite doi:10.1038/nbt.3820
      http://nextflow.io

## Singularity (for nf-core/eager and EAGER)
singularity --version
singularity version 3.5.2

## Conda (for palaeomix)
conda --version
conda 4.7.12

paleomix
PALEOMIX - pipelines and tools for NGS data analyses.
Version: 1.2.14

## For EAGER
cat ~/EAGER-test/output/Report_output_versions.txt
EAGER-CLI    1.92.55
EAGER-GUI   1.92.37
ReportTable Version     1.92.33

## For measuring run time
/usr/bin/time
GNU time 1.7

# hardware
lscpu
sudo lshw -short
hwinfo --short

```

### Benchmarking Data

For a straightforward testing dataset, we will be mapping shotgun sequencing
data of Viking Age cod (fish), from Star et al. ([2017,
PNAS](https://dx.doi.org/10.1073/pnas.1710186114)). For run-time purposes, we
will use a subset of these samples.

We will download the following sequencing data from the [EBI's ENA
archive](https://www.ebi.ac.uk/ena), and the reference genome from the ENA FTP
server.

#### Table 1 | List of sequencing data used for benchmarking

| Sample_Name | Library_ID     | Lane | study_accession | run_accession | tax_id | scientific_name | instrument_model    | library_layout | fastq_ftp                                                                                                                                         | fastq_aspera                                                                                                                                          | submitted_ftp                                                                                                                                                                                         |
|-------------|----------------|------|-----------------|---------------|--------|-----------------|---------------------|----------------|---------------------------------------------------------------------------------------------------------------------------------------------------|-------------------------------------------------------------------------------------------------------------------------------------------------------|-------------------------------------------------------------------------------------------------------------------------------------------------------------------------------------------------------|
| COD076      | COD076E1bL1    |    8 | PRJEB20524      | ERR1943600    |   8049 | Gadus morhua    | Illumina HiSeq 2500 | PAIRED         | ftp.sra.ebi.ac.uk/vol1/fastq/ERR194/000/ERR1943600/ERR1943600_1.fastq.gz;ftp.sra.ebi.ac.uk/vol1/fastq/ERR194/000/ERR1943600/ERR1943600_2.fastq.gz | fasp.sra.ebi.ac.uk:/vol1/fastq/ERR194/000/ERR1943600/ERR1943600_1.fastq.gz;fasp.sra.ebi.ac.uk:/vol1/fastq/ERR194/000/ERR1943600/ERR1943600_2.fastq.gz | ftp.sra.ebi.ac.uk/vol1/run/ERR194/ERR1943600/COD076E1bL1_CTCGCGC_L008_R1_001.fastq.gz;ftp.sra.ebi.ac.uk/vol1/run/ERR194/ERR1943600/COD076E1bL1_CTCGCGC_L008_R2_001.fastq.gz                           |
| COD076      | COD076E1bL1    |    6 | PRJEB20524      | ERR1943601    |   8049 | Gadus morhua    | Illumina HiSeq 2500 | PAIRED         | ftp.sra.ebi.ac.uk/vol1/fastq/ERR194/001/ERR1943601/ERR1943601_1.fastq.gz;ftp.sra.ebi.ac.uk/vol1/fastq/ERR194/001/ERR1943601/ERR1943601_2.fastq.gz | fasp.sra.ebi.ac.uk:/vol1/fastq/ERR194/001/ERR1943601/ERR1943601_1.fastq.gz;fasp.sra.ebi.ac.uk:/vol1/fastq/ERR194/001/ERR1943601/ERR1943601_2.fastq.gz | ftp.sra.ebi.ac.uk/vol1/run/ERR194/ERR1943601/COD076E1bL1_CTCGCGC_L006_R1_001.fastq.gz;ftp.sra.ebi.ac.uk/vol1/run/ERR194/ERR1943601/COD076E1bL1_CTCGCGC_L006_R2_001.fastq.gz                           |
| COD076      | COD076E1bL1    |    1 | PRJEB20524      | ERR1943602    |   8049 | Gadus morhua    | Illumina HiSeq 2500 | PAIRED         | ftp.sra.ebi.ac.uk/vol1/fastq/ERR194/002/ERR1943602/ERR1943602_1.fastq.gz;ftp.sra.ebi.ac.uk/vol1/fastq/ERR194/002/ERR1943602/ERR1943602_2.fastq.gz | fasp.sra.ebi.ac.uk:/vol1/fastq/ERR194/002/ERR1943602/ERR1943602_1.fastq.gz;fasp.sra.ebi.ac.uk:/vol1/fastq/ERR194/002/ERR1943602/ERR1943602_2.fastq.gz | ftp.sra.ebi.ac.uk/vol1/run/ERR194/ERR1943602/C9NP5ANXX_COD076E1bL1_CTCGCGC_L001_R1_001.fastq.gz;ftp.sra.ebi.ac.uk/vol1/run/ERR194/ERR1943602/C9NP5ANXX_COD076E1bL1_CTCGCGC_L001_R2_001.fastq.gz       |
| COD092      | COD092E1bL1i69 |    6 | PRJEB20524      | ERR1943607    |   8049 | Gadus morhua    | Illumina HiSeq 2500 | PAIRED         | ftp.sra.ebi.ac.uk/vol1/fastq/ERR194/007/ERR1943607/ERR1943607_1.fastq.gz;ftp.sra.ebi.ac.uk/vol1/fastq/ERR194/007/ERR1943607/ERR1943607_2.fastq.gz | fasp.sra.ebi.ac.uk:/vol1/fastq/ERR194/007/ERR1943607/ERR1943607_1.fastq.gz;fasp.sra.ebi.ac.uk:/vol1/fastq/ERR194/007/ERR1943607/ERR1943607_2.fastq.gz | ftp.sra.ebi.ac.uk/vol1/run/ERR194/ERR1943607/C9VJJANXX_COD092E1bL1i69_AACCTGC_L006_R1_001.fastq.gz;ftp.sra.ebi.ac.uk/vol1/run/ERR194/ERR1943607/C9VJJANXX_COD092E1bL1i69_AACCTGC_L006_R2_001.fastq.gz |
| COD092      | COD092E1bL1i69 |    7 | PRJEB20524      | ERR1943608    |   8049 | Gadus morhua    | Illumina HiSeq 2500 | PAIRED         | ftp.sra.ebi.ac.uk/vol1/fastq/ERR194/008/ERR1943608/ERR1943608_1.fastq.gz;ftp.sra.ebi.ac.uk/vol1/fastq/ERR194/008/ERR1943608/ERR1943608_2.fastq.gz | fasp.sra.ebi.ac.uk:/vol1/fastq/ERR194/008/ERR1943608/ERR1943608_1.fastq.gz;fasp.sra.ebi.ac.uk:/vol1/fastq/ERR194/008/ERR1943608/ERR1943608_2.fastq.gz | ftp.sra.ebi.ac.uk/vol1/run/ERR194/ERR1943608/C9VJJANXX_COD092E1bL1i69_AACCTGC_L007_R1_001.fastq.gz;ftp.sra.ebi.ac.uk/vol1/run/ERR194/ERR1943608/C9VJJANXX_COD092E1bL1i69_AACCTGC_L007_R2_001.fastq.gz |
| COD092      | COD092E1bL1i69 |    8 | PRJEB20524      | ERR1943609    |   8049 | Gadus morhua    | Illumina HiSeq 2500 | PAIRED         | ftp.sra.ebi.ac.uk/vol1/fastq/ERR194/009/ERR1943609/ERR1943609_1.fastq.gz;ftp.sra.ebi.ac.uk/vol1/fastq/ERR194/009/ERR1943609/ERR1943609_2.fastq.gz | fasp.sra.ebi.ac.uk:/vol1/fastq/ERR194/009/ERR1943609/ERR1943609_1.fastq.gz;fasp.sra.ebi.ac.uk:/vol1/fastq/ERR194/009/ERR1943609/ERR1943609_2.fastq.gz | ftp.sra.ebi.ac.uk/vol1/run/ERR194/ERR1943609/C9VJJANXX_COD092E1bL1i69_AACCTGC_L008_R1_001.fastq.gz;ftp.sra.ebi.ac.uk/vol1/run/ERR194/ERR1943609/C9VJJANXX_COD092E1bL1i69_AACCTGC_L008_R2_001.fastq.gz |
Alternatively, for the ASPERA version

The reference genome can be downloaded from the [NCBI's SRA FTP
server](http://ftp.sra.ebi.ac.uk/) from
[https://ftp.ncbi.nlm.nih.gov/genomes/refseq/vertebrate_other/Gadus_morhua/representative/GCF_902167405.1_gadMor3.0/GCF_902167405.1_gadMor3.0_genomic.fna.gz](https://ftp.ncbi.nlm.nih.gov/genomes/refseq/vertebrate_other/Gadus_morhua/representative/GCF_902167405.1_gadMor3.0/GCF_902167405.1_gadMor3.0_genomic.fna.gz)

```bash
mkdir -p ~/benchmarks/input ~/benchmarks/output ~/benchmarks/reference
cd ~/benchmarks/input
```

We will place these in directories and rename in a form primarily compatible
with EAGER, which is most heavily dependent on folder structure.

```bash
screen -R downloads

for i in fasp.sra.ebi.ac.uk:/vol1/fastq/ERR194/000/ERR1943600/ERR1943600_1.fastq.gz fasp.sra.ebi.ac.uk:/vol1/fastq/ERR194/000/ERR1943600/ERR1943600_2.fastq.gz fasp.sra.ebi.ac.uk:/vol1/fastq/ERR194/001/ERR1943601/ERR1943601_1.fastq.gz fasp.sra.ebi.ac.uk:/vol1/fastq/ERR194/001/ERR1943601/ERR1943601_2.fastq.gz fasp.sra.ebi.ac.uk:/vol1/fastq/ERR194/002/ERR1943602/ERR1943602_1.fastq.gz fasp.sra.ebi.ac.uk:/vol1/fastq/ERR194/002/ERR1943602/ERR1943602_2.fastq.gz fasp.sra.ebi.ac.uk:/vol1/fastq/ERR194/007/ERR1943607/ERR1943607_1.fastq.gz fasp.sra.ebi.ac.uk:/vol1/fastq/ERR194/007/ERR1943607/ERR1943607_2.fastq.gz fasp.sra.ebi.ac.uk:/vol1/fastq/ERR194/008/ERR1943608/ERR1943608_1.fastq.gz fasp.sra.ebi.ac.uk:/vol1/fastq/ERR194/008/ERR1943608/ERR1943608_2.fastq.gz fasp.sra.ebi.ac.uk:/vol1/fastq/ERR194/009/ERR1943609/ERR1943609_1.fastq.gz fasp.sra.ebi.ac.uk:/vol1/fastq/ERR194/009/ERR1943609/ERR1943609_2.fastq.gz; do
    ascp -QT -l 300m -P33001 \
    -i /home/cloud/.aspera/cli/etc/asperaweb_id_dsa.openssh \
    era-fasp@"$i" \
    .
done

mkdir COD076 COD092

mv ERR194360{0..2}_* COD076
mv ERR194360{7..9}_* COD092

rename s/_/_R/ */*.gz
rename s/.fastq/_000.fastq/ */*.gz
rename s/_R/_S0_L000_R/ */*.gz

## Now manually rename the Lane for each one
rename s/ERR1943600_S0_L000_R/ERR1943600_S0_L008_R/ */*.gz
rename s/ERR1943601_S0_L000_R/ERR1943601_S0_L006_R/ */*.gz
rename s/ERR1943602_S0_L000_R/ERR1943602_S0_L001_R/ */*.gz
rename s/ERR1943607_S0_L000_R/ERR1943607_S0_L006_R/ */*.gz
rename s/ERR1943608_S0_L000_R/ERR1943608_S0_L007_R/ */*.gz
rename s/ERR1943609_S0_L000_R/ERR1943609_S0_L008_R/ */*.gz

## And make sample names consistent
rename s/ERR.*_S/COD076E1bL1_S/ COD076/*.gz
rename s/ERR.*_S/COD092E1bL1i69_S/ COD092/*.gz
```

To download the reference genome

```bash
cd ~/benchmarks/reference
wget https://ftp.ncbi.nlm.nih.gov/genomes/refseq/vertebrate_other/Gadus_morhua/representative/GCF_902167405.1_gadMor3.0/GCF_902167405.1_gadMor3.0_genomic.fna.gz

## unzip as EAGER can't cope with gzipped
gunzip GCF_902167405.1_gadMor3.0_genomic.fna.gz
## Rename to .fasta as paleomix can't cope
rename s/.fna/.fasta/
```

As each pipeline can generate reference indices themselves, (and we don't want
one pipeline to generate all of them for us), we will make symlinks into each
pipelines' own reference directory.

```bash
mkdir EAGER/ paleomix/ nfcore-eager/
ln -s ~/benchmarks/reference/GCF_902167405.1_gadMor3.0_genomic.fasta ~/benchmarks/reference/EAGER/
ln -s ~/benchmarks/reference/GCF_902167405.1_gadMor3.0_genomic.fasta ~/benchmarks/reference/paleomix/
ln -s ~/benchmarks/reference/GCF_902167405.1_gadMor3.0_genomic.fasta ~/benchmarks/reference/nfcore-eager/

rename 's/.fna/.fasta/' ~/benchmarks/reference/paleomix/*fna -n
```

## Benchmarking

### EAGER1 Setup Instructions

Set up XMLs

```bash
## ensure logged in with shh -X to open the GUI window!
cd ~/benchmarks
mkdir -p output/EAGER
singularity exec -B .:/data ~/.singularity/cache/EAGER-cache/EAGER-GUI_latest.sif eager
```

#### Table 2 | Input settings for EAGER benchmark. All values are default unless specified. Modules run are FastQC, AdapterRemoval, Mapping, RemoveDuplicates, Damage calculation

Section | Field | Value
--------|------|------
Input   | Path | ~/benchmarks/input/
Input   | Organism Type | Other
Input   | Age of Dataset | Ancient
Input   | Treated Data   | non-UDG Treated
Input   | Pairment | Paired Data |
Input   | Capture Data | FALSE
Input   | Calculate on target | FALSE
Input   | Input is already concatenated (skip merging) | FALSE
Input   | Concatenate lanewise together | TRUE
Input   | MTCapture Data | FALSE
Output  | Path | ~/benchmarks/output/EAGER
Reference | Path | ~/benchmarks/reference/EAGER/GCF_902167405.1_gadMor3.0_genomic.fna
Reference | Name of mitochondrial chromosome | NC_002081.1 Gadus
Resources | CPU Cores | 32
Resources | Memory in GB | 250
FastQC | Activate | FALSE
Mapping | Activate | TRUE
Mapping | Tool | BWA
Mapping | BWA SeedLength (-l) | 1024
Mapping | Max #diff (-n) | 0.04
Mapping | BWA Qualityfilter (-q) | 25
Remove Duplicates | MarkDuplicates
Damage Calculation | Activate | True
Damage Calculation | Tool | DamageProfiler

Note that the pipeline does now allow the `--mm` parameter for AdapterRemoval2,
and will not be exactly comparable to paleomix

> Memory is set to slightly less than total on system to allow buffer space.

```bash
## Important! Before doing this log out and log back in again WITHOUT -X in the shh command, else damageProfiler will crash!
screen -R EAGER
cd ~/benchmarks/output

for i in {1..10}; do
    { time singularity exec -B ./EAGER:/data ~/.singularity/cache/EAGER-cache/EAGER-GUI_latest.sif eagercli /data ; } 2> time_EAGER_"$i".log
    if [[ $i != 10 ]]; then
         rm -r ~/benchmarks/output/EAGER/*/*/ ~/benchmarks/output/EAGER/*.{csv,html,png,ReportGenerator,txt} EAGER/*/*.log ~/benchmarks/output/EAGER/*/DONE*
         rm ~/benchmarks/reference/EAGER*.{dict,amb,ann,bwt,fai,pac,sa} ~/benchmarks/reference/EAGER/DONE*
    fi
done
```

### Paleomix Setup

```bash
cd ~/benchmarks
mkdir -p output/paleomix
paleomix bam_pipeline mkfile > output/paleomix/makefile_paleomix.yaml
```

Now modify the makefile to match our data

```bash
sed -i 's/CompressionFormat: bz2/CompressionFormat: gz/' output/paleomix/makefile_paleomix.yaml
sed -i 's/--mm: 3/# --mm: 3/' output/paleomix/makefile_paleomix.yaml ## remove as EAGER1 doesn't provide this option
sed -i 's/--minlength: 25/--minlength: 30/' output/paleomix/makefile_paleomix.yaml
sed -i 's/MinQuality: 0/MinQuality: 25/' output/paleomix/makefile_paleomix.yaml # To follow Star paper
sed -i 's/NAME_OF_PREFIX:/GCF_902167405.1_gadMor3.0_genomic:/' output/paleomix/makefile_paleomix.yaml
sed -i 's+Path: PATH_TO_PREFIX+Path: /home/cloud/benchmarks/reference/paleomix/GCF_902167405.1_gadMor3.0_genomic.fasta+' output/paleomix/makefile_paleomix.yaml
sed -i 's/UseSeed: yes/UseSeed: no/' output/paleomix/makefile_paleomix.yaml

sed -i 's/#NAME_OF_TARGET:/COD076:/' output/paleomix/makefile_paleomix.yaml
sed -i 's/#  NAME_OF_SAMPLE:/  COD076:/' output/paleomix/makefile_paleomix.yaml
sed -i 's/#    NAME_OF_LIBRARY:/    COD076E1bL1:/' output/paleomix/makefile_paleomix.yaml
sed -i 's+#      NAME_OF_LANE: PATH_WITH_WILDCARDS+      Lane_8: /home/cloud/benchmarks/input/COD076/COD076E1bL1_S0_L008_R{Pair}_*.fastq.gz+' output/paleomix/makefile_paleomix.yaml
echo "      Lane_6: /home/cloud/benchmarks/input/COD076/COD076E1bL1_S0_L006_R{Pair}_*.fastq.gz" >> output/paleomix/makefile_paleomix.yaml
echo "      Lane_1: /home/cloud/benchmarks/input/COD076/COD076E1bL1_S0_L001_R{Pair}_*.fastq.gz" >> output/paleomix/makefile_paleomix.yaml
echo "" >> output/paleomix/makefile_paleomix.yaml
echo "COD092:" >>  output/paleomix/makefile_paleomix.yaml
echo "  COD092:" >>  output/paleomix/makefile_paleomix.yaml
echo "    COD092E1bL1i69:" >>  output/paleomix/makefile_paleomix.yaml
echo "      Lane_6: /home/cloud/benchmarks/input/COD092/COD092E1bL1i69_S0_L006_R{Pair}_*.fastq.gz" >> output/paleomix/makefile_paleomix.yaml
echo "      Lane_7: /home/cloud/benchmarks/input/COD092/COD092E1bL1i69_S0_L007_R{Pair}_*.fastq.gz" >> output/paleomix/makefile_paleomix.yaml
echo "      Lane_8: /home/cloud/benchmarks/input/COD092/COD092E1bL1i69_S0_L008_R{Pair}_*.fastq.gz" >> output/paleomix/makefile_paleomix.yaml
```

To then run

```bash
conda activate paleomix
cd ~/benchmarks/output/paleomix

for i in {1..10}; do
    { time paleomix bam_pipeline run makefile_paleomix.yaml ; } 2> ../time_paleomix_"$i".log
    if [[ $i != 10 ]]; then
         rm -r ~/benchmarks/output/paleomix/!(makefile_paleomix.yaml)
         rm ~/benchmarks/reference/paleomix/*.{dict,amb,ann,bwt,fai,pac,sa,validated}

    fi
done
```

## paleomix optimised

The paleomix run however is not configured very nicely, as the
`bam_pipeline.ini` does not multi-thread mapping steps, which is not a fair
comparison to EAGER. Therefore we will re-run but allowing multi-threading for
bwa, one of the longest running steps.

```bash
mkdir -p ~/benchmarks/output/paleomix_optimised
cp ~/benchmarks/output/paleomix/makefile_paleomix.yaml ~/benchmarks/output/paleomix_optimised/makefile_paleomix.yaml
```

For comparison with nf-core/eager, that also allows non-sequential job running,
we will set this value to the same number of CPUs which is 4.

```bash
screen -R paleomix_optimised
conda activate paleomix
cd ~/benchmarks/output/paleomix_optimised

for i in {1..10}; do
    { time paleomix bam_pipeline run makefile_paleomix.yaml --max-threads 4 ; } 2> ../time_paleomix_optimised_"$i".log
    if [[ $i != 10 ]]; then
         rm -r ~/benchmarks/output/paleomix_optimised/!(makefile_paleomix.yaml)
         rm ~/benchmarks/reference/paleomix/*.{dict,amb,ann,bwt,fai,pac,sa,validated}

    fi
done
```

### nf-core/eager setup

Go into directory and set up file

```bash
mkdir ~/benchmarks/output/nfcore-eager
cd !$

nano nfcore-eager_tsv.tsv
```

Paste the following (ensure you're pasting TABS and not spaces) then save

```text
Sample_Name Library_ID  Lane  Colour_Chemistry  SeqType Organism  Strandedness  UDG_Treatment R1  R2  BAM
COD076  COD076E1bL1 1 4 PE  g_morhua  double  none  /home/cloud/benchmarks/input/COD076/COD076E1bL1_S0_L001_R1_000.fastq.gz /home/cloud/benchmarks/input/COD076/COD076E1bL1_S0_L001_R2_000.fastq.gz NA
COD076  COD076E1bL1 6 4 PE  g_morhua  double  none  /home/cloud/benchmarks/input/COD076/COD076E1bL1_S0_L006_R1_000.fastq.gz /home/cloud/benchmarks/input/COD076/COD076E1bL1_S0_L006_R2_000.fastq.gz NA
COD076  COD076E1bL1 8 4 PE  g_morhua  double  none  /home/cloud/benchmarks/input/COD076/COD076E1bL1_S0_L008_R1_000.fastq.gz /home/cloud/benchmarks/input/COD076/COD076E1bL1_S0_L008_R2_000.fastq.gz NA
COD092  COD092E1bL1i69  6 4 PE  g_morhua  double  none  /home/cloud/benchmarks/input/COD092/COD092E1bL1i69_S0_L006_R1_000.fastq.gz  /home/cloud/benchmarks/input/COD092/COD092E1bL1i69_S0_L006_R2_000.fastq.gz  NA
COD092  COD092E1bL1i69  7 4 PE  g_morhua  double  none  /home/cloud/benchmarks/input/COD092/COD092E1bL1i69_S0_L007_R1_000.fastq.gz  /home/cloud/benchmarks/input/COD092/COD092E1bL1i69_S0_L007_R2_000.fastq.gz  NA
COD092  COD092E1bL1i69  8 4 PE  g_morhua  double  none  /home/cloud/benchmarks/input/COD092/COD092E1bL1i69_S0_L008_R1_000.fastq.gz  /home/cloud/benchmarks/input/COD092/COD092E1bL1i69_S0_L008_R2_000.fastq.gz  NA
```

To then run

```bash

for i in {1..10}; do
    { time nextflow run nf-core/eager -r dev \
      --input 'nfcore-eager_tsv.tsv' \
      -c ~/.nextflow/pub_eager_vikingfish.conf \
      -profile pub_eager_vikingfish,benchmarking_vikingfish,singularity \
            --fasta '/home/cloud/benchmarks/reference/GCF_902167405.1_gadMor3.0_genomic.fasta' \
            -name 'gwdg_test' \
            --outdir ~/benchmarks/output/nfcore-eager/results/ \
            -w ~/benchmarks/output/nfcore-eager/work/ \
            --skip_fastqc \
            --skip_preseq \
            --run_bam_filtering \
            --bam_mapping_quality_threshold 25 \
            --bam_discard_unmapped \
            --bam_unmapped_type 'discard' \
            --dedupper 'markduplicates' } 2> ../time_nf-core_eager_"$i".log
    if [[ $i != 10 ]]; then
         rm -r ~/benchmarks/output/nfcore-eager/!(nfcore-eager_tsv.tsv) ~/benchmarks/output/nfcore-eager/.nex*
    fi
done
```

### nf-core/eager optimised

The nf-core/eager optimised version (to do fair comparison between optimised
paleomix with more realistic bwa multi-threading), is the same as default
nf-core/eager but with different output directory.

```bash
cp -r ~/benchmarks/output/nfcore-eager/ ~/benchmarks/output/nfcore-eager-optimised/
```

### Final Benchmarking Command

To ensure as fair as possible comparison, rather than running each program
separately, we will run each iteration of each command next to each other.

Therefore if IO gets busy on other VMs in the network, each pipeline run of
that iteration will fall approximately in the same period.

```bash
cd ~/benchmarks/output

mkdir runtimes results

for i in {1..10}; do
    ## EAGER
    unset DISPLAY && { time singularity exec -B ~/benchmarks/output/EAGER:/data ~/.singularity/cache/EAGER-cache/EAGER-GUI_latest.sif eagercli /data ; } 2> runtimes/time_EAGER_"$i".log
    cp ~/benchmarks/output/EAGER/Report_EAGER.csv results/Report_EAGER_$i.csv
    if [[ $i != 10 ]]; then
         rm -r ~/benchmarks/output/EAGER/*/*/ ~/benchmarks/output/EAGER/*.{csv,html,png,ReportGenerator,txt} ~/benchmarks/output/EAGER/*/*.log ~/benchmarks/output/EAGER/*/DONE*
         rm ~/benchmarks/reference/EAGER/*.{dict,amb,ann,bwt,fai,pac,sa} ~/benchmarks/reference/EAGER/DONE*
    fi
    ## Paleomix Default
    conda activate paleomix
    { time paleomix bam_pipeline run ~/benchmarks/output/paleomix/makefile_paleomix.yaml ; } 2> runtimes/time_paleomix_"$i".log
    rename -e "s/.summary/_$i.summary/g;s/COD/paleomix_COD/" paleomix/*summary
    cp paleomix/*summary results/
    if [[ $i != 10 ]]; then
         rm -r ~/benchmarks/output/paleomix/!(makefile_paleomix.yaml)
         rm ~/benchmarks/reference/paleomix/*.{dict,amb,ann,bwt,fai,pac,sa,validated}
    fi
    ## Paleomix Better defaults=
    { time paleomix bam_pipeline run ~/benchmarks/output/paleomix_optimised/makefile_paleomix.yaml --bwa-max-threads 4 ; } 2> runtimes/time_paleomix_optimised_"$i".log
    rename -e "s/.summary/_$i.summary/g;s/COD/paleomix_optimised_COD/" paleomix-optimised/*summary
    if [[ $i != 10 ]]; then
         rm -r ~/benchmarks/output/paleomix_optimised/!(makefile_paleomix.yaml)
         rm ~/benchmarks/reference/paleomix/*.{dict,amb,ann,bwt,fai,pac,sa,validated}
    fi
    conda deactivate
    ## nf-core/eager
    cd ~/benchmarks/output/nfcore-eager/
    { time nextflow run nf-core/eager -r dev --input ~/benchmarks/output/nfcore-eager/nfcore-eager_tsv.tsv -c ~/.nextflow/pub_eager_vikingfish.conf -profile pub_eager_vikingfish,singularity --fasta ~/benchmarks/reference/GCF_902167405.1_gadMor3.0_genomic.fasta --outdir ~/benchmarks/output/nfcore-eager/results/ -w ~/benchmarks/output/nfcore-eager/work/ --skip_fastqc --skip_preseq --run_bam_filtering --bam_mapping_quality_threshold 25 --bam_discard_unmapped --bam_unmapped_type 'discard' --dedupper 'markduplicates' ; } 2> ../runtimes/time_nf-core-eager_"$i".log
    cd ~/benchmarks/output/
    cp ~/benchmarks/output/nfcore-eager/results/multiqc/multiqc_data/multiqc_general_stats.txt results/nfcore-eager_multiqc_general_stats_$i.csv
    if [[ $i != 10 ]]; then
         rm -r ~/benchmarks/output/nfcore-eager/!(nfcore-eager_tsv.tsv) ~/benchmarks/output/nfcore-eager/.nex*
    fi
    ## nf-core/eager optimised
    cd ~/benchmarks/output/nfcore-eager-optimised/
    { time nextflow run nf-core/eager -r dev --input ~/benchmarks/output/nfcore-eager-optimised/nfcore-eager_tsv.tsv -c ~/.nextflow/pub_eager_vikingfish.conf -profile pub_eager_vikingfish_optimised,pub_eager_vikingfish,singularity --fasta ~/benchmarks/reference/GCF_902167405.1_gadMor3.0_genomic.fasta --outdir ~/benchmarks/output/nfcore-eager-optimised/results/ -w ~/benchmarks/output/nfcore-eager-optimised/work/ --skip_fastqc --skip_preseq --run_bam_filtering --bam_mapping_quality_threshold 25 --bam_discard_unmapped --bam_unmapped_type 'discard' --dedupper 'markduplicates' ; } 2> ../runtimes/time_nf-core-eager-optimised_"$i".log
    cd ~/benchmarks/output/
    if [[ $i != 10 ]]; then
         rm -r ~/benchmarks/output/nfcore-eager-optimised/!(nfcore-eager_tsv.tsv) ~/benchmarks/output/nfcore-eager-optimised/.nex*
    fi
done
```

### Results Clean-Up

To gather the results of the runtimes in a single place, we can run the following
command.

```bash
grep -n -e 'real' -e 'sys' -e 'user' runtimes/*.log > benchmarking_aggregated_runtimes.txt
```

The aggregated runtime and pipeline 'report' files was then downloaded to a
local PC (via scp, in this directory under `results`), and we summarised in the
results with R.

The R environment is as follows

```r
library(tidyverse)
library(knitr)

sessionInfo()

R version 3.6.3 (2020-02-29)
Platform: x86_64-pc-linux-gnu (64-bit)
Running under: Ubuntu 20.04.1 LTS

Matrix products: default
BLAS:   /usr/lib/x86_64-linux-gnu/blas/libblas.so.3.9.0
LAPACK: /usr/lib/x86_64-linux-gnu/lapack/liblapack.so.3.9.0

locale:
 [1] LC_CTYPE=en_GB.UTF-8       LC_NUMERIC=C               LC_TIME=en_GB.UTF-8        LC_COLLATE=en_GB.UTF-8     LC_MONETARY=en_GB.UTF-8
 [6] LC_MESSAGES=en_GB.UTF-8    LC_PAPER=en_GB.UTF-8       LC_NAME=C                  LC_ADDRESS=C               LC_TELEPHONE=C
[11] LC_MEASUREMENT=en_GB.UTF-8 LC_IDENTIFICATION=C

attached base packages:
[1] stats     graphics  grDevices utils     datasets  methods   base

other attached packages:
 [1] knitr_1.29      forcats_0.5.0   stringr_1.4.0   dplyr_1.0.1     purrr_0.3.4     readr_1.3.1     tidyr_1.1.1     tibble_3.0.3    ggplot2_3.3.2  
[10] tidyverse_1.3.0

loaded via a namespace (and not attached):
 [1] Rcpp_1.0.5       highr_0.8        cellranger_1.1.0 pillar_1.4.6     compiler_3.6.3   dbplyr_1.4.4     tools_3.6.3      evaluate_0.14    jsonlite_1.7.0  
[10] lubridate_1.7.9  lifecycle_0.2.0  gtable_0.3.0     pkgconfig_2.0.3  rlang_0.4.7      reprex_0.3.0     cli_2.0.2        DBI_1.1.0        rstudioapi_0.11
[19] xfun_0.16        haven_2.3.1      withr_2.2.0      xml2_1.3.2       httr_1.4.2       fs_1.5.0         generics_0.0.2   vctrs_0.3.2      hms_0.5.3
[28] grid_3.6.3       tidyselect_1.1.0 glue_1.4.1       R6_2.4.1         fansi_0.4.1      readxl_1.3.1     modelr_0.1.8     blob_1.2.1       magrittr_1.5
[37] backports_1.1.8  scales_1.1.1     ellipsis_0.3.1   rvest_0.3.6      assertthat_0.2.1 colorspace_1.4-1 utf8_1.1.4       stringi_1.4.6    munsell_0.5.0
[46] broom_0.7.0      crayon_1.3.4
```

And the two R markdown scripts (`benchmarking_runtime_parsing.Rmd` and
`benchmarking_results_parsing.Rmd` in this directory) were used for generating
summary statistics and plots.
