## Supplemental file archive of walkthrough and results files for "Reproducible, portable, and efficient ancient genome reconstruction with nf-core/eager": paleomix_environment.yml

```
name: paleomix
channels:
    - defaults
    - bioconda
    - conda-forge
dependencies:
  - python=2.7 
  - pip=20.1.1
  - adapterremoval=2.3.1 
  - samtools=1.9 
  - picard=2.22.9 
  - bowtie2=2.3.5.1 
  - bwa=0.7.17
  - mapdamage2=2.0.9 
  - gatk=3.8 
  - r-base=3.5.1 
  - r-rcpp=1.0.4.6 
  - r-rcppgsl=0.3.7 
  - r-gam=1.16.1 
  - r-inline=0.3.15
```
