## Supplemental file archive of walkthrough and results files for "Reproducible, portable, and efficient ancient genome reconstruction with nf-core/eager": barquera2020_sdag_multiqc_1_9_report.html

MultiQC Report


### Toggle navigation v1.9

Loading report..

- General Stats
- FastQC (pre-AdapterRemoval)
  - Sequence Counts
  - Sequence Quality Histograms
  - Per Sequence Quality Scores
  - Per Base Sequence Content
  - Per Sequence GC Content
  - Per Base N Content
  - Sequence Length Distribution
  - Sequence Duplication Levels
  - Overrepresented sequences
  - Adapter Content
  - Status Checks
- Adapter Removal
  - Retained and Discarded Paired-End Collapsed
  - Length Distribution Paired End Collapsed
- FastQC (post-AdapterRemoval)
  - Sequence Counts
  - Sequence Quality Histograms
  - Per Sequence Quality Scores
  - Per Base Sequence Content
  - Per Sequence GC Content
  - Per Base N Content
  - Sequence Length Distribution
  - Sequence Duplication Levels
  - Overrepresented sequences
  - Adapter Content
  - Status Checks
- MALT
  - Metagenomic Mappability
  - Taxonomic assignment success
- Samtools Flagstat (pre-samtools filter)
- Samtools Flagstat (post-samtools filter)
- Picard
- Preseq
- DamageProfiler
  - 3P misincorporation plot
  - 5P misincorporation plot
  - Forward read length distribution
  - Reverse read length distribution
- QualiMap
  - Coverage histogram
  - Cumulative genome coverage
  - GC content distribution
- SexDetErrmine
  - Relative Coverage
  - Read Counts
- nf-core/eager Software Versions
- nf-core/eager Workflow Summary

Toolbox

##### MultiQC Toolbox

###### Apply Highlight Samples

This report has flat image plots that won't be highlighted.

+

Regex mode off
help
 Clear

###### Apply Rename Samples

This report has flat image plots that won't be renamed.  
See the documentation
for help.

+

Click here for bulk input.

Paste two columns of a tab-delimited table here (eg. from Excel).

First column should be the old name, second column the new name.

Add

Regex mode off
help
 Clear

###### Apply Show / Hide Samples

This report has flat image plots that won't be hidden.  
See the documentation
for help.

Hide matching samples

Show only matching samples

+

Warning! This can take a few seconds.

Regex mode off
help
 Clear

###### Export Plots

- Images
- Data

px

px

Aspect ratio

PNG
JPEG
SVG

Plot scaling

X

Download the raw data used to create the plots in this report below:

Format:

Tab-separated
Comma-separated
JSON

Note that additional data was saved in `multiqc_data` when this report was generated.

---

###### Choose Plots

 All
 None


---


   Download Plot Images

If you use plots from MultiQC in a publication or presentation, please cite:

> **MultiQC: Summarize analysis results for multiple tools and samples in a single report**  
> *Philip Ewels, Måns Magnusson, Sverker Lundin and Max Käller*  
> Bioinformatics (2016)  
> doi: 10.1093/bioinformatics/btw354  
> PMID: 27312411

###### Save Settings

You can save the toolbox settings for this report to the browser.

 Save


---

###### Load Settings

Choose a saved report profile from the dropdown box below:

[ select ]

Load
 Delete
 Set default
 Clear default

###### About MultiQC

This report was generated using MultiQC, version 1.9

You can see a YouTube video describing how to use MultiQC reports here:
https://youtu.be/qPbIlO\_KWN0

For more information about MultiQC, including other videos and
extensive documentation, please visit http://multiqc.info

You can report bugs, suggest improvements and find the source code for MultiQC on GitHub:
https://github.com/ewels/MultiQC

MultiQC is published in Bioinformatics:

> **MultiQC: Summarize analysis results for multiple tools and samples in a single report**  
> *Philip Ewels, Måns Magnusson, Sverker Lundin and Max Käller*  
> Bioinformatics (2016)  
> doi: 10.1093/bioinformatics/btw354  
> PMID: 27312411

A modular tool to aggregate results from bioinformatics analyses across many samples into a single report.

> This report has been generated by the nf-core/eager analysis pipeline. For information about how to interpret these results, please see the documentation.

###### JavaScript Disabled

MultiQC reports use JavaScript for plots and toolbox functions. It looks like
you have JavaScript disabled in your web browser. Please note that many of the report
functions will not work as intended.

Loading report..

Report
generated on 2020-10-05, 02:19
based on data in:

- `/projects1/clusterhomes/fellows/.nextflow/assets/nf-core/eager/assets/multiqc_config.yaml`
- `/projects1/users/fellows/nextflow/eager2/publication/benchmarking_pathogen/work/a0/b2cbe00df48f1846a7925baf78a7f6`

---

×
don't show again

**Welcome!** Not sure where to start?  
Watch a tutorial video
  *(6:06)*

#### General Statistics

 Copy table

 Configure Columns

 Sort by highlight

 Plot
Showing ^36^/_36_ rows and ^33^/_61_ columns.

| Sample Name | Seqs | Length | % GC | % Trimmed | Seqs | Length | % GC | Reads | Mapped | % Metagenomic Mapped | Tax assigned | % Tax assigned | Reads | Reads Mapped | Reads | Endogenous DNA (%) | Reads Mapped | Endogenous DNA Post (%) | % Dups | 5 Prime C>T 1st base | 5 Prime C>T 2nd base | 3 Prime G>A 1st base | 3 Prime G>A 2nd base | Mean read length | Median read length | MT genome reads | MT genome coverage | MT to Nuclear Ratio | Aligned | Mean cov | Median cov | ≥ 1X | ≥ 2X | ≥ 3X | ≥ 4X | ≥ 5X | % GC | % Dups | % Failed | Reads Trimmed | % Dups | % Failed | Read length std. dev. | Genome coverage | Genome reads | % Aligned | Total reads | Error rate | Err Rate X | Err Rate Y | Rate X | Rate Y | Number of SNPs | Contamination Estimate (Method1\_MOM) | Estimate Error (Method1\_MOM) | Contamination Estimate (Method1\_ML) | Estimate Error (Method1\_ML) | Contamination Estimate (Method2\_MOM) | Estimate Error (Method2\_MOM) | Contamination Estimate (Method2\_ML) | Estimate Error (Method2\_ML) |
| --- | --- | --- | --- | --- | --- | --- | --- | --- | --- | --- | --- | --- | --- | --- | --- | --- | --- | --- | --- | --- | --- | --- | --- | --- | --- | --- | --- | --- | --- | --- | --- | --- | --- | --- | --- | --- | --- | --- | --- | --- | --- | --- | --- | --- | --- | --- | --- | --- | --- | --- | --- | --- | --- | --- | --- | --- | --- | --- | --- | --- | --- |
| ERR4065478 | 7,392,875 | 62 bp | 63% | 12.7% | 7,369,645 | 62 bp | 63% |  |  |  |  |  |  |  |  |  |  |  |  |  |  |  |  |  |  |  |  |  |  |  |  |  |  |  |  |  |  | 6.3% | 18% | 940,363 | 6.2% | 18% |  |  |  |  |  |  |  |  |  |  |  |  |  |  |  |  |  |  |  |
| ERR4065479 | 7,416,023 | 63 bp | 62% | 12.8% | 7,397,995 | 63 bp | 62% |  |  |  |  |  |  |  |  |  |  |  |  |  |  |  |  |  |  |  |  |  |  |  |  |  |  |  |  |  |  | 6.0% | 18% | 946,466 | 6.0% | 18% |  |  |  |  |  |  |  |  |  |  |  |  |  |  |  |  |  |  |  |
| ERR4065480 | 6,217,740 | 59 bp | 58% | 14.7% | 6,189,668 | 59 bp | 58% |  |  |  |  |  |  |  |  |  |  |  |  |  |  |  |  |  |  |  |  |  |  |  |  |  |  |  |  |  |  | 6.9% | 18% | 911,344 | 6.9% | 18% |  |  |  |  |  |  |  |  |  |  |  |  |  |  |  |  |  |  |  |
| ERR4065481 | 8,814,520 | 61 bp | 61% | 13.4% | 8,787,931 | 61 bp | 61% |  |  |  |  |  |  |  |  |  |  |  |  |  |  |  |  |  |  |  |  |  |  |  |  |  |  |  |  |  |  | 6.2% | 18% | 1,184,176 | 6.2% | 18% |  |  |  |  |  |  |  |  |  |  |  |  |  |  |  |  |  |  |  |
| ERR4065482 | 6,746,607 | 65 bp | 58% | 13.4% | 6,730,715 | 65 bp | 58% |  |  |  |  |  |  |  |  |  |  |  |  |  |  |  |  |  |  |  |  |  |  |  |  |  |  |  |  |  |  | 8.0% | 18% | 904,045 | 7.9% | 18% |  |  |  |  |  |  |  |  |  |  |  |  |  |  |  |  |  |  |  |
| ERR4065483 | 8,925,991 | 65 bp | 58% | 13.5% | 8,910,925 | 65 bp | 58% |  |  |  |  |  |  |  |  |  |  |  |  |  |  |  |  |  |  |  |  |  |  |  |  |  |  |  |  |  |  | 6.8% | 18% | 1,205,996 | 6.7% | 18% |  |  |  |  |  |  |  |  |  |  |  |  |  |  |  |  |  |  |  |
| ERR4065497\_1 | 2,835,130 | 47 bp | 63% | 84.0% | 3,487,129 | 53 bp | 63% |  |  |  |  |  |  |  |  |  |  |  |  |  |  |  |  |  |  |  |  |  |  |  |  |  |  |  |  |  |  | 5.3% | 18% | 4,765,196 | 4.5% | 9% |  |  |  |  |  |  |  |  |  |  |  |  |  |  |  |  |  |  |  |
| ERR4065497\_2 | 2,835,130 | 47 bp | 63% |  |  |  |  |  |  |  |  |  |  |  |  |  |  |  |  |  |  |  |  |  |  |  |  |  |  |  |  |  |  |  |  |  |  | 5.0% | 18% |  |  |  |  |  |  |  |  |  |  |  |  |  |  |  |  |  |  |  |  |  |  |
| ERR4065498\_1 | 2,522,025 | 48 bp | 62% | 83.5% | 3,126,489 | 53 bp | 62% |  |  |  |  |  |  |  |  |  |  |  |  |  |  |  |  |  |  |  |  |  |  |  |  |  |  |  |  |  |  | 5.0% | 18% | 4,212,510 | 4.6% | 9% |  |  |  |  |  |  |  |  |  |  |  |  |  |  |  |  |  |  |  |
| ERR4065498\_2 | 2,522,025 | 48 bp | 62% |  |  |  |  |  |  |  |  |  |  |  |  |  |  |  |  |  |  |  |  |  |  |  |  |  |  |  |  |  |  |  |  |  |  | 4.8% | 18% |  |  |  |  |  |  |  |  |  |  |  |  |  |  |  |  |  |  |  |  |  |  |
| ERR4065499\_1 | 3,032,873 | 46 bp | 59% | 90.0% | 3,485,321 | 51 bp | 59% |  |  |  |  |  |  |  |  |  |  |  |  |  |  |  |  |  |  |  |  |  |  |  |  |  |  |  |  |  |  | 5.5% | 18% | 5,458,318 | 5.2% | 9% |  |  |  |  |  |  |  |  |  |  |  |  |  |  |  |  |  |  |  |
| ERR4065499\_2 | 3,032,873 | 46 bp | 59% |  |  |  |  |  |  |  |  |  |  |  |  |  |  |  |  |  |  |  |  |  |  |  |  |  |  |  |  |  |  |  |  |  |  | 5.3% | 18% |  |  |  |  |  |  |  |  |  |  |  |  |  |  |  |  |  |  |  |  |  |  |
| ERR4065500\_1 | 3,542,865 | 47 bp | 61% | 86.9% | 4,227,187 | 52 bp | 61% |  |  |  |  |  |  |  |  |  |  |  |  |  |  |  |  |  |  |  |  |  |  |  |  |  |  |  |  |  |  | 5.3% | 18% | 6,155,568 | 5.0% | 9% |  |  |  |  |  |  |  |  |  |  |  |  |  |  |  |  |  |  |  |
| ERR4065500\_2 | 3,542,865 | 47 bp | 61% |  |  |  |  |  |  |  |  |  |  |  |  |  |  |  |  |  |  |  |  |  |  |  |  |  |  |  |  |  |  |  |  |  |  | 5.1% | 18% |  |  |  |  |  |  |  |  |  |  |  |  |  |  |  |  |  |  |  |  |  |  |
| ERR4065501\_1 | 4,239,907 | 48 bp | 58% | 81.3% | 5,400,980 | 53 bp | 57% |  |  |  |  |  |  |  |  |  |  |  |  |  |  |  |  |  |  |  |  |  |  |  |  |  |  |  |  |  |  | 6.8% | 18% | 6,895,518 | 4.9% | 9% |  |  |  |  |  |  |  |  |  |  |  |  |  |  |  |  |  |  |  |
| ERR4065501\_2 | 4,239,907 | 48 bp | 58% |  |  |  |  |  |  |  |  |  |  |  |  |  |  |  |  |  |  |  |  |  |  |  |  |  |  |  |  |  |  |  |  |  |  | 6.5% | 18% |  |  |  |  |  |  |  |  |  |  |  |  |  |  |  |  |  |  |  |  |  |  |
| ERR4065502\_1 | 3,767,036 | 48 bp | 58% | 81.0% | 4,817,493 | 54 bp | 57% |  |  |  |  |  |  |  |  |  |  |  |  |  |  |  |  |  |  |  |  |  |  |  |  |  |  |  |  |  |  | 5.9% | 18% | 6,100,560 | 4.9% | 9% |  |  |  |  |  |  |  |  |  |  |  |  |  |  |  |  |  |  |  |
| ERR4065502\_2 | 3,767,036 | 48 bp | 58% |  |  |  |  |  |  |  |  |  |  |  |  |  |  |  |  |  |  |  |  |  |  |  |  |  |  |  |  |  |  |  |  |  |  | 5.6% | 18% |  |  |  |  |  |  |  |  |  |  |  |  |  |  |  |  |  |  |  |  |  |  |
| SJN001.A |  |  |  |  |  |  |  |  |  |  |  |  |  |  |  |  |  |  |  |  |  |  |  |  |  |  |  |  | 374,922 | 0.0X | 0.0X | 0.6% | 0.0% | 0.0% | 0.0% | 0.0% | 47% |  |  |  |  |  |  |  |  | 100.0% | 374,922 | 1.12% | 0.0 | 0.0 | 0.4 | 0.3 |  |  |  |  |  |  |  |  |  |
| SJN001.A0101 |  |  |  |  |  |  |  | 7,099,701 | 298,761 | 4.2% | 298,004 | 99.7% | 7,369,645 | 269,944 | 269,944 | 3.66 | 269,944 | 3.66 | 7.3% | 18.8% | 10.0% | 14.8% | 7.0% | 54.92bp | 53.00bp | 2,263 | 8.8 X | 2,034.7 |  |  |  |  |  |  |  |  |  |  |  |  |  |  | 15.03bp | 0.0 X | 247,972 |  |  |  |  |  |  |  | 1.0 | 0.0 | N/A | 0.0 | 0.0 | 0.0 | N/A | 0.0 | 0.0 |
| SJN001.A0102 |  |  |  |  |  |  |  | 3,353,804 | 197,712 | 5.9% | 197,094 | 99.7% | 3,487,129 | 133,325 | 133,325 | 3.82 | 133,325 | 3.82 | 6.5% | 7.6% | 1.2% | 6.6% | 1.3% | 50.15bp | 50.00bp | 1,230 | 4.1 X | 2,055.4 |  |  |  |  |  |  |  |  |  |  |  |  |  |  | 13.29bp | 0.0 X | 123,457 |  |  |  |  |  |  |  | 0.0 | N/A | N/A | 1.3 | N/A | N/A | N/A | 1.3 | N/A |
| SJN001.B |  |  |  |  |  |  |  |  |  |  |  |  |  |  |  |  |  |  |  |  |  |  |  |  |  |  |  |  | 177,348 | 0.0X | 0.0X | 0.3% | 0.0% | 0.0% | 0.0% | 0.0% | 45% |  |  |  |  |  |  |  |  | 100.0% | 177,348 | 0.82% | 0.0 | 0.1 | 0.4 | 0.3 |  |  |  |  |  |  |  |  |  |
| SJN001.B0101 |  |  |  |  |  |  |  | 7,267,202 | 241,567 | 3.3% | 240,779 | 99.7% | 7,397,995 | 130,793 | 130,793 | 1.77 | 130,793 | 1.77 | 9.2% | 13.6% | 7.2% | 8.4% | 4.1% | 65.40bp | 75.00bp | 726 | 2.9 X | 1,164.0 |  |  |  |  |  |  |  |  |  |  |  |  |  |  | 13.88bp | 0.0 X | 118,095 |  |  |  |  |  |  |  | 0.0 | N/A | N/A | 1.3 | N/A | N/A | N/A | 1.3 | N/A |
| SJN001.B0102 |  |  |  |  |  |  |  | 3,063,321 | 143,768 | 4.7% | 143,246 | 99.6% | 3,126,489 | 63,168 | 63,168 | 2.02 | 63,168 | 2.02 | 7.3% | 4.3% | 0.7% | 3.1% | 0.8% | 53.77bp | 51.00bp | 358 | 1.2 X | 1,171.5 |  |  |  |  |  |  |  |  |  |  |  |  |  |  | 12.06bp | 0.0 X | 58,169 |  |  |  |  |  |  |  | 0.0 | N/A | N/A | 1.3 | N/A | N/A | N/A | 1.3 | N/A |
| SJN002.A |  |  |  |  |  |  |  |  |  |  |  |  |  |  |  |  |  |  |  |  |  |  |  |  |  |  |  |  | 1,554,667 | 0.0X | 0.0X | 2.5% | 0.1% | 0.0% | 0.0% | 0.0% | 49% |  |  |  |  |  |  |  |  | 100.0% | 1,554,667 | 1.19% | 0.0 | 0.0 | 0.4 | 0.4 |  |  |  |  |  |  |  |  |  |
| SJN002.A0101 |  |  |  |  |  |  |  | 5,093,709 | 553,627 | 10.9% | 552,507 | 99.8% | 6,189,668 | 1,095,959 | 1,095,959 | 17.71 | 1,095,959 | 17.71 | 9.9% | 20.6% | 11.2% | 15.5% | 7.9% | 55.39bp | 53.00bp | 15,816 | 64.0 X | 3,741.5 |  |  |  |  |  |  |  |  |  |  |  |  |  |  | 15.26bp | 0.0 X | 971,504 |  |  |  |  |  |  |  | 5.0 | -0.0 | 0.0 | -0.0 | 0.0 | -0.0 | 0.1 | -0.0 | 0.0 |
| SJN002.A0102 |  |  |  |  |  |  |  | 2,866,859 | 352,929 | 12.3% | 351,966 | 99.7% | 3,485,321 | 618,462 | 618,462 | 17.74 | 618,462 | 17.74 | 8.3% | 8.0% | 1.2% | 7.1% | 1.2% | 49.74bp | 49.00bp | 11,566 | 39.8 X | 4,526.6 |  |  |  |  |  |  |  |  |  |  |  |  |  |  | 13.36bp | 0.0 X | 555,781 |  |  |  |  |  |  |  | 0.0 | N/A | N/A | 1.3 | N/A | N/A | N/A | 1.3 | N/A |
| SJN002.B |  |  |  |  |  |  |  |  |  |  |  |  |  |  |  |  |  |  |  |  |  |  |  |  |  |  |  |  | 359,232 | 0.0X | 0.0X | 0.6% | 0.0% | 0.0% | 0.0% | 0.0% | 52% |  |  |  |  |  |  |  |  | 100.0% | 359,232 | 1.16% | 0.0 | 0.0 | 0.3 | 0.4 |  |  |  |  |  |  |  |  |  |
| SJN002.B0101 |  |  |  |  |  |  |  | 8,524,971 | 438,874 | 5.1% | 438,110 | 99.8% | 8,787,931 | 262,960 | 262,960 | 2.99 | 262,960 | 2.99 | 7.9% | 19.2% | 9.8% | 14.6% | 7.0% | 55.87bp | 54.00bp | 1,271 | 4.8 X | 1,109.3 |  |  |  |  |  |  |  |  |  |  |  |  |  |  | 14.53bp | 0.0 X | 240,912 |  |  |  |  |  |  |  | 0.0 | N/A | N/A | 1.3 | N/A | N/A | N/A | 1.3 | N/A |
| SJN002.B0102 |  |  |  |  |  |  |  | 4,101,402 | 257,605 | 6.3% | 257,108 | 99.8% | 4,227,187 | 125,785 | 125,785 | 2.98 | 125,785 | 2.98 | 6.9% | 8.8% | 1.5% | 7.7% | 1.4% | 50.55bp | 50.00bp | 617 | 2.0 X | 1,046.7 |  |  |  |  |  |  |  |  |  |  |  |  |  |  | 13.24bp | 0.0 X | 116,432 |  |  |  |  |  |  |  | 0.0 | N/A | N/A | 1.3 | N/A | N/A | N/A | 1.3 | N/A |
| SJN003.A |  |  |  |  |  |  |  |  |  |  |  |  |  |  |  |  |  |  |  |  |  |  |  |  |  |  |  |  | 291,640 | 0.0X | 0.0X | 0.5% | 0.0% | 0.0% | 0.0% | 0.0% | 46% |  |  |  |  |  |  |  |  | 100.0% | 291,640 | 1.01% | 0.0 | 0.0 | 0.5 | 0.4 |  |  |  |  |  |  |  |  |  |
| SJN003.A0101 |  |  |  |  |  |  |  | 6,562,267 | 184,750 | 2.8% | 184,291 | 99.8% | 6,730,715 | 168,448 | 168,448 | 2.50 | 168,448 | 2.50 | 10.3% | 20.9% | 11.7% | 12.5% | 6.4% | 64.23bp | 74.00bp | 5,399 | 21.9 X | 7,343.9 |  |  |  |  |  |  |  |  |  |  |  |  |  |  | 14.27bp | 0.0 X | 145,631 |  |  |  |  |  |  |  | 0.0 | N/A | N/A | 1.3 | N/A | N/A | N/A | 1.3 | N/A |
| SJN003.A0102 |  |  |  |  |  |  |  | 5,247,729 | 186,396 | 3.6% | 185,847 | 99.7% | 5,400,980 | 153,251 | 153,251 | 2.84 | 153,251 | 2.84 | 8.2% | 7.3% | 1.1% | 4.8% | 0.9% | 53.96bp | 51.00bp | 5,088 | 17.2 X | 7,403.0 |  |  |  |  |  |  |  |  |  |  |  |  |  |  | 13.27bp | 0.0 X | 135,522 |  |  |  |  |  |  |  | 0.0 | N/A | N/A | 1.3 | N/A | N/A | N/A | 1.3 | N/A |
| SJN003.B |  |  |  |  |  |  |  |  |  |  |  |  |  |  |  |  |  |  |  |  |  |  |  |  |  |  |  |  | 1,978,323 | 0.0X | 0.0X | 3.6% | 0.1% | 0.0% | 0.0% | 0.0% | 46% |  |  |  |  |  |  |  |  | 100.0% | 1,978,323 | 1.13% | 0.0 | 0.0 | 0.4 | 0.4 |  |  |  |  |  |  |  |  |  |
| SJN003.B0101 |  |  |  |  |  |  |  | 7,578,321 | 251,066 | 3.3% | 250,630 | 99.8% | 8,910,925 | 1,332,604 | 1,332,604 | 14.95 | 1,332,604 | 14.95 | 9.3% | 20.7% | 11.1% | 10.7% | 5.8% | 66.08bp | 75.00bp | 1,319 | 5.3 X | 209.0 |  |  |  |  |  |  |  |  |  |  |  |  |  |  | 13.19bp | 0.0 X | 1,207,254 |  |  |  |  |  |  |  | 3.0 | -0.1 | 0.2 | -0.0 | 0.0 | -0.1 | 0.0 | 0.0 | 0.0 |
| SJN003.B0102 |  |  |  |  |  |  |  | 3,984,911 | 135,197 | 3.4% | 134,872 | 99.8% | 4,817,493 | 832,582 | 832,582 | 17.28 | 832,582 | 17.28 | 7.5% | 7.0% | 1.2% | 4.5% | 1.0% | 54.54bp | 51.00bp | 844 | 2.8 X | 210.3 |  |  |  |  |  |  |  |  |  |  |  |  |  |  | 12.87bp | 0.0 X | 768,906 |  |  |  |  |  |  |  | 2.0 | 0.0 | 0.0 | 0.0 | 0.0 | 0.0 | 0.0 | 0.0 | 0.0 |

×

###### General Statistics: Columns

Uncheck the tick box to hide columns. Click and drag the handle on the left to change order.

Show All
Show None

| Sort | Visible | Group | Column | Description | ID | Scale |
| --- | --- | --- | --- | --- | --- | --- |
| \|\| |  | FastQC (pre-AdapterRemoval) | Seqs | Total Sequences () | `total_sequences` | read\_count |
| \|\| |  | FastQC (pre-AdapterRemoval) | Length | Average Sequence Length (bp) | `avg_sequence_length` | None |
| \|\| |  | FastQC (pre-AdapterRemoval) | % GC | Average % GC Content | `percent_gc` | None |
| \|\| |  | Adapter Removal | % Trimmed | % trimmed reads | `percent_aligned` | percent\_aligned |
| \|\| |  | FastQC (post-AdapterRemoval) | Seqs | Total Sequences () | `total_sequences` | read\_count |
| \|\| |  | FastQC (post-AdapterRemoval) | Length | Average Sequence Length (bp) | `avg_sequence_length` | None |
| \|\| |  | FastQC (post-AdapterRemoval) | % GC | Average % GC Content | `percent_gc` | None |
| \|\| |  | MALT | Reads | Number of reads in sample () | `Num. of queries` | read\_count |
| \|\| |  | MALT | Mapped | Number of mapped reads () | `Total reads` | read\_count |
| \|\| |  | MALT | % Metagenomic Mapped | Percentage of mapped reads | `Mappability` | None |
| \|\| |  | MALT | Tax assigned | Number of reads assigned to a Taxonomic node () | `Assig. Taxonomy` | read\_count |
| \|\| |  | MALT | % Tax assigned | Percentage of mapped reads assigned to a taxonomic node | `Taxonomic assignment success` | None |
| \|\| |  | Samtools Flagstat (pre-samtools filter) | Reads | Total reads in the bam file () | `flagstat_total` | read\_count |
| \|\| |  | Samtools Flagstat (pre-samtools filter) | Reads Mapped | Reads Mapped in the bam file () | `mapped_passed` | read\_count |
| \|\| |  | Samtools Flagstat (post-samtools filter) | Reads | Total reads in the bam file () | `flagstat_total` | read\_count |
| \|\| |  | endorSpy | Endogenous DNA (%) | Endogenous DNA (%) | `endogenous_dna` | None |
| \|\| |  | Samtools Flagstat (post-samtools filter) | Reads Mapped | Reads Mapped in the bam file () | `mapped_passed` | read\_count |
| \|\| |  | endorSpy | Endogenous DNA Post (%) | Endogenous DNA Post (%) | `endogenous_dna_post` | None |
| \|\| |  | Picard | % Dups | Mark Duplicates - Percent Duplication | `PERCENT_DUPLICATION` | None |
| \|\| |  | DamageProfiler | 5 Prime C>T 1st base | 5 Prime 1st base substitution frequency for C>T | `5 Prime1` | None |
| \|\| |  | DamageProfiler | 5 Prime C>T 2nd base | 5 Prime 2nd base substitution frequency for C>T | `5 Prime2` | None |
| \|\| |  | DamageProfiler | 3 Prime G>A 1st base | 3 Prime 1st base substitution frequency for G>A | `3 Prime1` | None |
| \|\| |  | DamageProfiler | 3 Prime G>A 2nd base | 3 Prime 2nd base substitution frequency for G>A | `3 Prime2` | None |
| \|\| |  | DamageProfiler | Mean read length | Mean read length | `mean_readlength` | read\_length |
| \|\| |  | DamageProfiler | Median read length | Median read length | `median` | read\_length |
| \|\| |  | mtnucratio | MT genome reads | Reads on the mitochondrial genome () | `mtreads` | read\_count |
| \|\| |  | mtnucratio | MT genome coverage | Average coverage (X) on mitochondrial genome. | `mt_cov_avg` | None |
| \|\| |  | mtnucratio | MT to Nuclear Ratio | Mitochondrial to nuclear reads ratio (MTNUC) | `mt_nuc_ratio` | None |
| \|\| |  | QualiMap | Aligned | Number of mapped reads () | `mapped_reads` | read\_count |
| \|\| |  | QualiMap | Mean cov | Mean coverage | `mean_coverage` | None |
| \|\| |  | QualiMap | Median cov | Median coverage | `median_coverage` | None |
| \|\| |  | QualiMap | ≥ 1X | Fraction of genome with at least 1X coverage | `1_x_pc` | None |
| \|\| |  | QualiMap | ≥ 2X | Fraction of genome with at least 2X coverage | `2_x_pc` | None |
| \|\| |  | QualiMap | ≥ 3X | Fraction of genome with at least 3X coverage | `3_x_pc` | None |
| \|\| |  | QualiMap | ≥ 4X | Fraction of genome with at least 4X coverage | `4_x_pc` | None |
| \|\| |  | QualiMap | ≥ 5X | Fraction of genome with at least 5X coverage | `5_x_pc` | None |
| \|\| |  | QualiMap | % GC | Mean GC content | `avg_gc` | None |
| \|\| |  | FastQC (pre-AdapterRemoval) | % Dups | % Duplicate Reads | `percent_duplicates` | None |
| \|\| |  | FastQC (pre-AdapterRemoval) | % Failed | Percentage of modules failed in FastQC report (includes those not plotted here) | `percent_fails` | None |
| \|\| |  | Adapter Removal | Reads Trimmed | Total trimmed reads () | `aligned_total` | read\_count |
| \|\| |  | FastQC (post-AdapterRemoval) | % Dups | % Duplicate Reads | `percent_duplicates` | None |
| \|\| |  | FastQC (post-AdapterRemoval) | % Failed | Percentage of modules failed in FastQC report (includes those not plotted here) | `percent_fails` | None |
| \|\| |  | DamageProfiler | Read length std. dev. | Read length std. dev. | `std` | read\_length |
| \|\| |  | mtnucratio | Genome coverage | Average coverage (X) on nuclear genome. | `nuc_cov_avg` | None |
| \|\| |  | mtnucratio | Genome reads | Reads on the nuclear genome () | `nucreads` | read\_count |
| \|\| |  | QualiMap | % Aligned | % mapped reads | `percentage_aligned` | None |
| \|\| |  | QualiMap | Total reads | Number of reads () | `total_reads` | read\_count |
| \|\| |  | QualiMap | Error rate | Alignment error rate. Total edit distance (SAM NM field) over the number of mapped bases | `general_error_rate` | None |
| \|\| |  | SexDetErrmine | Err Rate X | Rate of Error for Chr X | `RateErrX` | snp\_err\_rate |
| \|\| |  | SexDetErrmine | Err Rate Y | Rate of Error for Chr Y | `RateErrY` | snp\_err\_rate |
| \|\| |  | SexDetErrmine | Rate X | Number of positions on Chromosome X vs Autosomal positions. | `RateX` | snp\_count |
| \|\| |  | SexDetErrmine | Rate Y | Number of positions on Chromosome Y vs Autosomal positions. | `RateY` | snp\_count |
| \|\| |  | nuclear\_contamination | Number of SNPs | Number of SNPs | `Num_SNPs` | None |
| \|\| |  | nuclear\_contamination | Contamination Estimate (Method1\_MOM) | Contamination Estimate (Method1\_MOM) | `Method1_MOM_estimate` | None |
| \|\| |  | nuclear\_contamination | Estimate Error (Method1\_MOM) | Estimate Error (Method1\_MOM) | `Method1_MOM_SE` | None |
| \|\| |  | nuclear\_contamination | Contamination Estimate (Method1\_ML) | Contamination Estimate (Method1\_ML) | `Method1_ML_estimate` | None |
| \|\| |  | nuclear\_contamination | Estimate Error (Method1\_ML) | Estimate Error (Method1\_ML) | `Method1_ML_SE` | None |
| \|\| |  | nuclear\_contamination | Contamination Estimate (Method2\_MOM) | Contamination Estimate (Method2\_MOM) | `Method2_MOM_estimate` | None |
| \|\| |  | nuclear\_contamination | Estimate Error (Method2\_MOM) | Estimate Error (Method2\_MOM) | `Method2_MOM_SE` | None |
| \|\| |  | nuclear\_contamination | Contamination Estimate (Method2\_ML) | Contamination Estimate (Method2\_ML) | `Method2_ML_estimate` | None |
| \|\| |  | nuclear\_contamination | Estimate Error (Method2\_ML) | Estimate Error (Method2\_ML) | `Method2_ML_SE` | None |

Close

#### FastQC (pre-AdapterRemoval)

FastQC (pre-AdapterRemoval) is a quality control tool for high throughput sequence data, written by Simon Andrews at the Babraham Institute in Cambridge.

##### Sequence Counts Help

Sequence counts for each sample. Duplicate read counts are an estimate only.

This plot show the total number of reads, broken down into unique and duplicate
if possible (only more recent versions of FastQC give duplicate info).

You can read more about duplicate calculation in the
FastQC documentation.
A small part has been copied here for convenience:

*Only sequences which first appear in the first 100,000 sequences
in each file are analysed. This should be enough to get a good impression
for the duplication levels in the whole file. Each sequence is tracked to
the end of the file to give a representative count of the overall duplication level.*

*The duplication detection requires an exact sequence match over the whole length of
the sequence. Any reads over 75bp in length are truncated to 50bp for this analysis.*

Number of reads
Percentages

loading..

---

##### Sequence Quality Histograms Help

The mean quality value across each base position in the read.

To enable multiple samples to be plotted on the same graph, only the mean quality
scores are plotted (unlike the box plots seen in FastQC reports).

Taken from the FastQC help:

*The y-axis on the graph shows the quality scores. The higher the score, the better
the base call. The background of the graph divides the y axis into very good quality
calls (green), calls of reasonable quality (orange), and calls of poor quality (red).
The quality of calls on most platforms will degrade as the run progresses, so it is
common to see base calls falling into the orange area towards the end of a read.*

loading..

---

##### Per Sequence Quality Scores Help

The number of reads with average quality scores. Shows if a subset of reads has poor quality.

From the FastQC help:

*The per sequence quality score report allows you to see if a subset of your
sequences have universally low quality values. It is often the case that a
subset of sequences will have universally poor quality, however these should
represent only a small percentage of the total sequences.*

loading..

---

##### Per Base Sequence Content Help

The proportion of each base position for which each of the four normal DNA bases has been called.

To enable multiple samples to be shown in a single plot, the base composition data
is shown as a heatmap. The colours represent the balance between the four bases:
an even distribution should give an even muddy brown colour. Hover over the plot
to see the percentage of the four bases under the cursor.

**To see the data as a line plot, as in the original FastQC graph, click on a sample track.**

From the FastQC help:

*Per Base Sequence Content plots out the proportion of each base position in a
file for which each of the four normal DNA bases has been called.*

*In a random library you would expect that there would be little to no difference
between the different bases of a sequence run, so the lines in this plot should
run parallel with each other. The relative amount of each base should reflect
the overall amount of these bases in your genome, but in any case they should
not be hugely imbalanced from each other.*

*It's worth noting that some types of library will always produce biased sequence
composition, normally at the start of the read. Libraries produced by priming
using random hexamers (including nearly all RNA-Seq libraries) and those which
were fragmented using transposases inherit an intrinsic bias in the positions
at which reads start. This bias does not concern an absolute sequence, but instead
provides enrichement of a number of different K-mers at the 5' end of the reads.
Whilst this is a true technical bias, it isn't something which can be corrected
by trimming and in most cases doesn't seem to adversely affect the downstream
analysis.*

Click a sample row to see a line plot for that dataset.

###### Rollover for sample name

 Export Plot

Position: -

%T: -

%C: -

%A: -

%G: -

---

##### Per Sequence GC Content Help

The average GC content of reads. Normal random library typically have a
roughly normal distribution of GC content.

From the FastQC help:

*This module measures the GC content across the whole length of each sequence
in a file and compares it to a modelled normal distribution of GC content.*

*In a normal random library you would expect to see a roughly normal distribution
of GC content where the central peak corresponds to the overall GC content of
the underlying genome. Since we don't know the the GC content of the genome the
modal GC content is calculated from the observed data and used to build a
reference distribution.*

*An unusually shaped distribution could indicate a contaminated library or
some other kinds of biased subset. A normal distribution which is shifted
indicates some systematic bias which is independent of base position. If there
is a systematic bias which creates a shifted normal distribution then this won't
be flagged as an error by the module since it doesn't know what your genome's
GC content should be.*

Percentages
Counts

loading..

---

##### Per Base N Content Help

The percentage of base calls at each position for which an `N` was called.

From the FastQC help:

*If a sequencer is unable to make a base call with sufficient confidence then it will
normally substitute an `N` rather than a conventional base call. This graph shows the
percentage of base calls at each position for which an `N` was called.*

*It's not unusual to see a very low proportion of Ns appearing in a sequence, especially
nearer the end of a sequence. However, if this proportion rises above a few percent
it suggests that the analysis pipeline was unable to interpret the data well enough to
make valid base calls.*

loading..

---

##### Sequence Length Distribution

The distribution of fragment sizes (read lengths) found.
See the FastQC help

loading..

---

##### Sequence Duplication Levels Help

The relative level of duplication found for every sequence.

From the FastQC Help:

*In a diverse library most sequences will occur only once in the final set.
A low level of duplication may indicate a very high level of coverage of the
target sequence, but a high level of duplication is more likely to indicate
some kind of enrichment bias (eg PCR over amplification). This graph shows
the degree of duplication for every sequence in a library: the relative
number of sequences with different degrees of duplication.*

*Only sequences which first appear in the first 100,000 sequences
in each file are analysed. This should be enough to get a good impression
for the duplication levels in the whole file. Each sequence is tracked to
the end of the file to give a representative count of the overall duplication level.*

*The duplication detection requires an exact sequence match over the whole length of
the sequence. Any reads over 75bp in length are truncated to 50bp for this analysis.*

*In a properly diverse library most sequences should fall into the far left of the
plot in both the red and blue lines. A general level of enrichment, indicating broad
oversequencing in the library will tend to flatten the lines, lowering the low end
and generally raising other categories. More specific enrichments of subsets, or
the presence of low complexity contaminants will tend to produce spikes towards the
right of the plot.*

loading..

---

##### Overrepresented sequences Help

The total amount of overrepresented sequences found in each library.

FastQC calculates and lists overrepresented sequences in FastQ files. It would not be
possible to show this for all samples in a MultiQC report, so instead this plot shows
the *number of sequences* categorized as over represented.

Sometimes, a single sequence may account for a large number of reads in a dataset.
To show this, the bars are split into two: the first shows the overrepresented reads
that come from the single most common sequence. The second shows the total count
from all remaining overrepresented sequences.

From the FastQC Help:

*A normal high-throughput library will contain a diverse set of sequences, with no
individual sequence making up a tiny fraction of the whole. Finding that a single
sequence is very overrepresented in the set either means that it is highly biologically
significant, or indicates that the library is contaminated, or not as diverse as you expected.*

*FastQC lists all of the sequences which make up more than 0.1% of the total.
To conserve memory only sequences which appear in the first 100,000 sequences are tracked
to the end of the file. It is therefore possible that a sequence which is overrepresented
but doesn't appear at the start of the file for some reason could be missed by this module.*

18 samples had less than 1% of reads made up of overrepresented sequences

---

##### Adapter Content Help

The cumulative percentage count of the proportion of your
library which has seen each of the adapter sequences at each position.

Note that only samples with ≥ 0.1% adapter contamination are shown.

There may be several lines per sample, as one is shown for each adapter
detected in the file.

From the FastQC Help:

*The plot shows a cumulative percentage count of the proportion
of your library which has seen each of the adapter sequences at each position.
Once a sequence has been seen in a read it is counted as being present
right through to the end of the read so the percentages you see will only
increase as the read length goes on.*

loading..

---

##### Status Checks Help

Status for each FastQC section showing whether results seem entirely normal (green),
slightly abnormal (orange) or very unusual (red).

FastQC assigns a status for each section of the report.
These give a quick evaluation of whether the results of the analysis seem
entirely normal (green), slightly abnormal (orange) or very unusual (red).

It is important to stress that although the analysis results appear to give a pass/fail result,
these evaluations must be taken in the context of what you expect from your library.
A 'normal' sample as far as FastQC is concerned is random and diverse.
Some experiments may be expected to produce libraries which are biased in particular ways.
You should treat the summary evaluations therefore as pointers to where you should concentrate
your attention and understand why your library may not look random and diverse.

Specific guidance on how to interpret the output of each module can be found in the relevant
report section, or in the FastQC help.

In this heatmap, we summarise all of these into a single heatmap for a quick overview.
Note that not all FastQC sections have plots in MultiQC reports, but all status checks
are shown in this heatmap.

Sort by highlight

loading..

---

#### Adapter Removal

Adapter Removal rapid adapter trimming, identification, and read merging

##### Retained and Discarded Paired-End Collapsed

The number of retained and discarded reads.

Number of Reads
Percentages

loading..

---

##### Length Distribution Paired End Collapsed

The length distribution of reads after processing adapter alignment.

All
Mate1
Mate2
Singleton
Collapsed
Collapsed Truncated
Discarded

loading..

---

#### FastQC (post-AdapterRemoval)

FastQC (post-AdapterRemoval) is a quality control tool for high throughput sequence data, written by Simon Andrews at the Babraham Institute in Cambridge.

##### Sequence Counts Help

Sequence counts for each sample. Duplicate read counts are an estimate only.

This plot show the total number of reads, broken down into unique and duplicate
if possible (only more recent versions of FastQC give duplicate info).

You can read more about duplicate calculation in the
FastQC documentation.
A small part has been copied here for convenience:

*Only sequences which first appear in the first 100,000 sequences
in each file are analysed. This should be enough to get a good impression
for the duplication levels in the whole file. Each sequence is tracked to
the end of the file to give a representative count of the overall duplication level.*

*The duplication detection requires an exact sequence match over the whole length of
the sequence. Any reads over 75bp in length are truncated to 50bp for this analysis.*

Number of reads
Percentages

loading..

---

##### Sequence Quality Histograms Help

The mean quality value across each base position in the read.

To enable multiple samples to be plotted on the same graph, only the mean quality
scores are plotted (unlike the box plots seen in FastQC reports).

Taken from the FastQC help:

*The y-axis on the graph shows the quality scores. The higher the score, the better
the base call. The background of the graph divides the y axis into very good quality
calls (green), calls of reasonable quality (orange), and calls of poor quality (red).
The quality of calls on most platforms will degrade as the run progresses, so it is
common to see base calls falling into the orange area towards the end of a read.*

loading..

---

##### Per Sequence Quality Scores Help

The number of reads with average quality scores. Shows if a subset of reads has poor quality.

From the FastQC help:

*The per sequence quality score report allows you to see if a subset of your
sequences have universally low quality values. It is often the case that a
subset of sequences will have universally poor quality, however these should
represent only a small percentage of the total sequences.*

loading..

---

##### Per Base Sequence Content Help

The proportion of each base position for which each of the four normal DNA bases has been called.

To enable multiple samples to be shown in a single plot, the base composition data
is shown as a heatmap. The colours represent the balance between the four bases:
an even distribution should give an even muddy brown colour. Hover over the plot
to see the percentage of the four bases under the cursor.

**To see the data as a line plot, as in the original FastQC graph, click on a sample track.**

From the FastQC help:

*Per Base Sequence Content plots out the proportion of each base position in a
file for which each of the four normal DNA bases has been called.*

*In a random library you would expect that there would be little to no difference
between the different bases of a sequence run, so the lines in this plot should
run parallel with each other. The relative amount of each base should reflect
the overall amount of these bases in your genome, but in any case they should
not be hugely imbalanced from each other.*

*It's worth noting that some types of library will always produce biased sequence
composition, normally at the start of the read. Libraries produced by priming
using random hexamers (including nearly all RNA-Seq libraries) and those which
were fragmented using transposases inherit an intrinsic bias in the positions
at which reads start. This bias does not concern an absolute sequence, but instead
provides enrichement of a number of different K-mers at the 5' end of the reads.
Whilst this is a true technical bias, it isn't something which can be corrected
by trimming and in most cases doesn't seem to adversely affect the downstream
analysis.*

Click a sample row to see a line plot for that dataset.

###### Rollover for sample name

 Export Plot

Position: -

%T: -

%C: -

%A: -

%G: -

---

##### Per Sequence GC Content Help

The average GC content of reads. Normal random library typically have a
roughly normal distribution of GC content.

From the FastQC help:

*This module measures the GC content across the whole length of each sequence
in a file and compares it to a modelled normal distribution of GC content.*

*In a normal random library you would expect to see a roughly normal distribution
of GC content where the central peak corresponds to the overall GC content of
the underlying genome. Since we don't know the the GC content of the genome the
modal GC content is calculated from the observed data and used to build a
reference distribution.*

*An unusually shaped distribution could indicate a contaminated library or
some other kinds of biased subset. A normal distribution which is shifted
indicates some systematic bias which is independent of base position. If there
is a systematic bias which creates a shifted normal distribution then this won't
be flagged as an error by the module since it doesn't know what your genome's
GC content should be.*

Percentages
Counts

loading..

---

##### Per Base N Content Help

The percentage of base calls at each position for which an `N` was called.

From the FastQC help:

*If a sequencer is unable to make a base call with sufficient confidence then it will
normally substitute an `N` rather than a conventional base call. This graph shows the
percentage of base calls at each position for which an `N` was called.*

*It's not unusual to see a very low proportion of Ns appearing in a sequence, especially
nearer the end of a sequence. However, if this proportion rises above a few percent
it suggests that the analysis pipeline was unable to interpret the data well enough to
make valid base calls.*

loading..

---

##### Sequence Length Distribution

The distribution of fragment sizes (read lengths) found.
See the FastQC help

loading..

---

##### Sequence Duplication Levels Help

The relative level of duplication found for every sequence.

From the FastQC Help:

*In a diverse library most sequences will occur only once in the final set.
A low level of duplication may indicate a very high level of coverage of the
target sequence, but a high level of duplication is more likely to indicate
some kind of enrichment bias (eg PCR over amplification). This graph shows
the degree of duplication for every sequence in a library: the relative
number of sequences with different degrees of duplication.*

*Only sequences which first appear in the first 100,000 sequences
in each file are analysed. This should be enough to get a good impression
for the duplication levels in the whole file. Each sequence is tracked to
the end of the file to give a representative count of the overall duplication level.*

*The duplication detection requires an exact sequence match over the whole length of
the sequence. Any reads over 75bp in length are truncated to 50bp for this analysis.*

*In a properly diverse library most sequences should fall into the far left of the
plot in both the red and blue lines. A general level of enrichment, indicating broad
oversequencing in the library will tend to flatten the lines, lowering the low end
and generally raising other categories. More specific enrichments of subsets, or
the presence of low complexity contaminants will tend to produce spikes towards the
right of the plot.*

loading..

---

##### Overrepresented sequences Help

The total amount of overrepresented sequences found in each library.

FastQC calculates and lists overrepresented sequences in FastQ files. It would not be
possible to show this for all samples in a MultiQC report, so instead this plot shows
the *number of sequences* categorized as over represented.

Sometimes, a single sequence may account for a large number of reads in a dataset.
To show this, the bars are split into two: the first shows the overrepresented reads
that come from the single most common sequence. The second shows the total count
from all remaining overrepresented sequences.

From the FastQC Help:

*A normal high-throughput library will contain a diverse set of sequences, with no
individual sequence making up a tiny fraction of the whole. Finding that a single
sequence is very overrepresented in the set either means that it is highly biologically
significant, or indicates that the library is contaminated, or not as diverse as you expected.*

*FastQC lists all of the sequences which make up more than 0.1% of the total.
To conserve memory only sequences which appear in the first 100,000 sequences are tracked
to the end of the file. It is therefore possible that a sequence which is overrepresented
but doesn't appear at the start of the file for some reason could be missed by this module.*

12 samples had less than 1% of reads made up of overrepresented sequences

---

##### Adapter Content Help

The cumulative percentage count of the proportion of your
library which has seen each of the adapter sequences at each position.

Note that only samples with ≥ 0.1% adapter contamination are shown.

There may be several lines per sample, as one is shown for each adapter
detected in the file.

From the FastQC Help:

*The plot shows a cumulative percentage count of the proportion
of your library which has seen each of the adapter sequences at each position.
Once a sequence has been seen in a read it is counted as being present
right through to the end of the read so the percentages you see will only
increase as the read length goes on.*

loading..

---

##### Status Checks Help

Status for each FastQC section showing whether results seem entirely normal (green),
slightly abnormal (orange) or very unusual (red).

FastQC assigns a status for each section of the report.
These give a quick evaluation of whether the results of the analysis seem
entirely normal (green), slightly abnormal (orange) or very unusual (red).

It is important to stress that although the analysis results appear to give a pass/fail result,
these evaluations must be taken in the context of what you expect from your library.
A 'normal' sample as far as FastQC is concerned is random and diverse.
Some experiments may be expected to produce libraries which are biased in particular ways.
You should treat the summary evaluations therefore as pointers to where you should concentrate
your attention and understand why your library may not look random and diverse.

Specific guidance on how to interpret the output of each module can be found in the relevant
report section, or in the FastQC help.

In this heatmap, we summarise all of these into a single heatmap for a quick overview.
Note that not all FastQC sections have plots in MultiQC reports, but all status checks
are shown in this heatmap.

Sort by highlight

loading..

---

#### MALT

MALT performs alignment of metagenomic reads against a database of reference sequences (such as NR, GenBank or Silva) and produces a MEGAN RMA file as output.

##### Metagenomic Mappability

Number of mapped reads.

Counts
Percentages

loading..

---

##### Taxonomic assignment success

Shows the number of mapped reads assigned to a taxonomic node.

Counts
Percentages

loading..

---

#### Samtools Flagstat (pre-samtools filter)

Samtools is a suite of programs for interacting with high-throughput sequencing data.

##### Samtools Flagstat

This module parses the output from `samtools flagstat`. All numbers in millions.

loading..

---

#### Samtools Flagstat (post-samtools filter)

Samtools is a suite of programs for interacting with high-throughput sequencing data.

##### Samtools Flagstat

This module parses the output from `samtools flagstat`. All numbers in millions.

loading..

---

#### Picard

Picard is a set of Java command line tools for manipulating high-throughput sequencing data.

##### Mark Duplicates Help

Number of reads, categorised by duplication state. **Pair counts are doubled** - see help text for details.

The table in the Picard metrics file contains some columns referring
read pairs and some referring to single reads.

To make the numbers in this plot sum correctly, values referring to pairs are doubled
according to the scheme below:

- `READS_IN_DUPLICATE_PAIRS = 2 * READ_PAIR_DUPLICATES`
- `READS_IN_UNIQUE_PAIRS = 2 * (READ_PAIRS_EXAMINED - READ_PAIR_DUPLICATES)`
- `READS_IN_UNIQUE_UNPAIRED = UNPAIRED_READS_EXAMINED - UNPAIRED_READ_DUPLICATES`
- `READS_IN_DUPLICATE_PAIRS_OPTICAL = 2 * READ_PAIR_OPTICAL_DUPLICATES`
- `READS_IN_DUPLICATE_PAIRS_NONOPTICAL = READS_IN_DUPLICATE_PAIRS - READS_IN_DUPLICATE_PAIRS_OPTICAL`
- `READS_IN_DUPLICATE_UNPAIRED = UNPAIRED_READ_DUPLICATES`
- `READS_UNMAPPED = UNMAPPED_READS`

Number of Reads
Percentages

loading..

---

#### Preseq

Preseq estimates the complexity of a library, showing how many additional
unique reads are sequenced for increasing total read count.
A shallow curve indicates complexity saturation. The dashed line
shows a perfectly complex library where total reads = unique reads.

##### Complexity curve

Note that the x axis is trimmed at the point where all the datasets show 80% of their maximum y-value, to avoid ridiculous scales.

loading..

---

#### DamageProfiler

DamageProfiler a tool to determine damage patterns on ancient DNA.

##### 3P misincorporation plot Help

3' misincorporation plot for G>A substitutions

This plot shows the frequency of G>A substitutions at the 3' read ends. Typically, one would observe high substitution percentages for ancient DNA, whereas modern DNA does not show these in higher extents.

loading..

---

##### 5P misincorporation plot Help

5' misincorporation plot for C>T substitutions

This plot shows the frequency of C>T substitutions at the 5' read ends. Typically, one would observe high substitution percentages for ancient DNA, whereas modern DNA does not show these in higher extents.

loading..

---

##### Forward read length distribution Help

Read length distribution for forward strand (+) reads.

This plot shows the read length distribution of the forward reads in the investigated sample. Reads below lengths of 30bp are typically filtered, so the plot doesn't show these in many cases. A shifted distribution of read lengths towards smaller read lengths (e.g around 30-50bp) is also an indicator of ancient DNA.

loading..

---

##### Reverse read length distribution Help

Read length distribution for reverse strand (-) reads.

This plot shows the read length distribution of the reverse reads in the investigated sample. Reads below lengths of 30bp are typically filtered, so the plot doesn't show these in many cases. A shifted distribution of read lengths towards smaller read lengths (e.g around 30-50bp) is also an indicator of ancient DNA.

loading..

---

#### QualiMap

QualiMap is a platform-independent application to facilitate the quality control of alignment sequencing data and its derivatives like feature counts.

##### Coverage histogram Help

Distribution of the number of locations in the reference genome with a given depth of coverage.

For a set of DNA or RNA reads mapped to a reference sequence, such as a genome
or transcriptome, the depth of coverage at a given base position is the number
of high-quality reads that map to the reference at that position
(Sims et al. 2014).

Bases of a reference sequence (y-axis) are groupped by their depth of coverage
(*0×, 1×, …, N×*) (x-axis). This plot shows
the frequency of coverage depths relative to the reference sequence for each
read dataset, which provides an indirect measure of the level and variation of
coverage depth in the corresponding sequenced sample.

If reads are randomly distributed across the reference sequence, this plot
should resemble a Poisson distribution (Lander & Waterman 1988), with a peak indicating approximate
depth of coverage, and more uniform coverage depth being reflected in a narrower
spread. The optimal level of coverage depth depends on the aims of the
experiment, though it should at minimum be sufficiently high to adequately
address the biological question; greater uniformity of coverage is generally
desirable, because it increases breadth of coverage for a given depth of
coverage, allowing equivalent results to be achieved at a lower sequencing depth
(Sampson
et al. 2011; Sims
et al. 2014). However, it is difficult to achieve uniform coverage
depth in practice, due to biases introduced during sample preparation
(van
Dijk et al. 2014), sequencing (Ross et al. 2013) and read mapping
(Sims et al. 2014).

This plot may include a small peak for regions of the reference sequence with
zero depth of coverage. Such regions may be absent from the given sample (due
to a deletion or structural rearrangement), present in the sample but not
successfully sequenced (due to bias in sequencing or preparation), or sequenced
but not successfully mapped to the reference (due to the choice of mapping
algorithm, the presence of repeat sequences, or mismatches caused by variants
or sequencing errors). Related factors cause most datasets to contain some
unmapped reads (Sims
et al. 2014).

loading..

---

##### Cumulative genome coverage Help

Percentage of the reference genome with at least the given depth of coverage.

For a set of DNA or RNA reads mapped to a reference sequence, such as a genome
or transcriptome, the depth of coverage at a given base position is the number
of high-quality reads that map to the reference at that position, while the
breadth of coverage is the fraction of the reference sequence to which reads
have been mapped with at least a given depth of coverage
(Sims et al. 2014).

Defining coverage breadth in terms of coverage depth is useful, because
sequencing experiments typically require a specific minimum depth of coverage
over the region of interest (Sims et al. 2014), so the extent of the reference sequence
that is amenable to analysis is constrained to lie within regions that have
sufficient depth. With inadequate sequencing breadth, it can be difficult to
distinguish the absence of a biological feature (such as a gene) from a lack
of data (Green 2007).

For increasing coverage depths (*1×, 2×, …, N×*),
coverage breadth is calculated as the percentage of the reference
sequence that is covered by at least that number of reads, then plots
coverage breadth (y-axis) against coverage depth (x-axis). This plot
shows the relationship between sequencing depth and breadth for each read
dataset, which can be used to gauge, for example, the likely effect of a
minimum depth filter on the fraction of a genome available for analysis.

loading..

---

##### GC content distribution Help

Each solid line represents the distribution of GC content of mapped reads for a given sample.

GC bias is the difference between the guanine-cytosine content
(GC-content) of a set of sequencing reads and the GC-content of the DNA
or RNA in the original sample. It is a well-known issue with sequencing
systems, and may be introduced by PCR amplification, among other factors
(Benjamini
& Speed 2012; Ross et al. 2013).

QualiMap calculates the GC-content of individual mapped reads, then
groups those reads by their GC-content (*1%, 2%, …, 100%*), and
plots the frequency of mapped reads (y-axis) at each level of GC-content
(x-axis). This plot shows the GC-content distribution of mapped reads
for each read dataset, which should ideally resemble that of the
original sample. It can be useful to display the GC-content distribution
of an appropriate reference sequence for comparison, and QualiMap has an
option to do this (see the Qualimap 2 documentation).

loading..

---

#### SexDetErrmine

SexDetErrmine A python script to calculate the relative coverage of X and Y chromosomes,
and their associated error bars, from the depth of coverage at specified SNPs.

##### Relative Coverage Help

The coverage on the X vs Y chromosome, relative to coverage on the Autosomes.

Males are expected to have a roughly equal X- and Y-rates, while females are expected to have a Y-rate of 0 and an X-rate of 1.
Placement between the two clusters can be indicative of contamination, while placement with higher than expected X- and/or Y-rates can be indicative of sex chromosome aneuploidy.

loading..

---

##### Read Counts

The number of reads covering positions on the autosomes, X and Y chromosomes.

Counts
Percentages

loading..

---

#### nf-core/eager Software Versions

are collected at run time from the software output.

nf-core/eager
:   `v2.2.0dev`

Nextflow
:   `v20.04.1`

FastQC
:   `v0.11.9`

MultiQC
:   `v1.9`

AdapterRemoval
:   `v2.3.0`

fastP
:   `v0.20.1`

BWA
:   `v0.7.17-r1188`

Bowtie2
:   `v2.4.1`

circulargenerator
:   `v1.0`

Samtools
:   `v1.9`

endorS.py
:   `v0.4`

DeDup
:   `v0.12.6`

Picard MarkDuplicates
:   `v2.22.9`

Qualimap
:   `v2.2.2-dev`

Preseq
:   `v2.0.3`

GATK HaplotypeCaller
:   `v4.1.7.0`

freebayes
:   `v1.3.2-dirty`

sequenceTools
:   `v1.4.0.5`

VCF2genome
:   `v0.91`

MTNucRatioCalculator
:   `v0.7`

bedtools
:   `v2.29.2`

DamageProfiler
:   `v0.4.9`

bamUtil
:   `v1.0.14`

pmdtools
:   `v0.50`

angsd
:   `v0.933`

sexdeterrmine
:   `v1.1.1`

multivcfanalyzer
:   `v0.85.2`

malt
:   `v0.4.1`

kraken
:   `v2.0.9-beta`

maltextract
:   `v1.7`

---

#### nf-core/eager Workflow Summary

- this information is collected when the pipeline is started.

Pipeline Name
:   `nf-core/eager`

Pipeline Version
:   `2.2.0dev`

Pipeline Release
:   `dev`

Run Name
:   `boring_gates`

Input
:   `barquera2020_pathogenscreening.tsv`

Convert input BAM?
:   `No`

Fasta Ref
:   `ftp://ftp-trace.ncbi.nih.gov/1000genomes/ftp/technical/reference/phase2_reference_assembly_sequence/hs37d5.fa.gz`

BAM Index Type
:   `CSI`

Skipping FASTQC?
:   `No`

Skipping AdapterRemoval?
:   `No`

Skip Read Merging
:   `fastq`

Skip Adapter Trimming
:   `No`

Running BAM filtering
:   `Yes`

Run Fastq Host Removal
:   `No`

Skipping Preseq?
:   `No`

Skipping Deduplication?
:   `No`

Skipping DamageProfiler?
:   `No`

Skipping Qualimap?
:   `No`

Run BAM Trimming?
:   `No`

Run PMDtools?
:   `No`

Run Genotyping?
:   `No`

Run MultiVCFAnalyzer
:   `No`

Run VCF2Genome
:   `No`

Run SexDetErrMine
:   `Yes`

Run Nuclear Contamination Estimation
:   `Yes`

Run Bedtools Coverage
:   `No`

Run Metagenomic Binning
:   `Yes`

Metagenomic Tool
:   `malt`

Run MaltExtract
:   `Yes`

Max Resources
:   `2 TB memory, 128 cpus, 24d 20h 31m 24s time per job`

Output Dir
:   `./results`

Working Dir
:   `/projects1/users/fellows/nextflow/eager2/publication/benchmarking_pathogen/work`

Container Engine
:   `singularity`

Container
:   `nfcore/eager:dev`

Current Home
:   `/projects1/clusterhomes/fellows`

Current User
:   `fellows`

Script Dir
:   `/projects1/clusterhomes/fellows/.nextflow/assets/nf-core/eager`

Config Profile
:   `microbiome_screening,sdag,shh`

User
:   `fellows`

Config Description
:   `nf-core/eager SHH profile provided by nf-core/configs`

Config Contact
:   `James Fellows Yates (@jfy133)`

Config URL
:   `https://shh.mpg.de`

**MultiQC v1.9** 
- Written by Phil Ewels,
available on GitHub.

This report uses HighCharts,
jQuery,
jQuery UI,
Bootstrap,
FileSaver.js and
clipboard.js.

×

##### Plot Table Data

Select Column

Select Column

Please select two table columns.

Close

×

##### Regex Help

Toolbox search strings can behave as regular expressions (regexes). Click a button below to see an example of it in action. Try modifying them yourself in the text box.

`^` (start of string)
`$` (end of string)
`[]` (character choice)
`\d` (shorthand for `[0-9]`)
`\w` (shorthand for `[0-9a-zA-Z_]`)
`.` (any character)
`\.` (literal full stop)
`()` `|` (group / separator)
`*` (prev char 0 or more)
`+` (prev char 1 or more)
`?` (prev char 0 or 1)
`{}` (char num times)
`{,}` (count range)

```
samp_1
samp_1_edited
samp_2
samp_2_edited
samp_3
samp_3_edited
prepended_samp_1
tmp_samp_1_edited
tmpp_samp_1_edited
tmppp_samp_1_edited
#samp_1_edited.tmp
samp_11
samp_11111
```

See regex101.com for a more heavy duty testing suite.

Close
