## Supplemental file archive of walkthrough and results files for "Reproducible, portable, and efficient ancient genome reconstruction with nf-core/eager": barquera2020_sdag_multiqc_1_10_report.html

Format:

Tab-separated
Comma-separated
JSON

Note that additional data was saved in `multiqc1_10_data` when this report was generated.

---

###### Choose Plots

 All
 None


---


   Download Plot Images

If you use plots from MultiQC in a publication or presentation, please cite:

> **MultiQC: Summarize analysis results for multiple tools and samples in a single report**  
> *Philip Ewels, Måns Magnusson, Sverker Lundin and Max Käller*  
> Bioinformatics (2016)  
> doi: 10.1093/bioinformatics/btw354  
> PMID: 27312411

##### Save Settings

You can save the toolbox settings for this report to the browser.

 Save


---

##### Load Settings

Choose a saved report profile from the dropdown box below:

[ select ]

Load
 Delete
 Set default
 Clear default

##### About MultiQC

This report was generated using MultiQC, version 1.10.dev0

You can see a YouTube video describing how to use MultiQC reports here:
https://youtu.be/qPbIlO\_KWN0

For more information about MultiQC, including other videos and
extensive documentation, please visit http://multiqc.info

You can report bugs, suggest improvements and find the source code for MultiQC on GitHub:
https://github.com/ewels/MultiQC

MultiQC is published in Bioinformatics:

> **MultiQC: Summarize analysis results for multiple tools and samples in a single report**  
> *Philip Ewels, Måns Magnusson, Sverker Lundin and Max Käller*  
> Bioinformatics (2016)  
> doi: 10.1093/bioinformatics/btw354  
> PMID: 27312411

A modular tool to aggregate results from bioinformatics analyses across many samples into a single report.

> This report has been generated by the nf-core/eager analysis pipeline. For information about how to interpret these results, please see the documentation.

##### JavaScript Disabled

MultiQC reports use JavaScript for plots and toolbox functions. It looks like
you have JavaScript disabled in your web browser. Please note that many of the report
functions will not work as intended.

Loading report..

Report
generated on 2020-10-06, 13:04
based on data in:
`/projects1/users/fellows/nextflow/eager2/publication/benchmarking_pathogen/results`

---

×
don't show again

**Welcome!** Not sure where to start?  
Watch a tutorial video
  *(6:06)*

Number of mapped reads.

Counts
Percentages

loading..

---

#### Taxonomic assignment success

Shows the number of mapped reads assigned to a taxonomic node.

Counts
Percentages

loading..

---

### HOPS

HOPS is an ancient DNA characteristics screening tool of output from the metagenomic aligner MALT.

#### Potential Candidates Help

Heatmap of candidate taxa for downstream aDNA analysis, with
intensity representing additive categories of possible 'positive'
hits.

HOPS assigns a category based on how many ancient DNA
characteristics a given node (i.e. taxon) in a sample has.
The colours indicate the following:

- **Grey** - No characteristics detected
- **Yellow** - Small edit distance from reference
- **Orange** - Typical aDNA damage pattern
- **Red** - Small edit distance *and* aDNA damage pattern

A red category typically indicates a good candidate for further investigation
in downstream analysis.

loading..

---

#### Read Counts

The number of reads covering positions on the autosomes, X and Y chromosomes.

Counts
Percentages

loading..

**MultiQC v1.10.dev0** 
- Written by Phil Ewels,
available on GitHub.

This report uses HighCharts,
jQuery,
jQuery UI,
Bootstrap,
FileSaver.js and
clipboard.js.

×

#### Plot Table Data

Select Column

Select Column

Please select two table columns.

Close

×

#### Regex Help

Toolbox search strings can behave as regular expressions (regexes). Click a button below to see an example of it in action. Try modifying them yourself in the text box.

`^` (start of string)
`$` (end of string)
`[]` (character choice)
`\d` (shorthand for `[0-9]`)
`\w` (shorthand for `[0-9a-zA-Z_]`)
`.` (any character)
`\.` (literal full stop)
`()` `|` (group / separator)
`*` (prev char 0 or more)
`+` (prev char 1 or more)
`?` (prev char 0 or 1)
`{}` (char num times)
`{,}` (count range)

```
samp_1
samp_1_edited
samp_2
samp_2_edited
samp_3
samp_3_edited
prepended_samp_1
tmp_samp_1_edited
tmpp_samp_1_edited
tmppp_samp_1_edited
#samp_1_edited.tmp
samp_11
samp_11111
```

See regex101.com for a more heavy duty testing suite.

Close
