## Supplemental file archive of walkthrough and results files for "Reproducible, portable, and efficient ancient genome reconstruction with nf-core/eager": demonstration_setup.md

# nf-core/eager paper - demonstration

This document describes the set up of the pipeline runs used to demonstrate the
additional functionality of nf-core/eager, primarily running metagenomic
screening for pathogens alongside running human DNA screening. This is run on an
institutional server at the Max Planck Institute for the Science of Human
History (MPI-SHH) due to the requirements of large servers for screening via
MALT (e.g. a 2 TB memory node for NCBI Nt database)

Here we will attempt to re-analyse the 'manual' human screening statistics and
pathogen screening  in Barquera et al. (2020, Current Biology) in a single
nf-core/eager run.

## Environment

The environment this analysis was run on was on a cluster of multiple nodes
running Ubuntu 18.04.4 LTS (kernal v4.15.0-24-generic) with Intel(R) Xeon(R) CPU
E5 series CPUs. The cluster has a SLURM scheduling system, with the
nf-core/eager config files avaliable in the nf-core configs central repository:
https://github.com/nf-core/configs/blob/master/conf/shh.config (for standard
profiles) and
https://github.com/nf-core/configs/blob/master/conf/pipeline/eager/shh.config
(for pipeline specific parameters).

This analysis further uses singularity version XXX, Nextflow version 20.04.1
(build 5335), and nf-core/eager version 2.2.0dev commit e7471a78a3.

## Set up

Barquera et al. originally generated shotgun data from individuals excavated
from a 16th century burial ground in Mexico City (Mexico) for human population
genetics analysis. In addition, they screened the off-target reads from the
teeth samples for a list of pathogenic taxa. The results from the various
analyses applied to shotgun screening was then used to guide in-solution
enrichment experiments.

Two teeth (sample) from each individual (e.g. SJN001) were analysed, and for
each sample (e.g. SJN001.A and .B) two libraries were constructed (e.g.
SJN001.A0101 and SJN001.A0102), with the former constructed without UDG
treatment and the latter with the 'UDG-half' variant to have one library with
full damage and one library with damage on the first base respectively.

A summary of the results at and Individual level can be seen here:

| Lab Individual ID | Raw Reads (millions) | Mean Damage 1st Base 5' (UDG half) | Mean Median Fragment Length (UDG Half) | Biological Sex | Pathogen           |
|-------------------|----------------------|------------------------------------|----------------------------------------|----------------|--------------------|
| SJN001            | ~10                  | 0.065                              | 50.5                                   | Male           | Hepatitis B        |
| SJN002            | ~10                  | 0.08                               | 49.5                                   | Male           |                    |
| SJN003            | ~10                  | 0.065                              | 51                                     | Male           | Treponema Pallidum |

These samples and libraries represent different sequencing strategies, and
therefore the following TSV file as input to the nf-core/eager run was set up,
with the primary difference between a mixture of paired-end and single-end
sequencing chemistries (SeqType)

| Sample_Name | Library_ID   | Lane | Colour_Chemistry | SeqType | Organism     | Strandedness | UDG_Treatment | R1                                                                             | R2                                                                             | BAM |
|-------------|--------------|------|------------------|---------|--------------|--------------|---------------|--------------------------------------------------------------------------------|--------------------------------------------------------------------------------|-----|
| SJN001      | SJN001.A0101 | 3    | 4                | SE      | Homo sapiens | double       | none          | ftp://ftp.sra.ebi.ac.uk/vol1/fastq/ERR406/008/ERR4065478/ERR4065478.fastq.gz   | NA                                                                             | NA  |
| SJN001      | SJN001.B0101 | 3    | 4                | SE      | Homo sapiens | double       | none          | ftp://ftp.sra.ebi.ac.uk/vol1/fastq/ERR406/009/ERR4065479/ERR4065479.fastq.gz   | NA                                                                             | NA  |
| SJN002      | SJN002.A0101 | 3    | 4                | SE      | Homo sapiens | double       | none          | ftp://ftp.sra.ebi.ac.uk/vol1/fastq/ERR406/000/ERR4065480/ERR4065480.fastq.gz   | NA                                                                             | NA  |
| SJN002      | SJN002.B0101 | 3    | 4                | SE      | Homo sapiens | double       | none          | ftp://ftp.sra.ebi.ac.uk/vol1/fastq/ERR406/001/ERR4065481/ERR4065481.fastq.gz   | NA                                                                             | NA  |
| SJN003      | SJN003.A0101 | 3    | 4                | SE      | Homo sapiens | double       | none          | ftp://ftp.sra.ebi.ac.uk/vol1/fastq/ERR406/002/ERR4065482/ERR4065482.fastq.gz   | NA                                                                             | NA  |
| SJN003      | SJN003.B0101 | 3    | 4                | SE      | Homo sapiens | double       | none          | ftp://ftp.sra.ebi.ac.uk/vol1/fastq/ERR406/003/ERR4065483/ERR4065483.fastq.gz   | NA                                                                             | NA  |
| SJN001      | SJN001.A0102 | 3    | 4                | PE      | Homo sapiens | double       | half          | ftp://ftp.sra.ebi.ac.uk/vol1/fastq/ERR406/007/ERR4065497/ERR4065497_1.fastq.gz | ftp://ftp.sra.ebi.ac.uk/vol1/fastq/ERR406/007/ERR4065497/ERR4065497_2.fastq.gz | NA  |
| SJN001      | SJN001.B0102 | 3    | 4                | PE      | Homo sapiens | double       | half          | ftp://ftp.sra.ebi.ac.uk/vol1/fastq/ERR406/008/ERR4065498/ERR4065498_1.fastq.gz | ftp://ftp.sra.ebi.ac.uk/vol1/fastq/ERR406/008/ERR4065498/ERR4065498_2.fastq.gz | NA  |
| SJN002      | SJN002.A0102 | 3    | 4                | PE      | Homo sapiens | double       | half          | ftp://ftp.sra.ebi.ac.uk/vol1/fastq/ERR406/009/ERR4065499/ERR4065499_1.fastq.gz | ftp://ftp.sra.ebi.ac.uk/vol1/fastq/ERR406/009/ERR4065499/ERR4065499_2.fastq.gz | NA  |
| SJN002      | SJN002.B0102 | 3    | 4                | PE      | Homo sapiens | double       | half          | ftp://ftp.sra.ebi.ac.uk/vol1/fastq/ERR406/000/ERR4065500/ERR4065500_1.fastq.gz | ftp://ftp.sra.ebi.ac.uk/vol1/fastq/ERR406/000/ERR4065500/ERR4065500_2.fastq.gz | NA  |
| SJN003      | SJN003.A0102 | 3    | 4                | PE      | Homo sapiens | double       | half          | ftp://ftp.sra.ebi.ac.uk/vol1/fastq/ERR406/001/ERR4065501/ERR4065501_1.fastq.gz | ftp://ftp.sra.ebi.ac.uk/vol1/fastq/ERR406/001/ERR4065501/ERR4065501_2.fastq.gz | NA  |
| SJN003      | SJN003.B0102 | 3    | 4                | PE      | Homo sapiens | double       | half          | ftp://ftp.sra.ebi.ac.uk/vol1/fastq/ERR406/002/ERR4065502/ERR4065502_1.fastq.gz | ftp://ftp.sra.ebi.ac.uk/vol1/fastq/ERR406/002/ERR4065502/ERR4065502_2.fastq.gz | NA  |

For the human processing, we aim follow as close as possible to settings used in
the original paper, with the main difference being the use of Picard
MarkDuplicates instead of DeDup, due to more accurate duplicate removal when
single-end or unmerged singleton reads exist in the library. We set the mapping
parameters to that of the defaults in EAGER v1.92.55 (used by Barquera et al.).

The original publication used the HOPS pipeline for pathogen screening, however
did not report the database used ("We therefore decided to screen these
individuals for potential pathogenic agents using a bioinformatic approach [68]
to screen and filter reads from the genomic libraries), therefore we opted here
to use a broad database typically used for BLAST screening - the NCBI Nucleotide
(nt) database
[https://www.ncbi.nlm.nih.gov/nucleotide/](https://www.ncbi.nlm.nih.gov/nucleotide/).
This includes a large range of sequences from most organisms with data in the
NCBI databases. We downloaded the FASTA file on 2017-10-26 12:39 from:
ftp://ftp-trace.ncbi.nih.gov/blast/db/FASTA/nt.gz, and indexed using MALT v0.4.0
as follows:

```bash
malt-build -J-Xmx1800G \
--step 2 \
-i ../raw/full-nt_2017_10/nt.gz \
-s DNA \
-d full-nt_2017-10 \
-t 112 \
-a2taxonomy ../acc2tax/nucl_acc2tax-May2017.abin
```

We were unable to use a later NT database as once indexed these are too large to
fit in a 2 TB memory cluster node.

We will also use the same pathogen screening list (including Yersinia pestis) as
was used in the original publication and parameters, as supplied on the HOPS
repository
(https://github.com/rhuebler/HOPS/blob/external/Resources/default_list.txt,
Zenodo: https://doi.org/10.5281/zenodo.3362248), and also the same default MALT
parameters as set in the HOPS pipeline, as presumably these were used in the
HOPS screening by Barquera et al.

## Running

In addition to the settings above, we also turned on Mitochondrial to Nuclear
ratio, Sex Determination, and Nuclear Contamination modules to represent typical
quality control analyses of interest for human population-genetics, however we
know the nuclear-contamination module will likely not produce sufficient results
due to the shallow shotgun-sequenced data not covering sufficient SNPs for
estimation with ANGSD.

The following command was used, with the profile set to the custom settings used
for running nf-core/eager on MPI-SHH, using a Singularity
(v3.5.1+124-g54a90cd63+dirty) container.

```bash
nextflow run nf-core/eager -r dev \
-profile microbiome_screening,sdag,shh \
-with-tower \
--input 'barquera2020_pathogenscreening.tsv' \
--fasta 'ftp://ftp-trace.ncbi.nih.gov/1000genomes/ftp/technical/reference/phase2_reference_assembly_sequence/hs37d5.fa.gz' \
--bwaalnn 0.01 \
--bwaalnl 32 \
--run_bam_filtering \
--bam_discard_unmapped \
--bam_unmapped_type fastq \
--dedupper markduplicates \
--run_mtnucratio \
--run_nuclear_contamination \
--run_sexdeterrmine \
--sexdeterrmine_bedfile 'https://github.com/nf-core/test-datasets/raw/eager/reference/Human/1240K.pos.list_hs37d5.0based.bed.gz' \
--run_metagenomic_screening \
--metagenomic_tool malt \
--run_maltextract \
--percent_identity 90 \
--malt_top_percent 1 \
--malt_min_support_mode 'reads' \
--metagenomic_min_support_reads 1 \
--malt_max_queries 100 \
--malt_memory_mode load \
--maltextract_taxon_list 'https://raw.githubusercontent.com/rhuebler/HOPS/external/Resources/default_list.txt' \
--maltextract_filter def_anc \
--maltextract_toppercent 0.01 \
--maltextract_destackingoff \
--maltextract_downsamplingoff \
--maltextract_duplicateremovaloff \
--maltextract_matches \
--maltextract_megansummary \
--maltextract_percentidentity 90.0 \
--maltextract_topalignment \
--database 'malt/databases/indexed/index040/full-nt_2017-10/' \
--maltextract_ncbifiles 'resources/'
```

The MultiQC report was then loaded from `results/multiqc/multiqc_report.html`
(see file `barquera2020_sdag_multiqc_1_9_report.html` in this directory) and
the HOPS post-processing script's heatmap (`heatmap_overview_Wevid.pdf` in this
directory) to check the Human and Pathogen screening
results respectively.

## Post-Processing

The HOPS summary heatmap has been additionally integrated in the upcoming next
version of MultiQC, and we wished to demonstrate how all results for both
aspects of typical aDNA screening is now integrated into a single report.

To do this we installed latest stable version of MultiQC with conda

```bash
conda create -n multiqc bioconda::multiqc
```

and then updated this installation to the v1.10dev version

```bash
pip install --upgrade --force-reinstall git+https://github.com/ewels/MultiQC.git
```

To run this version of MultiQC, we navigated to the `results/` directory of the
nf-core/eager run, copied the nf-core/eager  MultiQC config file  (v2.2.0dev
version) from the pulled Nextflow repository, and then re-ran MultiQC.

```bash
cd results/
cp ~/.nextflow/assets/nf-core/eager/assets/multiqc_config.yaml .

## Run MultiQC
multiqc . -c multiqc_config.yaml -n multiqc1_10.html -o multiqc1_10
```

The resulting file which is a replicate of the original MultiQC 1.9 report
but with the additional inclusion of the HOPS heatmap can be seen in the file
`barquera2020_sdag_multiqc_1_10_report.html` in this directory.
