## Supplementary figures and images for "Reproducible, portable, and efficient ancient genome reconstruction with nf-core/eager"

### heatmap_overview_Wevid.pdf

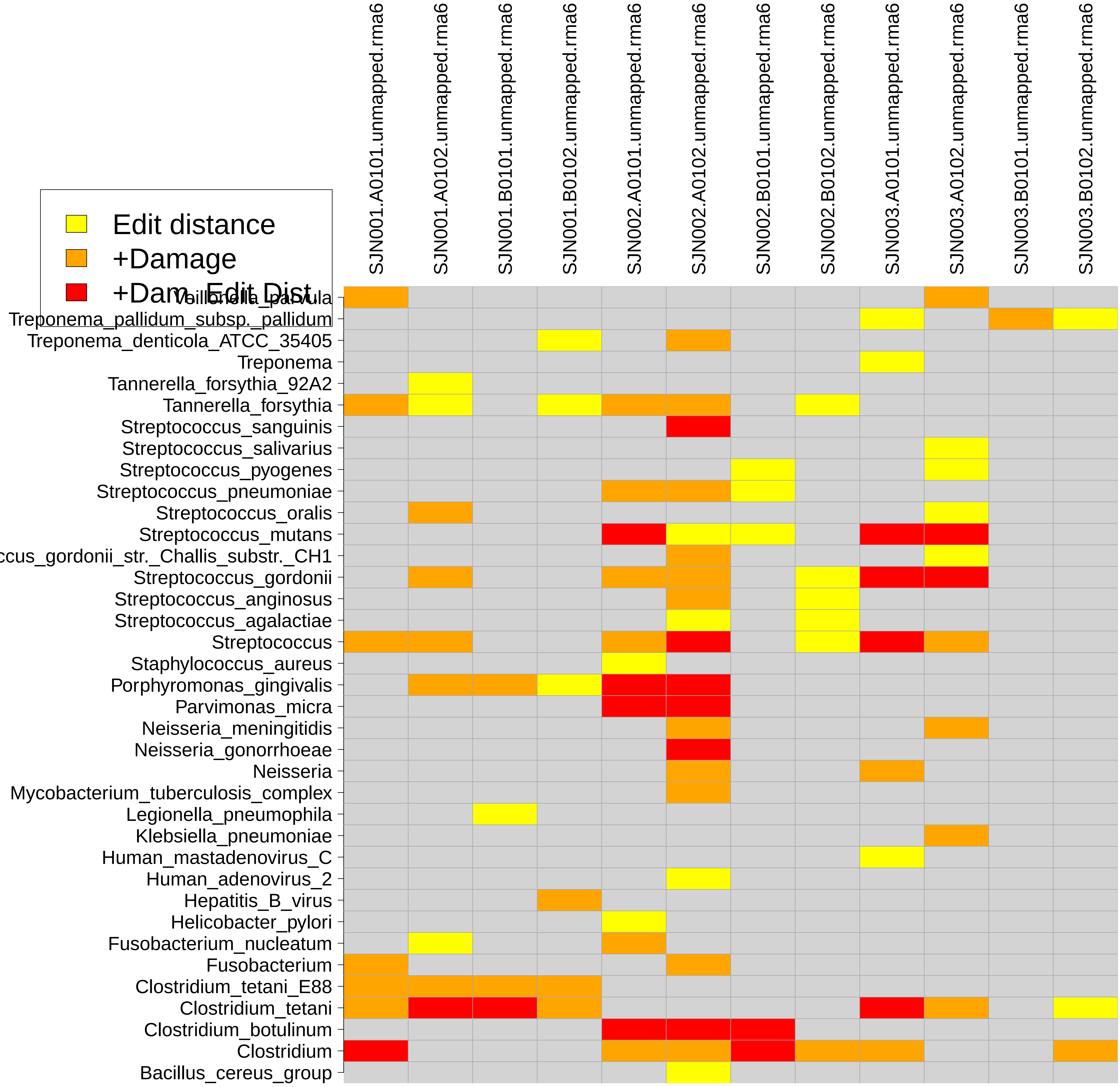
